# Quantification of disease-associated RNA tandem repeats by nanopore sensing

**DOI:** 10.1101/2025.05.15.653973

**Authors:** G. Patiño-Guillén, J. Pešović, M. Panić, M. Earle, A. Ninković, S. Petrușca, D. Savić-Pavićević, U.F. Keyser, F. Bošković

## Abstract

Short tandem repeat expansions underlie a class of neurological and neuromuscular diseases known as repeat expansion disorders (REDs), yet the precise characterization of these repeats remains technically challenging. Conventional amplification-based methods fail to resolve repeat length accurately due to amplification bias and sequence homogeneity. Here, we present a single-molecule nanopore-based strategy that enables direct quantification of tandem repeats in native RNA. By assembling RNA:DNA nanostructures that encode specific repeat number, we achieve repeat size discrimination with a resolution of 18 nucleotides. Using tandem repeat-containing RNA, we successfully detect and discriminate disease-relevant repeat lengths associated with myotonic dystrophy types 1 (DM1) and 2 (DM2), and congenital central hypoventilation syndrome-1 (CCHS1). Finally, we apply our method to total RNA extracted from a DM1 human cell line model, demonstrating its compatibility with complex biological samples. Our approach offers a platform for studying repeat expansion biology at the single-molecule level, with broad implications for diagnostics, clinical research and multiplexed repeat profiling.

## Main Text

Short tandem repeats are DNA sequences repeated consecutively with lengths of repeated motifs^1^. Expansions of short tandem repeats are implicated in over 60 monogenic disorders, of which roughly half occur within protein-coding regions of genes, and are collectively known as repeat expansion disorders (REDs)^2–5^. Among them, polyglutamine and polyalanine diseases, such as congenital central hypoventilation syndrome-1 (CCHS1), are caused by relatively small repeat expansions (less than a hundred) in the coding region of the gene. In addition, diseases caused by repeat expansions in non-coding regions of gene, include myotonic dystrophy type 1 (DM1) and type 2 (DM2), fragile X syndrome, and amyotrophic lateral sclerosis^3,6^. The severity and onset of REDs typically depend on the number of repeats and interruptions, i.e. expansion size, making their accurate quantification critical^6^. In addition, repeat expansions are transcribed in RNA which plays a central role in the molecular pathogenesis of the majority of REDs and represents the major target for the developing gene modifying therapies^7,8^.

Depending on the expansion size, current methods for quantifying tandem repeats rely on PCR-based amplification combined with fragment analysis, and/or Illumina sequencing for small expansions^9,10^, and southern blot and Small-pool PCR (SP-PCR) for large expansions^11,12^. Nevertheless, DNA polymerase slippage during PCR amplification of tandem repeats^13^, a phenomenon extensively studied in trinucleotide repeats^14^, introduces biases that can prevent precise assessment of repeat expansion sizes. Amplification biases also represent a challenge for sequencing, as preferential amplification of wild-type alleles may occur^15,16^. Since the expanded repeat regions often exceed the short read lengths, these methods struggle with sequencing through the full expansion, leading to incomplete or ambiguous results^17,18^. Recently, long-read nanopore sequencing has been implemented for amplification-free studying of tandem repeat expansions^19–21^. Nevertheless, error rates per-base are generally higher than short-read sequencing technologies, with particularly pronounced substitution and indel errors in homopolymer regions^4,22^. Moreover, the lack of specialized base calling algorithms^23^, the sequence complexity^4^, and the amounts of material needed for experiments prevents their immediate use for diagnostics purposes. Similarly, PacBio sequencing has been used to characterize DNA tandem repeats at whole genome scale and targeting specific loci *via* PacBio PureTarget^24,25^. PacBio offers long reads and high consensus accuracy though HiFi reads^24^. However, it also demands for high DNA inputs^26^, and high error rates prevail for repetitive or GC-rich motifs which demand specialized algorithms for data processing^27,28^. Due to these challenges, there is still a clear need for more precise and amplification-free methods to quantify tandem repeat expansions.

Here, we introduce an amplification-free tool for determination of tandem repeat expansions that relies on RNA:DNA nanotechnology and solid-state nanopores^29–34^. Our method addresses current unresolved challenges related to reverse transcription and amplification biases that limit accurate assessment of repeat sizes in expanded RNA molecules. We differentiate pathogenic and non-pathogenic sizes of tandem repeat arrays at the single-molecule level by careful design of RNA:DNA hybrid nanostructures from RNA transcripts containing clinically relevant tandem repeats. RNA:DNA nanostructures are passed through a nanopore for quantification of the tandem repeats. The method is suitable for characterization of transcribed small tandem repeats. We characterize RNA tandem repeats arrays of varying sizes derived from the loci associated with DM1, DM2, and CCHS1, showing a resolution of 18 nucleotides. Our resolution allows the distinction of non-pathogenic and pathogenic tandem repeat sizes. Finally, we study tandem repeats in total RNA extracted from a DM1 human cell line model, performing direct determination of expansions in complex biological backgrounds. Our method offers a platform for studying repeat expansion biology at transcript level, and with broad implications for RED diagnostics.

### Nanostructure-mediated nanopore sensing of RNA tandem repeats associated with myotonic dystrophy type 1 and type 2

We characterize tandem repeats in RNA using nanopore sensing through the assembly of RNA:DNA nanostructures (**Figure 1a**). First, we cloned DNA tandem repeats into a plasmid from human loci. Then, we transcribed the DNA plasmid *in vitro* to produce RNA containing tandem repeats^35^ (**Figure S1, S2, S3**). The sequences of the DNA constructs used for RNA synthesis are detailed in **Tables S1** and **S2**. We positioned tandem repeats towards the 5’ end. *In vitro* transcription yields full-length and prematurely terminated transcripts^35^. In this study, we use the full-length RNA to characterize tandem repeats. The RNA is hybridized with complementary single-stranded DNA oligonucleotides (40 nt) to produce an RNA:DNA nanostructure. By hybridization, we introduced structural labels, such as monovalent streptavidin^36^, which enable mapping of the RNA and identification of tandem repeats *via* nanopore sensing.

**Figure 1.**
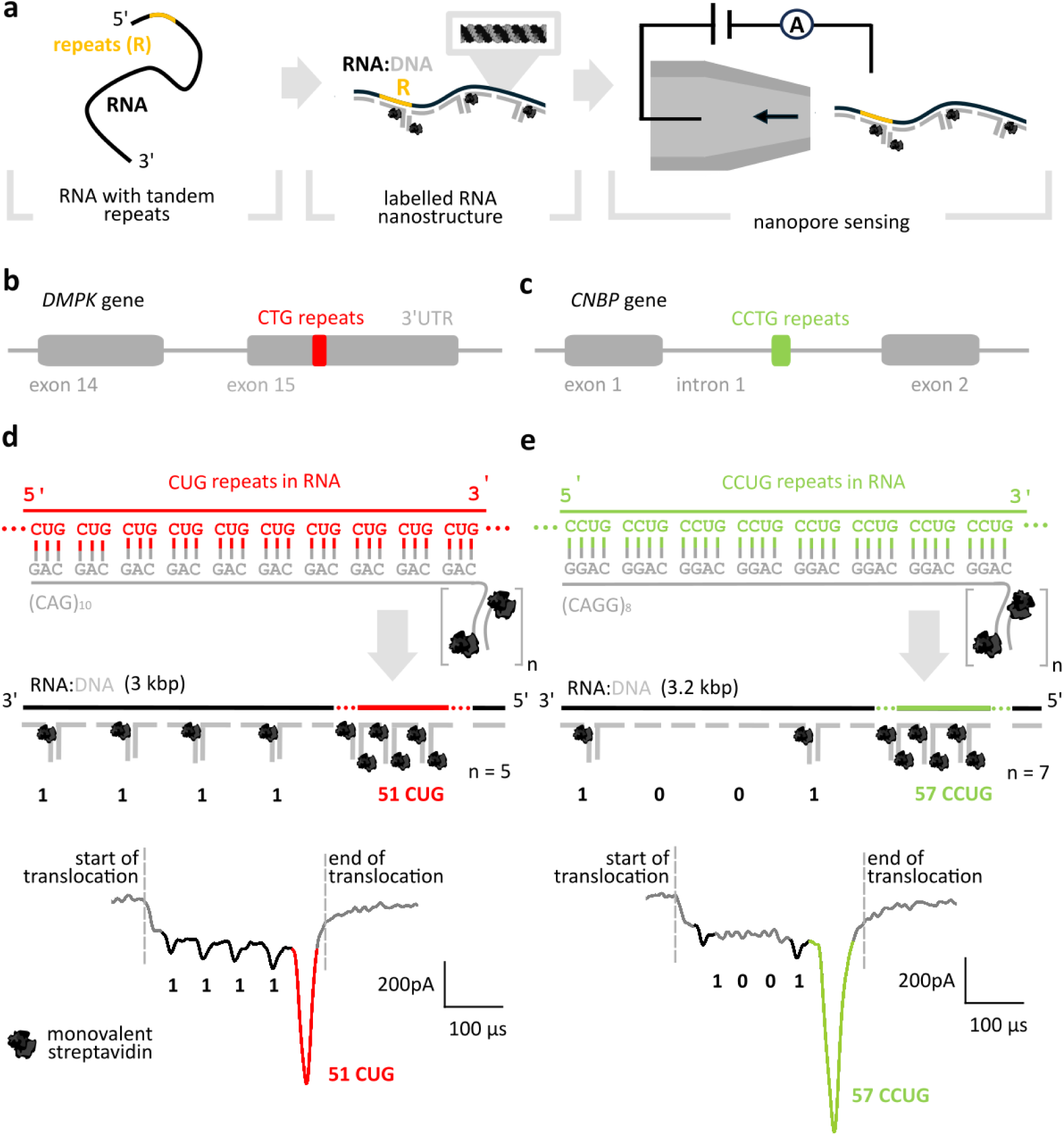
Identification of CUG and CCUG tandem repeats using RNA:DNA nanotechnology and nanopore sensing. **a** We *in vitro* transcribed ∼3 kb RNA containing CUG or CCUG repeats from human-derived DNA plasmids. The resulting RNA is hybridized with short oligonucleotides (∼40 nt) to produce an RNA:DNA nanostructure. Then, we translocated the nanostructure through a nanopore facilitating single-molecule characterization of tandem repeats. **b** CTG tandem repeat expansions (CUG in RNA) in *DMPK* gene, causing DM1. **c** CCTG tandem repeat expansions (CCUG in RNA) in the *CNBP* gene, causing DM2. **d** 3 kb RNA with 51 CUG repeats is hybridized with oligonucleotides to produce an RNA:DNA nanostructure. The nanostructure includes a barcode region and a repeat-sensing region. The barcode comprises a series of monovalent streptavidin distributed throughout the sequence, each representing a ‘1’; and their absence indicating a ‘0’. We assigned this nanostructure with a ‘1111’ barcode. The repeats are labelled using ‘n’ (CAG)_10_ oligonucleotides that bind two streptavidins each. Translocation of the nanostructure through a solid-state nanopore produces a drop in ionic current ascribed to the nanostructure’s backbone (gray), four downward current spikes are produced from the ‘1111’ barcode (black), and a prominent current spike (red) is produced from the five (CAG)_10_ oligonucleotides bound to the 51 CUG repeats. **e** We assigned for CCUG-containing nanostructures (3.2 kbp) a ‘1001’ barcode, that yields two downward current spikes in the nanopore readout (black). CCUG repeats detection is achieved using seven (CAGG)_8_ oligonucleotides, which produce a prominent current spike (green).

Initially, we detected trinucleotide and tetranucleotide repeats causing DM1 and DM2. Expansion of CTG trinucleotide DNA tandem repeats (transcribed as CUG repeats in RNA) in the 3′ untranslated region (3′UTR) of *DMPK* (myotonic dystrophy protein kinase) gene, is responsible for DM1^37^ (**Figure 1b**). The number of CTG repeats varies from 5 to 34 in wild-type alleles, while the pathogenic alleles have more than 50 repeats ^38^. Similarly, DM2 is caused by the expansion of CCTG tetranucleotide tandem repeats (transcribed as CCUG in RNA) and are found in intron 1 of the *CNBP* (CCHC-Type Zinc Finger Nucleic Acid Binding Protein) gene.^39^ For DM2, the wild-type tandem repeat array contains fewer than 26 repeats; alleles with 26 to 75 repeats correspond to a premutation. A repeat length of over 75 defines the pathogenic cutoff. The term “expansion” collectively refers to both the premutation and pathogenic categories (**Figure 1c**)^40,41^.

We assembled an RNA:DNA nanostructure from a ∼3 kb transcript containing 51 CUG repeats, for a proof-of-concept detection of trinucleotide tandem repeat expansions (**Figure 1d**). The RNA:DNA nanostructure is designed with a barcode and a repeat-sensing region. The barcode allows for specific identification of the transcript. In this region, the presence of an attached streptavidin denotes a ‘1’, while its absence signifies a ‘0’. The nanostructure was engineered to have a ‘1111’ barcode. In the repeat-sensing region, CUG repeats are identified though hybridization with a number ‘n’ of complementary (CAG)_10_ oligonucleotides, each containing an overhang for streptavidin binding and a complementary DNA strand, which binds an additional streptavidin. While it is possible for different numbers of (CAG)_10_ oligonucleotides to hybridize to the repeat array, the arrangement where the number of oligonucleotides maximizes base pairing provides the lowest free energy and greatest stability^42^, as demonstrated *via* agarose gel electrophoresis (**Figure S4**). Therefore, the binding of five (CAG)₁₀ oligonucleotides to 51 CUG repeats correspond to the most favourable assembly conformation. The detailed nanostructure design is presented in **Figure S5a**, **S6a** and **Table S3**.

We then ‘read’ the nanostructures *via* nanopore sensing. Upon applying a voltage bias, the negatively charged RNA:DNA nanostructure translocated through the nanopore towards the positively charged electrode^43,44^. As each nanostructure passes through, it partially obstructs the pore, depleting ion flow, resulting in a transient drop in the ionic current^45,46^. Every time a structural label (i.e. streptavidin) passes through the pore, either from the barcode or the repeat-sensing region, a larger number of ions are prevented from passing through the nanopore producing a deeper drop in the ionic current^30,47^.

The nanostructures with CUG repeats translocate through the nanopore (**Figure 1d**) inducing an initial transient drop in ionic current, that corresponds to the RNA:DNA duplex backbone (gray). Each streptavidin in the barcode region then generates a distinct downward current spike (black), that together represent the ‘1111’ barcode of the nanostructure. Finally, a prominent current spike (red) arises from the consecutive streptavidins bound to the repeats through five hybridized (CAG)_10_ oligonucleotides. The depth of this prominent spike is proportional to the number of streptavidins that attached to the repeats^30^ (translocation events with labelled and unlabelled repeats are presented in **Figure S7** and **Figure S8**, respectively). To ensure that our repeats are labelled as described, we performed electrophoretic mobility shift assay (**Figure S4**).

Likewise, we perform nanopore sensing of CCUG tetranucleotide repeats (**Figure 1e**). We assemble an RNA:DNA nanostructure with a ‘1001’ barcode and seven (CAGG)_8_ oligonucleotides with bound streptavidin for sensing the tandem repeats (the nanostructure design is detailed in **Figure S5b** and **Table S4**). As with the previous construct, the backbone of the nanostructure induces a drop in current (gray) and each ‘1’ in the barcode produces a downward current spike (black). Streptavidin labelling of the CCUG repeats in the RNA produces a spike (green) that signals the tandem repeats in the transcript (additional translocation events are presented in **Figure S9**). Interruptions or mismatches in CCUG repeat arrays, such as the inclusion of GCTG or TCTG motifs, do not prevent the binding of (CAGG)_8_ oligonucleotides and are not expected to cause significant variation in the ionic current readout with the present labelling strategy.

We demonstrated detection of trinucleotide and tetranucleotide RNA tandem repeat expansions using nanopore readout of our nanostructures. The RNA:DNA nanostructure design facilitates tandem repeat positioning on the individual transcripts.

### Quantification of RNA-containing CUG repeats discriminates non-pathogenic and pathogenic repeat sizes in DM1

Determination of tandem repeat sizes is essential for accurate RED diagnosis^6^. We quantify tandem repeats on RNA produced from DNA plasmids holding 12, 24 and 30 CUG repeats (wild-type sizes), and 51 CUG repeats (pathogenic size), showcasing our counting resolution. The repeat size distribution for each DNA plasmid used to synthetize these transcripts was confirmed through Sanger sequencing and fragment analysis as shown in **Figure S10**.

We targeted the CUG repeats using (CAG)_10_ oligonucleotides that bind up to two streptavidins. Larger CUG repeat arrays facilitate the attachment of more (CAG)_10_ oligonucleotides with streptavidin, which are then detected with nanopores as current blockages (**Figure 2a**).

**Figure 2.**
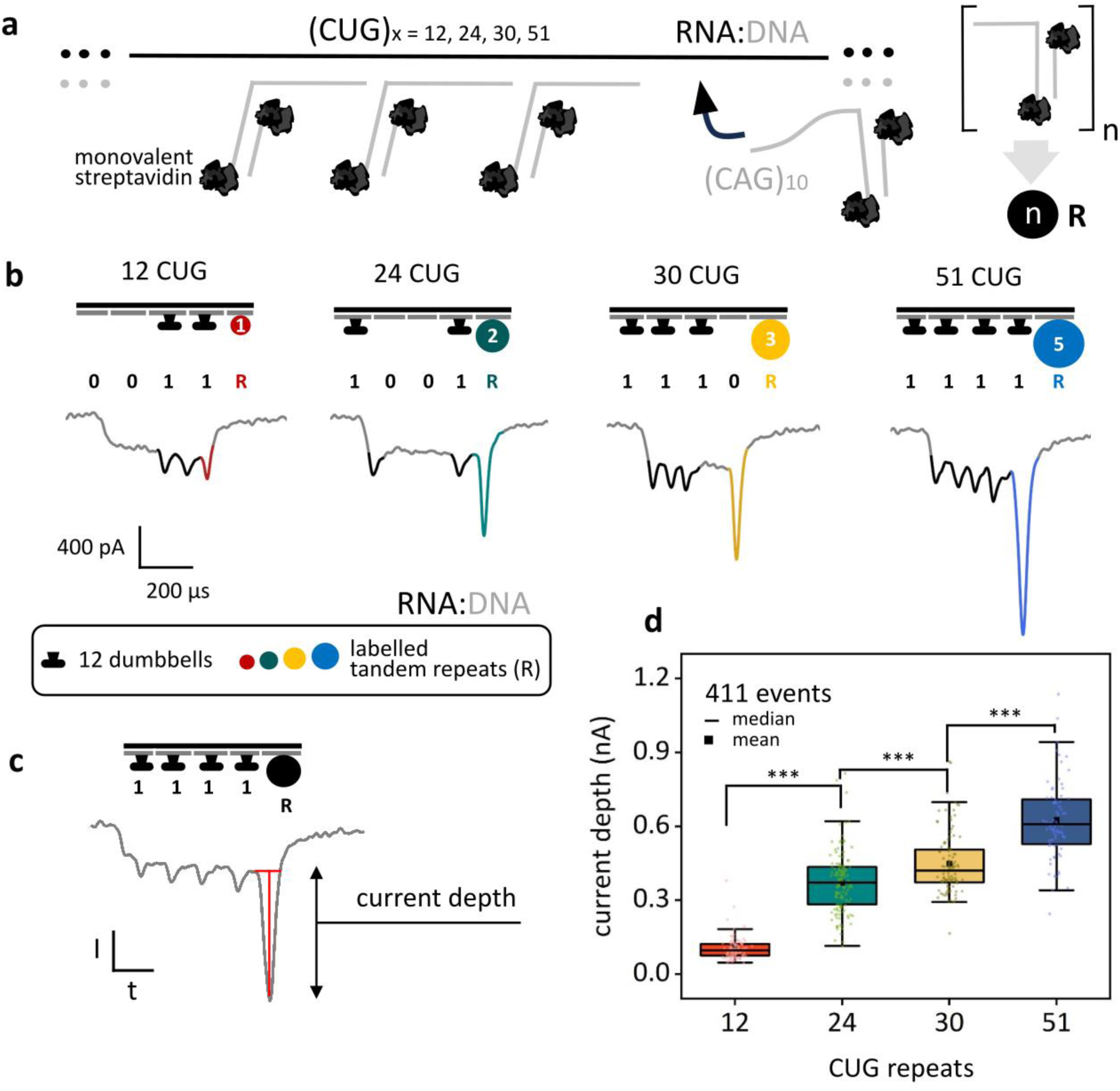
RNA:DNA nanostructures enable quantification of CUG tandem repeats *via* nanopore sensing. **a** We investigate various CUG tandem repeats lengths in RNA, including wild-type sizes – 12, 24, and 30 repeats, and a pathogenic size – 51 repeats, by assembling RNA:DNA nanostructures and translocating them through nanopores. Each repeat array hybridizes with ‘n’ complementary (CAG)₁₀ oligonucleotides, allowing two streptavidins to bind per oligonucleotide. Larger repeat arrays accommodate more (CAG)₁₀ oligonucleotides on the nanostructures, facilitating quantification of the repeats as the nanostructures translocate through a nanopore. **b** We assembled RNA:DNA nanostructures with distinct barcodes corresponding to specific CUG repeat sizes and translocated them through the same nanopore. RNA with 12 CUG repeats was barcoded as ‘0011’, 24 repeats as ‘1001’, 30 repeats as ‘1110’, and 51 repeats as ‘1111’ where ‘1’ corresponds to 12 consecutive DNA dumbbells. A greater number ‘n’ of consecutive oligonucleotides with streptavidins can bind to larger repeat array sizes, resulting in increased ion depletion during the translocation of the nanostructure through a nanopore, thus generating a more prominent current spike in the nanopore readout. **c** We measure the depth of the current spike caused by streptavidin binding to the repeats, to quantify tandem repeats. **d** We observed an increase in the median current spike depth as the number of repeats increases, with all populations showing mean independence, Welch’s t-test ***p-values < 0.001: 4.9×10^-57^ (12 and 24), 2.4×10^-6^ (24 and 30), 6.2 ×10^-13^ (30 and 51).

We assembled the RNA:DNA nanostructures with distinct barcodes to differentiate transcripts containing different repeat numbers for multiplexed characterization. Specifically, RNA:DNA nanostructures with 12 CUG repeats were tagged with the barcode ‘0011’, those with 24 repeats with ‘1001’, 30 repeats with ‘0111’, and 51 repeats with ‘1111’ (‘1’ corresponds to 12 consecutive DNA dumbbells). The nanostructures were translocated through the same solid-state nanopore, allowing for direct comparison of the ionic current readout across the four nanostructures while eliminating pore-dependent variability^48^ (**Figure 2b**). Detailed descriptions of the nanostructures are included in **Figure S11** and **Tables S5** to **S8**.

Upon translocation of the nanostructures through the nanopore, downward current spikes (black) reveal the barcode ascribed to each construct, while an additional current spike is originated from the consecutive streptavidin tagging the repeats. Repeat arrays of larger sizes can bind more streptavidins, leading to increased ion depletion as the nanostructure translocates through the nanopore. The depth of this current spike is dependent on the number of streptavidins that bind the repeats. We use the depth of this current spike to quantitatively assess the size of tandem repeats in each RNA (**Figure 2c**). We include additional translocation events for each RNA construct in **Figures S12**, and relative positioning of features on RNA:DNA nanostructure in **Figure S13**.

The depth of the current spike of the repeats increases with the number of CUG repeats in the RNA (**Figure 2d**). We observe median current spikes of 0.09 nA, 0.37 nA, 0.42 nA, and 0.61 nA, for 12, 24, 30 and 51 CUG repeats, respectively. A one-way ANOVA test f-ratio value is 213.9 and the p-value is < 0.00001. The p-value of t-test with unequal variance (Welch’s test) was 4.9×10^-57^ for the comparison between 12 and 24 CUG repeats, 2.4×10^-6^ between 24 and 30 CUG repeats and 6.2 ×10^-13^ between 30 and 51 CUG repeats. The experiments were done in triplicate, an additional dataset is presented in **Figure S14**. The tandem repeat size distributions agree with the spread observed for the repeat sizes of the initial DNA plasmids used for RNA synthesis. In **Figure S15**, we have included visual representation of the allele sizes from the ionic current readout. We also show the characterization of a repeat array with significant size heterogeneity typically seen in patient cells due to repeat instability in **Figure S16**. Beyond spike depth, spike charge deficit also proves effective for repeat array size differentiation (**Figure S17**). The current spike depth may approach saturation for larger repeat arrays. However, the integrated signal, which represents the spike charge deficit, continues to increase even when the spike depth plateaus^49^. As a result, reliable discrimination of larger repeat arrays remains possible by analysing the spike area. Moreover, RNA:DNA nanostructures can translocate through the nanopore in both 5′-3′ and 3′-5′ orientations. Notably, characterization of the current spikes and discrimination of repeat array sizes is possible in both translocation directions (**Figure S18**). As shown in **Figure S19**, the length of the oligonucleotides used for labelling the repeats can also be adjusted to different sizes of repeat arrays to enhance discrimination precision.

By measuring the spike depth associated with the repeat-sensing region of the RNA:DNA nanostructures, we successfully quantify the repeat arrays in the original RNA transcript. This approach provides an accurate method for discriminating a pathogenic size of 51 repeats from non-pathogenic sizes of 12, 24 and 30 repeats, relying on the hybridization of DNA oligonucleotides to RNA without introducing amplification or short-read assembly biases. The achieved repeat resolution of 18 nucleotides (six trinucleotide repeats), is sufficient to clearly discriminate wild-type alleles and DM1 expanded alleles that are within a lower boundary of the pathogenic size.

### Six-repeat resolution in RNA sensing discriminates pathogenic alleles in CCHS1

We determine GCN tandem repeat expansion as a model where six repeat resolution is required. GCN repeats occur in *PHOX2B* gene and their expansion causes congenital central hypoventilation syndrome-1 (CCHS1). GCN repeats, where N can be adenine, guanine, cytosine or thymine (uracil in RNA) code for polyalanine tracts. There are two short polyalanine tracts within exon 3 of the *PHOX2B* gene. The second polyalanine tract most frequently has 20 GCN repeats^50^. However, an expansion in this region can lead to CCHS1, a life-threatening disease primarily characterized by an impaired respiratory control^51,52^. Alleles causing neonatal onset form have 26 to 33 GCN repeats^53^ (**Figure 3a**).

**Figure 3.**
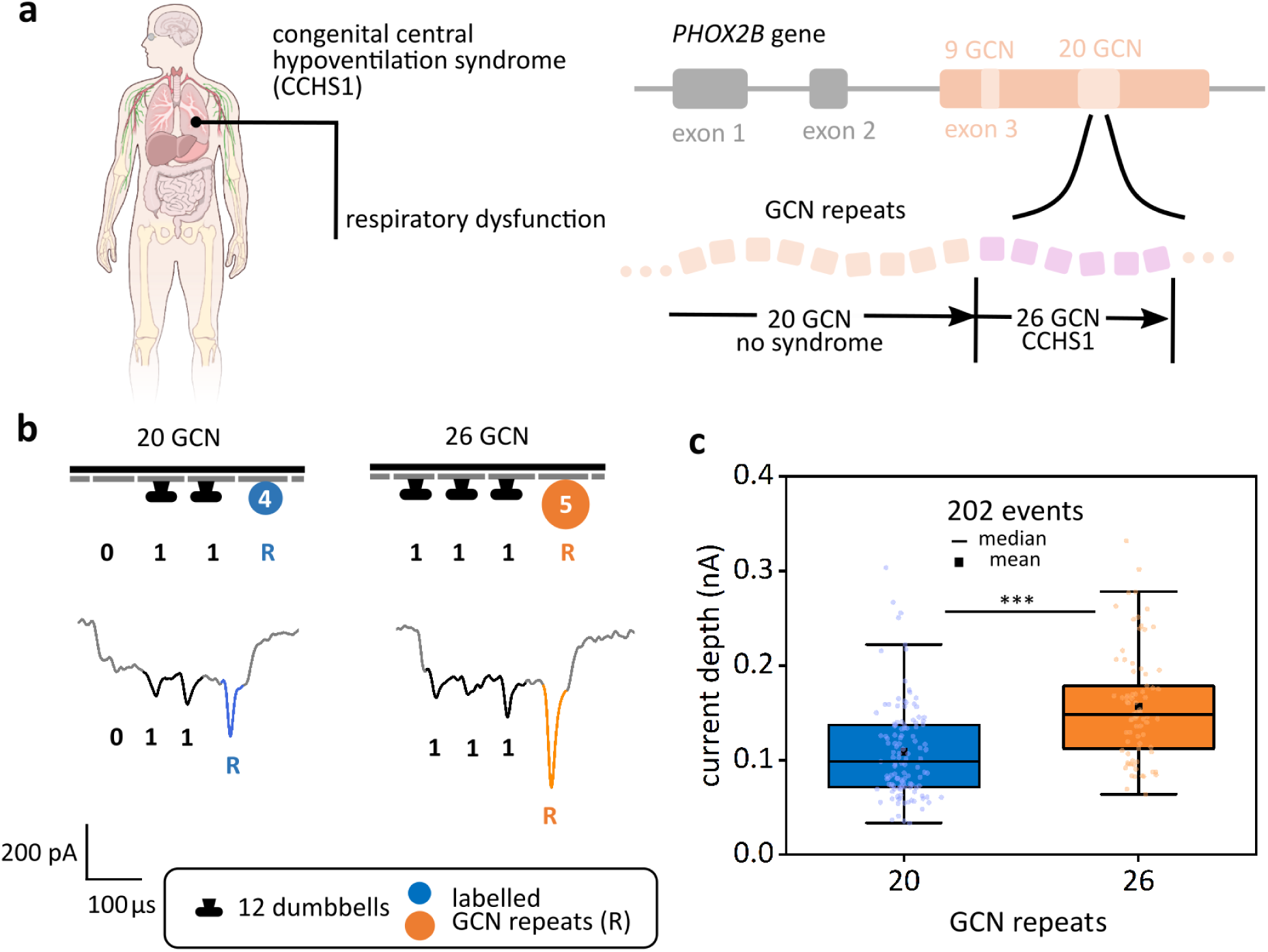
Quantitative characterization of repeats associated with CCHS1. **a** Expansion of the second, polyalanine coding GCN triplets in exon 3 of the *PHOX2B* gene causes CCHS1. **b** We produced RNA:DNA nanostructures with a ‘011’ dumbbell barcode from transcripts containing 20 GCN repeats, and RNA:DNA nanostructure with a ‘111’ barcode targeting 26 GCN repeats. The six additional repeats of the ‘111’ barcoded RNA allow for an extra (NGC)_6_ DNA oligonucleotide and streptavidin to bind to the construct, producing a larger drop in the ionic current upon translocation through a nanopore, compared to the construct with only 20 GCN repeats. **c** The current spike depth associated to the repeats labelling shows a higher median for constructs with 26 GCN repeats (0.8 and 0.15 nA, respectively), and a p-value between the distributions of 9.3×10^-9^ for Welch’s t-test. This clear distinction between the two populations enables precise differentiation of pathogenic expansion sizes, holding clinical relevance of this method for detection of CCHS1.

We produced two different DNA constructs with GCN repeats for *in vitro* transcription aided by a T7 RNA polymerase promoter (**Tables S9** and S**10**). The first construct corresponds to the wild-type allele size, with 20 GCN repeats, while the second one has 26 GCN repeats, representing a pathogenic allele size. *In vitro* transcription yielded RNA with GCN repeats of the corresponding repeat array sizes (**Figure S20**). The distribution of the repeat number in the DNA constructs was verified through Sanger sequencing and fragment analysis (**Figure S21**).

We assembled RNA:DNA nanostructures from the GCN-containing transcripts and the GCN repeats were targeted with complementary NGC DNA oligonucleotides that bound streptavidin. The details for nanostructure assembly are presented in **Tables S11** and **S12** and **Figure S22**. For 20 GCN repeats, the nanostructures were produced with a ‘011’ barcode and they bound four (NGC)_5_ oligonucleotides (**Figure 3b**). Nanopore characterization revealed a current signature with the two downward current spikes of the barcode (black) and a current spike ascribed to the labelling of the repeats (blue). For 26 GCN repeats, we engineered the nanostructure with a ‘111’ barcode and held four (NGC)_5_ oligonucleotides and an additional (NGC)_6_ oligonucleotide (**Figure 3b**). The nanopore readout showed a modulation in current with three downward current spikes, attributed to the barcode (black) and a spike associated to the streptavidin attached to the GCN repeats (orange). Additional nanopore events are presented in **Figure S23.**

The spike depth was compared for both GCN repeat arrays (**Figure 3c**), showing a greater spike depth for 26 GCN repeats than for 20 GCN repeats, with median values of 0.8 nA and 0.15 nA, respectively. The p-value of 9.3×10^-9^ indicated high fidelity of the method. Additional comparisons of spike area (**Figure S24**) and translocation directionality (**Figure S25**) supported these findings. The six trinucleotide repeat resolution (18 nucleotides), which discriminates non-pathogenic GCN repeat array size from disease-causing GCN repeat expansion size demonstrates the clinical relevance of our RNA characterization method for diagnosing CCHS1 directly from RNA.

### Identification of CUG tandem repeats in total RNA from DM1 cell line model

To validate the adaptability of our tandem repeat characterization method, we proceed to study tandem repeats in human total RNA. For this task, we constructed a DM1 model using HEK293T cells transfected with a EGFP-based expression plasmid carrying CTG tandem repeats inserted into the 3’ UTR (**Figure 4a**; plasmid construct sequences provided in **Table S13**), while the EGFP vector without the CTG repeats is used as a control. We confirmed that our cell lines contain corresponding expression plasmid by detecting EGFP expression in cells using fluorescence microscopy (**Figure 4b**) and brightfield microscopy (**Figure S26**). Both the samples transfected with the EGFP vector with and without CTG repeats in the 3’ UTR showed similar levels of EGFP fluorescence and therefore similar levels of EGFP expression. We extracted total mRNA from the cells, and the concentration of the target RNA was determined using qPCR (**Figure S27**), which indicated a concentration of 2.1 nM of target mRNA (∼1.2 kb) in total RNA.

**Figure 4.**
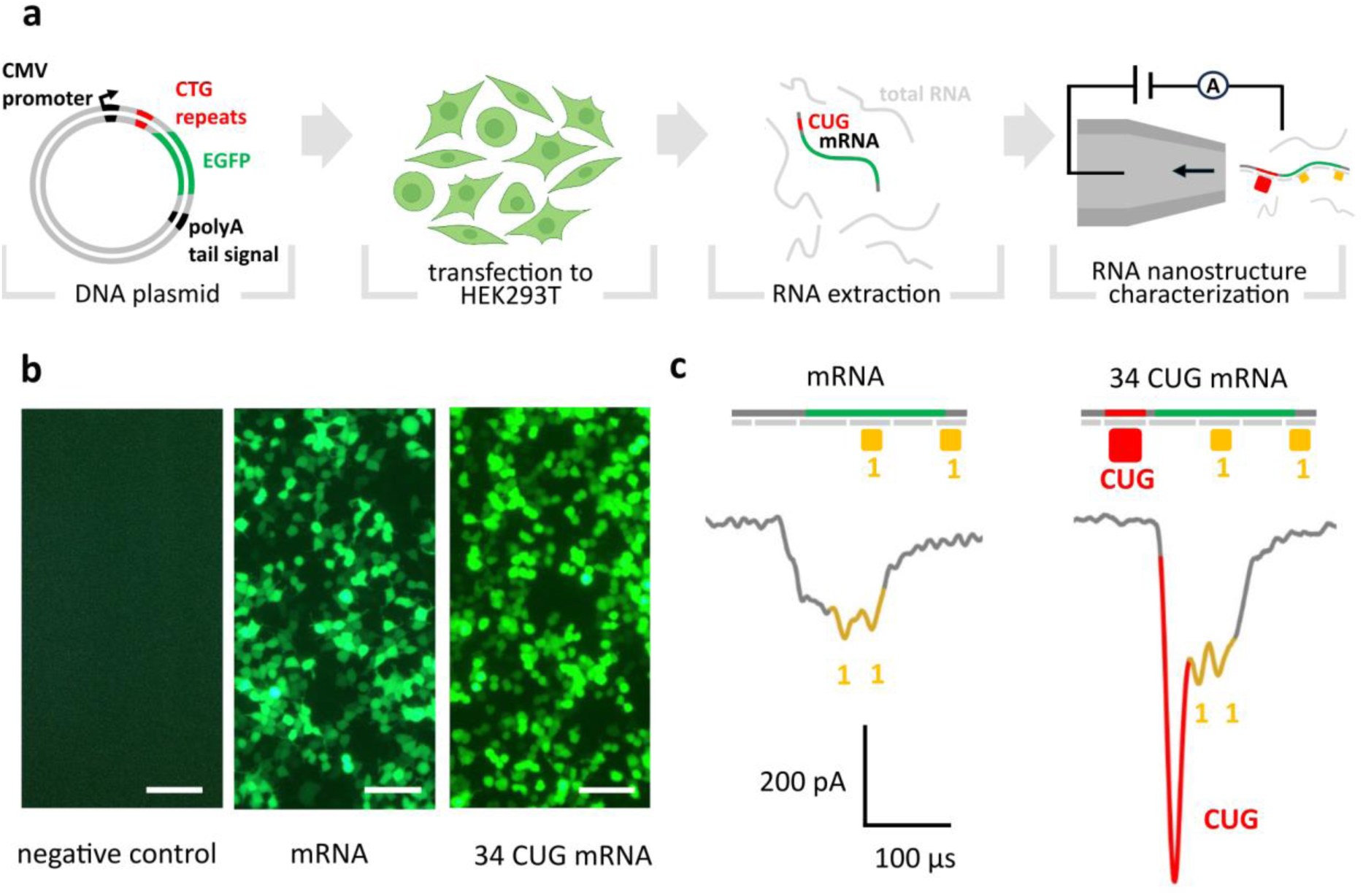
Characterization of CUG tandem repeats in human total RNA *via* nanopore sensing. **a** An overview of the transient transfection experiment. A GFP-based expression plasmid carrying CTG tandem repeats inserted into the 3’ untranslated region (UTR) was used to transiently transfect HEK293T cells. Following transfection, cells were lysed, and total RNA was extracted. The mRNA resulting from transfection (∼1.2 kb), was targeted with DNA oligonucleotides and labels to produce RNA:DNA nanostructures, which were subsequently profiled using nanopore sensing. **b** Expression of EGFP upon transient transfection was confirmed using fluorescence microscopy and therefore the presence of the mRNA of interest in the lysates HEK293T cells. Fluorescence characterization was performed for transfections incorporating the CTG repeats into the DNA construct, and unmodified DNA construct (without the repeats) as control. Scale bars are 500 µm. **c** RNA:DNA nanostructures were assembled for unmodified and repeat-containing mRNA, each tagged with a ‘11’ barcode. Nanopore mearuements show two current spikes associated to the barcode (gold). For samples with CUG repeats, a third downward current spike (red) is observed, ascribed to labelling of CUG repeats with streptavidin.

Next, we mixed total RNA (**Figure S28**) containing mRNA with CUG tandem repeats, with DNA oligonucleotides (sequences are provided in **Table S14**) that targeted the mRNA to produce an RNA:DNA nanostructure which was subsequently labelled with streptavidin and detected *via* nanopore sensing. We only assemble RNA:DNA nanostructures from the mRNA and do not target the remaining total RNA background. To ensure the robustness of our method for low mRNA concentrations, we optimized the assembly protocol for generating RNA:DNA nanostructures using MS2 RNA^54^ (**Figures S29** and **S30**, **Table S15**) and in the presence of total RNA background (**Figure S31**).

Lastly, we performed nanopore measurements with RNA:DNA nanostructures assembled in total RNA (**Figure 4c**). The construct lacking tandem repeats, tagged with a ‘11’ barcode, produces a current trace with two downward current spikes associated to the barcode (gold) and a plateau region ascribed to the rest of the construct’s backbone (grey). Similarly, the current modulations ascribed to the RNA:DNA nanostructure containing CUG repeats, also tagged with a ‘11’ barcode, present the two barcode associated spikes, with an additional current spike arising from the streptavidin bound to the CUG repeats (**Figure 4c**), showing the capability of our method to characterize tandem repeats from a complex mixture of total RNA. Detailed designs of the nanostructures are presented in **Figure S32**. To further validate the selectivity and specificity of the method, we also performed nanopore measurements of MS2 RNA nanostructures in total RNA background (**Figure S33**, **S34, Table S16**). Moreover, we estimated the translocation frequency of MS2 RNA nanostructures and mRNA nanostructures with and without repeats, which further validated the specificity of our assay (**Figure S35**). Translocations of unlabelled total RNA generate non-specific ionic current drops that are readily distinguishable from the highly specific RNA:DNA nanostructure nanopore readout^34^, which we exclude from the analysis.

RNA:DNA nanostructure assembly and subsequent characterization *via* nanopores is not limited to the detection of overexpressed or highly abundant RNA. Characterization of mRNA with low copy numbers has been demonstrated previously^30,44^. Alternatively, the nanostructure capture rate and throughput can be increased through simultaneous ionic current measurements using multiple nanopores^55^. Parallelization of nanopore measurements does not necessarily require a proportional increase in equipment cost, as multiple nanopore acquisitions can be performed in parallel using a single-amplifier architecture^55^. In addition, the barcode labelling features remain resolvable at moderate sampling rates (∼200 kHz)^56^.

These results highlight the capability of our method to assemble and detect RNA:DNA nanostructures derived from transcripts containing tandem repeats from human cell line models. Our approach exhibits high selectivity and effectively enables single-molecule detection of low-abundance transcripts in the presence of abundant RNA background.

## Conclusions

In this work, we demonstrate the use of nanopore sensing and RNA:DNA nanotechnology to identify and quantify clinically relevant tandem repeats directly on RNA molecules produced *in vitro* and *in vivo*. Our conclusions extend to *ex situ* detection and discrimination of RNA targets with different repeat array lengths. By sensing RNA:DNA nanostructures derived from CUG, CCUG and GCN repeat-containing transcripts, we identify variable-length repeat arrays in kilobase-range RNA molecules. Additionally, we map their relative position within the RNA molecules while detecting other transcript elements, such as the introduced barcodes that allow for multiplexed characterization. We characterized CUG and CCUG repeats as reliable validation models for quantification of tandem repeat arrays with variable motif lengths. We studied GCN repeats for validation of our model for non-uniform arrays and to demonstrate its resolution holds clinical relevance. Our approach may also be applicable to other STRs with 100% (e.g. C9orf72) and low GC content (e.g. FAME associated loci), and to variable number tandem repeats, provided that further optimization of the hybridization of oligonucleotides is performed.

Our method bypasses reverse transcription biases^57^ which are linked to the capability of different enzymes to reverse transcribe molecules with high GC-content and complex secondary structure such as long hairpins formed by CUG and CCUG expanded repeats^58^.

We quantify tandem repeats with no need for amplification, another important bias source^59^, and we discriminate CUG tandem repeat arrays with 12, 24, 30 and 51 repeats. Notably, the 51-repeat array is associated with asymptomatic or mild DM1 phenotypes but is likely to expand into the full disease range upon paternal transmission^60^, showing that our sizing strategy holds diagnostic relevance. For GCN repeats, our method also successfully discriminates between the wild-type number of 20 repeats, and pathogenic 26 repeats associated with CCHS1, further demonstrating the applicability of this approach for diagnostics of multiple REDs with expansion sizes that result in significantly different clinical outcomes^3^.

Despite successful, single-molecule identification of biological targets in complex backgrounds has been proven challenging in previous studies, especially when the target is found in low copy numbers^61,62^. Here, we have successfully identified clinically relevant CUG-containing transcripts expressed in transfected human cell lines, without requiring reverse transcription, amplification or pull-down protocols. This represents an advantage over sequencing platforms, where small amounts and quality of starting materials remain limiting factors for diagnostic applications when amplification is not performed^63,64^.

Our quantification strategy also avoids the systematic errors associated to RNA sequencing in heteropolymer and short homopolymer regions^65^. The counting resolution of our method is determined by the number of hybridized DNA oligonucleotides to the target RNA. Here, we demonstrate a resolution of 6 trinucleotide repeats (18 nucleotides) which is achieved by the hybridization of a single additional oligonucleotide to the nanostructure. While our approach does require knowledge on the transcript sequence for nanostructure assembly, repeat quantification is entirely based on the number of complementary oligonucleotides that bind to the tandem repeats, offering determination of the size of the tandem repeat array while avoiding the systematic errors associated to RNA sequencing of heteropolymer regions.

In conclusion, we have introduced a reliable tool for detecting and quantifying tandem repeats in clinically relevant samples with a single-molecule resolution, offering an alternative amplification-free method for RED diagnostics, basic and clinical research of toxic RNAs.

## Methods and Materials

### DNA Plasmid Production

For constructs containing CTG repeats, plasmids were produced by cloning and subcloning reactions of CTG repeats originating from human DMPK locus using a pJet1.2/blunt cloning vector from CloneJET PCR Cloning Kit (Thermo Fisher Scientific, Catalog number K1232). The same approach was used to clone CCTG and GCN repeats originating from human *CNBP* and *PHOX2B* loci, respectively. Sanger sequencing was used to verify the sequence of all cloned and subcloned regions (**Table S17**, **S18**, **S19**). Sequencing of the entire constructs was done by the DNA Sequencing facility, Department of Biochemistry, University of Cambridge. Propagation of plasmids was performed in *Escherichia coli* JM110 strain upon chemical transformation, as previously described^35^. Plasmid DNA was purified using GeneJET Plasmid Miniprep Kit (Thermo Fisher Scientific, Catalog number K0503 and eluted in nuclease-free water (Qiagen, Catalog number 129117). The integrity of the plasmid was verified using agarose gel electrophoresis, and its concentration was determined using Qubit 2.0 fluorometer and Qubit™ dsDNA BR Assay Kit (Thermo Fisher Scientific, Catalog number Q32851).

### Fragment analysis

To verify the number of CTG and GCN repeats within constructs, PCR was performed with fluorescently labelled primers using Expand Long Template PCR System (Roche, Catalog number 11681842001) and amplicon lengths were determined using fragment analysis.For DNA constructs with CTG repeats, the reaction mixture of 15µl contained: 1× Expand Long Template Buffer 3 with 2.75 mM MgCl2 (Roche), 0.2 mM dNTPs (Thermo Fisher Scientific, Catalog number R0181), 0.04 μM M13_JP017 primer (5′-TGTAAAACGACGGCCAGTCAGCAAGATCATGGTGCAGTG-3′), 0.2 μM JP018 primer (5′-TTCCATGGCAGCTGAGAATATTG-3′), 0.2 μM FAM-M13 primer (5′-FAM-TGTAAAACGACGGCCAGT-3′), 1 M Betaine (Serva, Catalog number 14992), 0.3 U DNA Pol mix (Roche), 5-10 ng plasmid DNA and nuclease-free water (Qiagen, Catalog number 129117). PCR was performed using ProFlex™ 3 x 32-well PCR System (Thermo Fisher Scientific, Catalog number 4484073) with the following temperature profile: initial denaturation for 3 min at 96°C, 20 cycles of 30 sec at 95°C, 45 sec at 60°C, 45 sec at 68°C, 5 cycles of 30 sec at 95°C, 45 sec at 53°C, 45 sec at 68°C, followed by final elongation step for 30 min at 68°C. For DNA constructs with GCN repeats, the reaction mixture of 15µl contained: 1x Expand Long Template Buffer 3 with 2.75 mM MgCl2 (Roche), 0.2 mM dNTPs (Thermo Fisher Scientific, Catalog number R0181), 0.2 μM VIC-PHOX2B_F primer (5′-VIC-CCAGGTCCCAATCCCAAC-3′), 0.2 μM pJET1.2_R primer (5′-AAGAACATCGATTTTCCATGGCAG-3′), 1.5 M Betaine (Serva, Catalog number 14992), 0.3 U DNA Pol mix (Roche), 5-10 ng plasmid DNA and nuclease-free water (Qiagen, Catalog number 129117). PCR was performed using ProFlex™ 3 x 32-well PCR System (Thermo Fisher Scientific, Catalog number 4484073) with the following temperature profile: initial denaturation for 3 min at 96°C, 25 cycles of 30 sec at 95°C, 45 sec at 60°C, 1 min 30 sec at 68°C, followed by final elongation step for 30 min at 68°C. Amplified products were separated by capillary electrophoresis performed on the ABI3500 Genetic Analyzer (Applied Biosystems by Thermo Fisher Scientific, Catalog number 4406017) using the GeneScan™ 600 LIZ™ dye Size Standard v2.0 (Thermo Fisher Scientific, Catalog number 4408399) as an internal size standard

### RNA synthesis

RNA was produced by *in vitro* transcription of DNA constructs as reported previously^35^. Circular DNA constructs containing tandem repeats were linearized using DraIII-HF (New England Biolabs (NEB), Catalog number R3510S) following the manufacturer’s recommendations. The digestion reaction was performed at 37 °C for 1 h. The linearized DNA product was purified using the Monarch PCR & DNA Cleanup Kit (5 μg) (NEB, Catalog number T1030S) and the final DNA concentration was obtained with a NanoDrop spectrophotometer. *In vitro* transcription was then carried out using the HiScribe™ T7 Quick High Yield RNA Synthesis Kit (NEB, Catalog number E2050S) following the manufacture’s recommendations. Transcription was carried out at 37 °C for 4 h using 250 ng of DNA as template. The DNA template was subsequently removed from the transcription products by treating the reaction products with 4 units of DNase I (NEB, Catalog number M0303S). After DNA removal, the RNA products were purified using the Monarch RNA Cleanup Kit (50 μg) (NEB, Catalog number T2040S) for subsequent characterization in agarose gel electrophoresis and for nanostructure assembly.

### Assembly of RNA:DNA nanostructures

RNA:DNA nanostructures were assembled from combining tandem repeats-containing RNA with small single-stranded DNA oligonucleotides (**Figure S36**). The DNA oligonucleotides were complementary to the full-length RNA sequence which enabled the formation of the RNA:DNA backbone of the nanostructure. These DNA oligonucleotides also allowed for the integration of structural labels (such as DNA dumbbells) into the nanostructure, as well as for streptavidin binding. The assembly reaction was performed in 100 mM LiCl (Sigma-Aldrich, Catalog number L9650) and 1 × TE buffer (10 mM Tris-HCl buffer, 1 mM ethylenediaminetetraacetic acid, pH 8.0, Sigma-Aldrich, Catalog number T9285) to prevent magnesium fragmentation^66^ and nuclease-degradation of RNA, which relies on magnesium ions^67^. All buffers, and nuclease-free water (Ambion, Catalog number AM9937) used for this procedure were previously filtered in 0.22 µm Millipore syringe filter units (MF-Merck Millipore™, Catalog number GSWP04700) and treated with UV light for 10 minutes. A reaction volume of 40 μl was implemented, which included 800 fmol of the RNA scaffold (final concentration of 20 nM) and a five-fold excess of DNA staples, corresponding to 4 pmol of the complementary oligos (final concentration of 100 nM). For the tandem repeat labelling oligos, a five-fold excess relative to the available binding sites in the repeat array ensures complete RNA–DNA hybridization. A reduced oligonucleotide excess of threefold can also be used, while still yielding efficient hybridization and nanostructure assembly^30^. The assembly was done by incubating the reaction mixture at 70 °C for 30 s, followed by cooling down over 45 min to room temperature (90 cycles of 30 s with a 0.5 °C drop per cycle). Finally, to remove the excess of DNA oligonucleotides not bound to the nanostructure, the reaction product was filtered twice using 100 kDa cut-off Amicon filters. The purification step was performed using 10 mM Tris-HCl (pH 8.0) with 0.5 mM MgCl_2_ as a washing buffer. The sequence-selective assembly of the nanostructures and repeat hybridization was validated by nanopore sensing and polyacrylamide gel electrophoresis, as shown in **Figures S37** and **S38**.

### Electrophoretic Mobility Shift Assays

Agarose gel electrophoresis was used to evaluate the transcription products and the assembly of the RNA:DNA nanostructures. The assays were run on 1%(w/v) gels containing 1 × TBE buffer and 0.02% sodium hypochlorite. 1 × TBE buffer was used as running buffer and the samples (50 to 200 ng) were loaded with 1 × TriTrack loading dye (Thermo Fisher Scientific, Catalog number R1161). The assays were performed applying a constant voltage of 70 V for 180 minutes. Native polyacrylamide gel electrophoresis was also performed. We used Novex™ TBE Gels, 10% (Catalog number EC62752BOX). The samples were run following the manufacture’s recommendation with 1 × TriTrack loading dye (Thermo Fisher Scientific, Catalog number R1161), loading 150 to 200 ng of the sample to each lane and applying a constant potential of 80 V for 90 minutes. Upon electrophoresis, gels were stained in 3 × GelRed buffer (Biotium, Catalog number 41001) and imaged using a GelDoc-It™(UVP). Gel images were processed using ImageJ (Fiji)^68^, which was used to invert the grayscale, increase contrast, subtract the background and smoothen the surface.

### Nanopore chip fabrication

Nanopore measurements were performed in a polydimethylsiloxane (PDMS) device holding the solid-state nanopores. The nanopores were manufactured separately using a laser-assisted capillary puller P-2000 (Sutter Instruments, USA). We pulled quartz glass capillaries with 0.5 mm outer diameter and 0.2 mm inner diameter (Sutter Instruments, USA) which yielded two nanopores each. The puller parameters for nanopore fabrication were as follows: HEAT, 470; VEL, 25; DEL, 170; and PUL, 200; resulting in nanopores with diameters of 8 to 15 nm^29^. The capillaries are then placed in a PDMS device with radial geometry. The device can hold up to 16 solid-state nanopores, and each separates the *cis* and *trans* chambers of the device. The PDMS device was produced from the combination of a 10:1 (v/v) of PDMS monomer and curing agent (Sylgard 184 silicone elastomer kit, The Dow Chemical Company), which were mixed for 10 min and poured into a preheated mold (60 °C, 3 min) that defined the geometry of the PDMS device. After air bubbles exit the PDMS mixture poured into the mold, the PDMS was cured at 60°C for 48 hours. The glass nanopores were then positioned on the channels that separate the *trans* chambers from the *cis* chamber and the device was closed by pressing the PDMS holding the nanopores against a glass slide, after treating both components in air plasma for 15 seconds at maximum power (Femto, Diener). Finally, liquid PDMS mixture (10:1) was introduced to the channels holding the nanopores and the entire chip was heated in a hot plate at 150 °C for 30 min to solidify the PDMS and seal the chambers. Before each measurement, the nanopore chip was treated in air plasma for 5 minutes at maximum power (Femto, Diener) and 4M LiCl was added to the chambers.

### Nanopore measurements

For nanopore measurements, the RNA:DNA nanostructure was first diluted to 400 pM in 4 M LiCl, 1 × TE, pH 9.4 and it was subsequently introduced to the *cis* chamber of the nanopore chip. The measurements were performed by applying a fixed potential of 600 mV. The voltage was provided, and the corresponding current data was collected with an Axopatch 200B amplifier (Molecular Devices). The amplifier held a 100 kHz internal filter, and we used an eight-pole analog low-pass Bessel filter (900CT, Frequency Devices) with a cut-off frequency of 50 kHz to filter the output signal. The data were acquired using a sampling frequency of 1 MHz in a PCI-6251 data card (National Instruments). During the nanopore measurements, we monitor that there is no significant ionic current baseline drift that could considerably affect signal depth, dwell time, or event charge deficit (**Figure S39**). To minimize LiCl concentration changes caused by evaporation, the nanopore reservoir is effectively sealed within the PDMS device. In addition, a non-adhesive sealing film is placed over the reservoir opening to further reduce evaporation^55^.

After each measurement, we extracted individual translocation events from the ionic current trace recorded. Exemplary measurement durations are presented in **Table S20**. Subsequently, we used parameters such as the translocation time (> 0.05ms), the charge deficit (10 to 400 fC), the mean current of the translocation (< -100 pA), and the design of the RNA:DNA nanostructure to identify translocations ascribed to full-length unfolded RNA:DNA nanostructures. In **Figure S40**, we include a detailed workflow illustrating the quantitative distinction of truncated RNA products and the constructs assembled from full-length RNA. For each translocation event, we identified the number and position of downward current spikes and contrasted it to the initial design of the nanostructure. Once each event, corresponding to the translocation of an unfolded nanostructure, was linked to a specific design, we conducted further quantitative analysis. This included analysing the current drop and the area associated with the tandem repeat downward current spike. **Table S21** shows the IV curve for each nanopore used in the main text (ranging from -600 mV to 600 mV) along with the noise recorded at 600 mV. The Tukey’s Honest Significant Difference test (HSD) for distributions in **Figure 2** and **Figure S17** are included in **Table S22**.

### Production of plasmids expressing CUG tandem repeats

The vectors used for transient transfection in HEK293T cells were derived from pEGFP-N1 (Clontech). First, the BsaI restriction site in the original pEGFP-N1 vector was abolished using site directed mutagenesis to create the pEGFP-N1 ΔBsaI vector, in order to enable the following cloning steps. Subsequently, a synthetic fragment with 10 CTG repeats flanked with BsaI and BbsI restriction sites was inserted into the KpnI site in the pEGFP-N1 ΔBsaI vector. In order to obtain longer lengths of the CTG expansion, two cloning steps were performed using IIs restriction enzymes BsaI and BbsI, together with NotI as previously described^69^. In brief, the IIs restriction enzyme sites allowed the creation of a stepwise and seamless elongation of repetitive sequences and we have obtained the plasmid with 34 CTG repeats. The plasmids for transient transfection were purified using a miniprep kit (Qiagen) according to the manufacturers instructions. The concentration and purity of purified plasmids was determined using a microvolume spectrophotometer (Epoch, BioTek).

### Cell culture and transfection

HEK293T cells were cultured in Dulbecco’s Modified Eagle’s Medium (DMEM, Sigma-Aldrich), prepared from powder according to manufacturer’s instructions. The medium was supplemented with sodium bicarbonate buffer system, 10% (v/v) heat inactivated fetal bovine serum (Capricorn Scientific), and 1% (v/v) penicillin/streptomycin (Capricorn Scientific). Cells were maintained at 37°C in a humidified atmosphere containing 5% CO_2_ and passaged under aseptic conditions. To ensure that the cells were Mycoplasma-free, routine PCR testing for Mycoplasma contamination was performed through this study as previously described^70^.

Plasmid transfection was carried out using Lipofectamine 2000 (Life Technologies) according to the manufacturers instructions. Briefly, transfections were conducted in 24-well plates, where cells were transfected at 80% confluency using 1 µg of plasmid with 1 µL of Lipofectamine2000. The transfection mixture was made in DMEM devoid of serum and antibiotics, and applied dropwise to attached cells. The media was changed after 4 hours, and the cells were transferred to a 6-well plate on the following day. Fluorescence was observed two days post-transfection using Primovert inverted microscope (Zeiss) equipped with Axiocam 280 color camera (Zeiss). Image analysis was performed using ImageJ software^71^.

### Total RNA purification and quantification

HEK293T cells were trypsinized, collected and centrifuged at 5000 × g for 5 minutes. The supernatant was discarded and the cell pellet was resuspended in 50 µL of 1 × PBS. Total RNA was isolated from HEK293T cells using the Monarch Total RNA Miniprep kit (NEB, Catalog number T2010S) following the manufacturer’s instructions. RNA, treated with DNase I, was eluted using nuclease-free water, and RNAse inactivation reagent (RNA Secure, Thermo Fisher Scientific, Catalog number AM7005) was added per the manufacturer’s instructions. RNA concentration and purity were determined using a microvolume spectrophotometer (Epoch, BioTek) and verified with a microvolume fluorometer (Qubit, Thermo Fisher Scientific). RNA integrity was confirmed via agarose gel electrophoresis in 0.5 x TBE buffer.

### Reverse transcription and determination of absolute concentration of cDNA

The absolute concentration of target mRNA was determined using a previously described method involving quantitative PCR (qPCR) and standard RNA concentration curves^72,73^. Total RNA was reverse-transcribed into cDNA using random hexamer primers and the High-Capacity cDNA Reverse Transcription Kit (Thermo Fisher Scientific) following the manufacturer’s instructions. The resulting cDNA was used in qPCR with Power SYBR® Green PCR Master Mix (Thermo Fischer Scientific) and 0.25 µM of the following primers: FW GFP primer AAGCAGAAGAACGGCATCAA, Reverse GFP Primer GGGGGTGTTCTGCTGGTAGT. The temperature profile was 95 °C 10 minutes, 40 × cycles (95 °C 15 seconds, 60 °C 60 seconds). The qPCR was performed using Step One Plus Real-Time PCR system (Applied Biosystems). To generate a standard curve for GFP cDNA quantification, the pEGFP-N1 plasmid was serially diluted to known concentrations. Ct values from these dilutions were plotted against the number of plasmid molecules, and a regression analysis was performed to derive a standard equation. The Ct values from the experimental samples were applied to this equation to calculate the absolute number of mRNA molecules. Using the known molecular weight of the target mRNA, the absolute concentration of mRNA in the sample was determined.

### MS2 nanostructure assembly at 0.5 nM concentration

The assembly reactions took place in 100 mM LiCl, 1 × TE buffer, using prefiltered solutions in 0.22 µm Millipore syringe filter units (MF-Merck Millipore™, Catalog number GSWP04700). A reaction volume of 40 μl was implemented, which included 20 fmol of the MS2 RNA (Roche, Catalog number 10165948001) scaffold (0.5 nM in final reaction volume) and a ten to five hundred times excess of DNA oligonucleotides. Assembly was done by incubating the reaction mixture at 70 °C for 30 s and cooling down over 45 min to room temperature (90 cycles of 30 s with 0.5 °C drops per cycle).

### CUG repeats mRNA nanostructure assembly and detection in total RNA background

The assembly reaction took place in 100 mM LiCl, 1 × TE buffer, using prefiltered solutions in 0.22 µm Millipore syringe filter units (MF-Merck Millipore™, Catalog number GSWP04700). A reaction volume of 20 μl was implemented, which included 10 μl of total human RNA (2.1 nM of CUG mRNA in 1.6 μg/ μl), and an excess of two hundred times excess of DNA oligonucleotides with respect to the CUG mRNA. Assembly was done by incubating the reaction mixture at 70 °C for 30 s and cooling down over 45 min to room temperature (90 cycles of 30 s with 0.5 °C drops per cycle). Finally, the reaction mixture was filtered three times in 100 kDa cut-off Amicon filters, using 10 mM Tris-HCl (pH 8.0) with 0.5 mM MgCl_2_ as a washing buffer for each purification step. Extraction method of translocation events is presented in **Figure S41**.

## Statistics and reproducibility

Nanopore experiments were conducted continuously to collect as many translocation events as possible. Experient duration is often limited to partial clogging of the nanopore, for this reason sample size varies between experiments. For *in vitro* transcribed RNA samples, we only present data set where we measure at least 100 translocation events ascribed to nanostructures passing through the pore in an unfolded conformation. For all experiments, we perform further analysis on unfolded full-length RNA translocations. We perform each experiment in triplicate. The experiments were not randomized. The investigators were not blinded to allocation during experiments and outcome assessment.

## Code availability

The data presented was analysed using home-built software written with National Instruments LabVIEW 2017 and Python. All the data presented in this work and in the Supplementary Materials can be plotted and analysed manually or using any software of preference. Software for facilitating extraction of translocation events from samples with complex backgrounds are available at https://github.com/maxearle/nas2.

## Acknowledgements

G.P.-G. acknowledges funding from EPSRC CDT MRes/PhD Studentship in Nanoscience and Nanotechnology (NanoDTC Cambridge EP/S022953/1) and Trinity-Henry Barlow Scholarship. U.F.K. acknowledges funding from UK Research and Innovation (UKRI) under the UK government’s Horizon Europe funding guarantee EP/ X023311/1, European Research Council (ERC) consolidator grant (DesignerPores no. 647144), an ERC-2019-PoC grant (PoreDetect no. 899538) and European Union under the Horizon 2020 Program, FET- Open: DNA-FAIRYLIGHTS, Grant Agreement No. 964995. F.B. acknowledges funding from the George and Lilian Schiff Foundation Studentship, the Winton Program for the Physics of Sustainability PhD Scholarship, and St John’s College Benefactors’ Scholarship. J.P., M.P., A.N. and D.S.-P. acknowledge support by grants from the Science Fund of the Republic of Serbia, Grant number #7754217, *Understanding repeat expansion dynamics and phenotype variability in myotonic dystrophy type 1 through human studies, nanopore sequencing and cell models –* READ-DM1. M. E. acknowledges funding from the European Union under the Horizon 2020 Program, FET-Open: DNA-FAIRYLIGHTS, Grant Ahorigreement No. 964995.

S.P. acknowledges funding from EPSRC CDT MRes/PhD Studentship in Nanoscience and Nanotechnology (NanoDTC Cambridge EP/S022953/1). We thank Dr. Casey Platnich and Dr. David Jordan for reading the manuscript and Thieme Schmidt and Simon Brauburger for insightful discussions.

## Data availability

Data supporting the findings of this study are available in the Main text and the Supplementary Materials. Current traces are provided in Main text and Supplementary Materials, and source data is provided with this paper.

