## Supplementary Material for "Quantification of disease-associated RNA tandem repeats by nanopore sensing"

Patiño-Guillén, G.[1], Pešović, J.[2], Panic, M.[2,3], Earle, M. [1], Ninković, A.[2], Petrușca, S. [1], Savic-Pavicevic, D.[2,*], Keyser, U.F. [1,*], Bošković, F.[1,*]

This PDF file includes:

→ Figures S1 to S41

→ Tables S1 to S22

**Figure S1.**

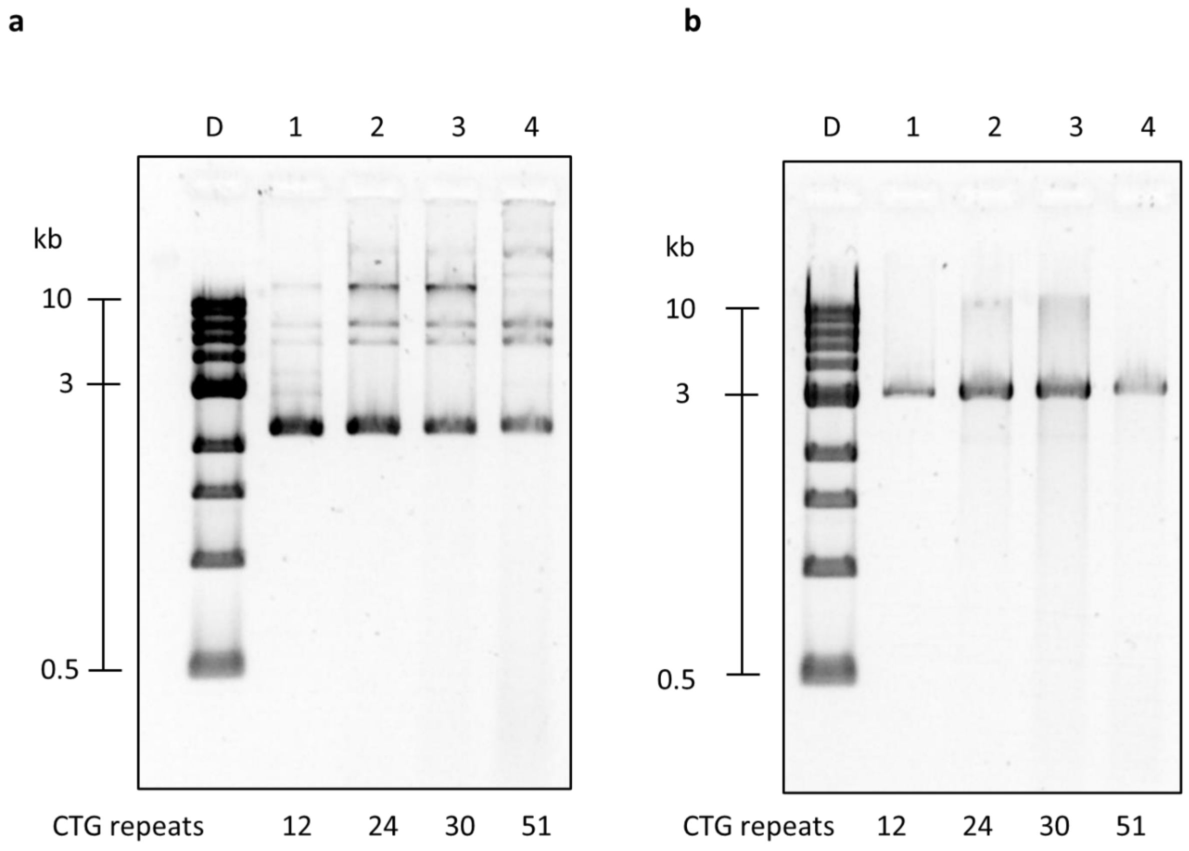


**Figure S1.** Linearization of DNA constructs containing CTG repeats with restriction enzyme DraIII. **a** Four ~3 kbp DNA constructs are presented, each containing a different number of CTG repeats. Lane D: 1 kbp DNA ladder, lane 1: circular DNA with 12 CTG repeats, lane 2: 24 CTG repeats, lane 3: 30 CTG repeats, lane 4: 51 CTG repeats. **b** The DNA constructs were digested using restriction enzyme DraIII producing linear DNA templates for *in vitro* transcription. Gels: 1 % (w/v) agarose, 1 × TBE, 0.02% sodium hypochlorite.

**Figure S2.**

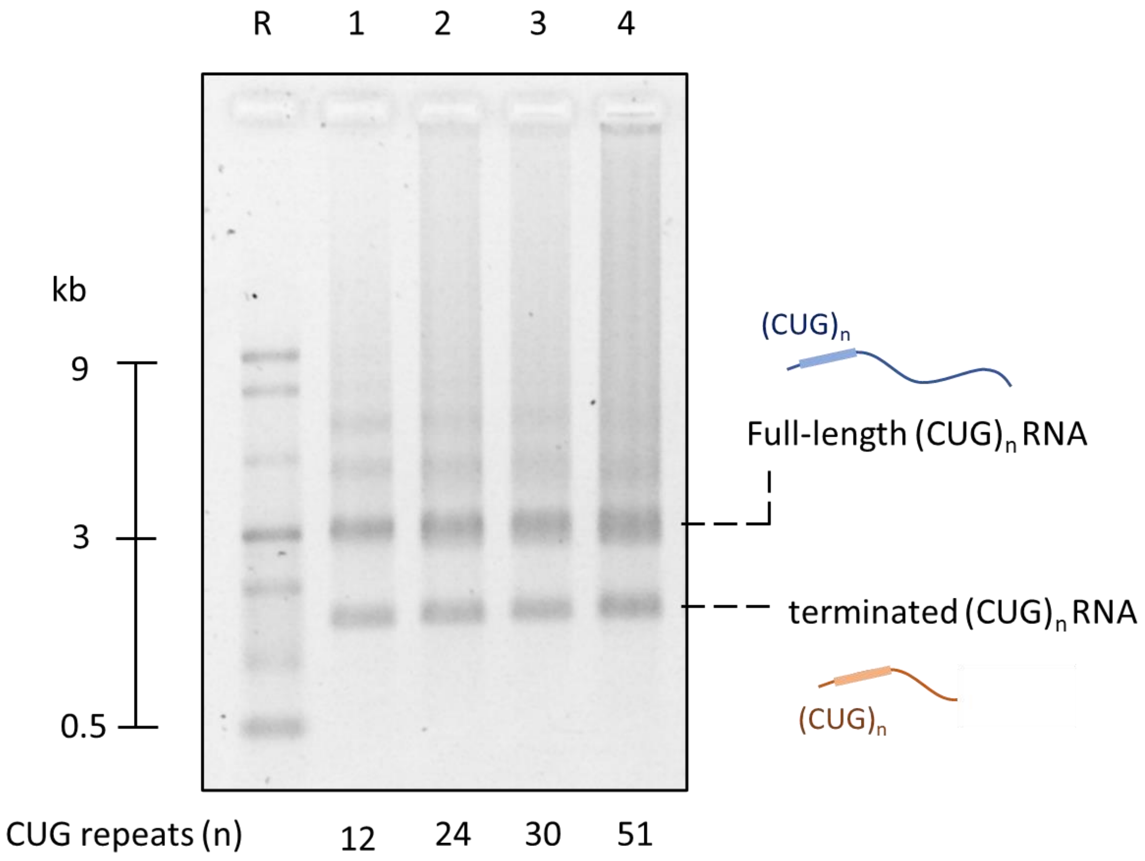


**Figure S2.** *In vitro* transcription of CTG repeats-containing DNA constructs to produce CUG repeats-containing RNA. ~3 kbp linearized DNA constructs containing CTG repeats were in vitro transcribed with T7RNAP. Transcription yielded full-length RNA with the corresponding number of CUG repeats. Prematurely terminated transcripts containing the repeats were also produced due to the presence of a premature transcription termination sequence in the DNA constructs, as extensively studied in[1]. For RNA barcoding and characterization of the repeats, in this study we focus on the full-length transcripts. R corresponds to RNA ladder. Gel: 1 % (w/v) agarose, 1 × TBE, 0.02% sodium hypochlorite.

#### Figure S3.

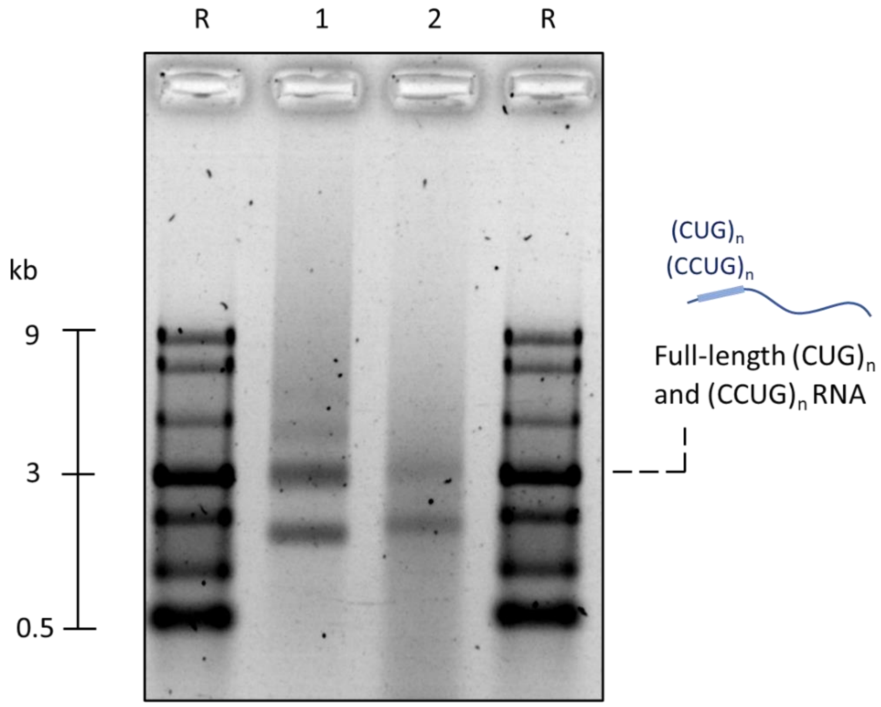


**Figure S3.** *In vitro* transcription of CTG and CCTG repeats-containing DNA constructs to produce CUG and CCUG repeats-containing RNA. ~3 kbp linearized DNA constructs containing CTG and CCTG repeats were in vitro transcribed with T7RNAP. Transcription yielded full-length RNA with the corresponding number of CUG (lane 1) and CCUG repeats (lane 2). R corresponds to RNA ladder. Gel: 1 % (w/v) agarose, 1 × TBE, 0.02% sodium hypochlorite.

**Figure S4.**

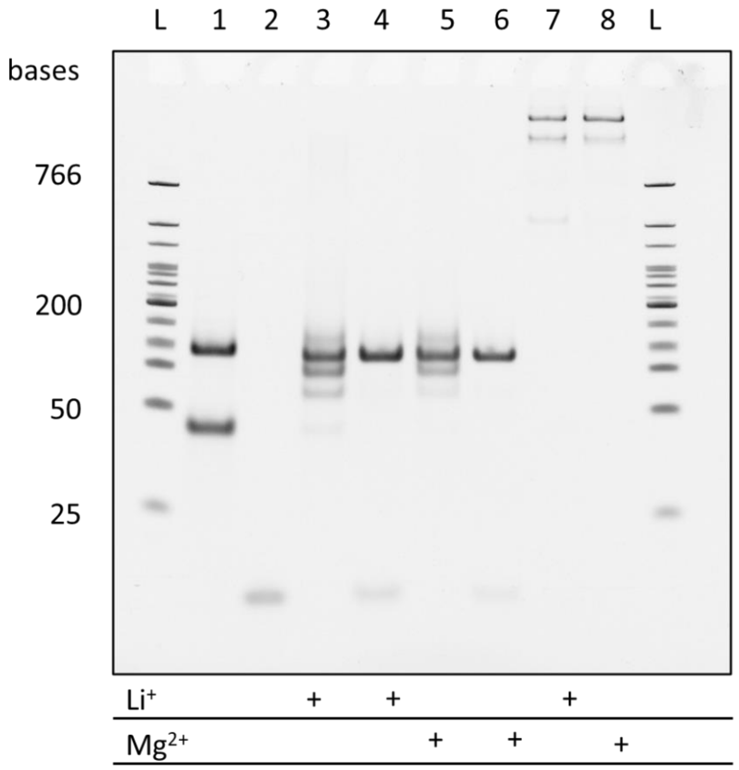


**Figure S4.** Labelling of CTG tandem repeats with monovalent streptavidin through hybridization with complementary DNA oligonucletoides. Lane L: Low range DNA ladder. Lane 1: $(CTG)_{30}$ DNA. Lane 2: $(CAG)_{10}$-biotin DNA. Lane 3: $(CTG)_{30}$ + $(CAG)_{10}$-biotin in ~ 1:3 ratio hybridized in 100 mM LiCl. The 1:3 molar excess corresponds to a 1:1 excess of $(CAG)_{10}$-biotin with respect to the available binding sites for full hybridization in the 30 CTG array. Lane 4: $(CTG)_{30}$ + $(CAG)_{10}$-biotin in ~ 1:9 ratio hybridized in 100 mM LiCl. The 1:9 molar excess corresponds to a 1:3 excess of $(CAG)_{10}$-biotin with respect to the available binding sites for full hybridization in the 30 CTG array. Lane 5: $(CTG)_{30}$ + $(CAG)_{10}$-biotin in ~ 1:3 ratio hybridized in 10 mM $MgCl_2$. Lane 6: $(CTG)_{30}$ + $(CAG)_{10}$-biotin in ~ 1:9 ratio hybridized in 10 mM $MgCl_2$. Lane 7: $(CTG)_{30}$ + $(CAG)_{10}$-biotin in a 1:9 ratio hybridized in 100 mM LiCl with excess of monovalent streptavidin. Lane 8: $(CTG)_{30}$ + $(CAG)_{10}$-biotin in a 1:9 ratio hybridized in 10 mM $MgCl_2$ with excess of monovalent streptavidin. Gel: 10 % polyacrylamide, 1 × TBE. Each $(CTG)_{30}$ DNA strand hybridizes to 3 $(CAG)_{10}$ oligonucleotides. The assay demonstrates efficient hybridization and streptavidin labelling of the $(CTG)_{30}$ repeats upon reaction with excess $(CAG)_{10}$ oligonucleotides in both 100 mM LiCl and 10 mM $MgCl_2$. Lanes 4 and 6 show a sharp band, which indicates a predominant assembly conformation, while lanes 7 and 8 exhibit constructs with lower mobility, attributed to streptavidin attachment.

#### Figure S5.

**a**

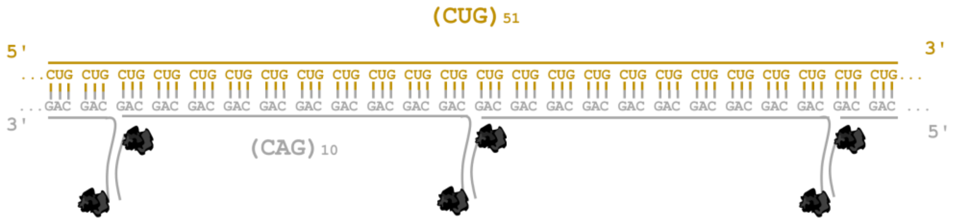


| Oligonucleotide | Sequence (5′→ 3′) |
|---|---|
| $(CAG)_{10}$ | CAGCAGCAGCAGCAGCAGCAGCAGCAGCAGTTTGGATATCACTCATTAGTGGT/ 3′-biotin/ |
| Second strand | ACCACTAATGAGTGATATCC/3′-biotin/ |

**b**

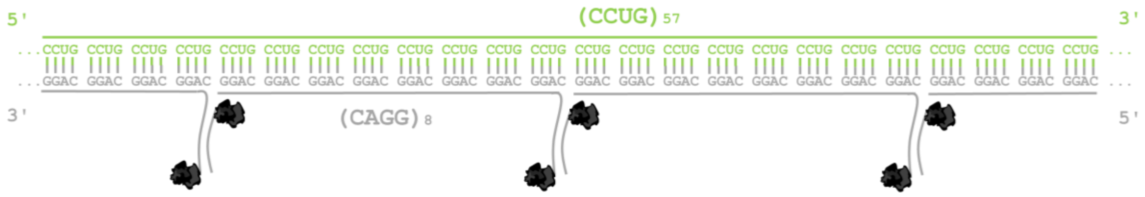


| Oligonucleotide | Sequence (5′→ 3′) |
|---|---|
| $(CAGG)_{8}$ | CAGGCAGGCAGGCAGGCAGGCAGGCAGGCAGGTTTGGATATCACTCATTAGTGGT/ 3′-biotin/ |
| Second strand | ACCACTAATGAGTGATATCC/3′-biotin/ |

**Figure S5.** Schematic of RNA tandem repeat expansion labelling. **a** CUG tandem repeats are detected through hybridization of $(CAG)_{10}$ oligonucleotides which hold an overhang biotinylated sequence. This overhang enables the binding of a complementary biotinylated DNA strand. Each DNA strands binds one streptavidin which is used as structural label to sense the CUG repeats. **b** Similarly, $(CAGG)_8$ oligonucleotides are used to sense CCUG repeats, each enabling the binding of two streptavidins.

**Figure S6.**

a

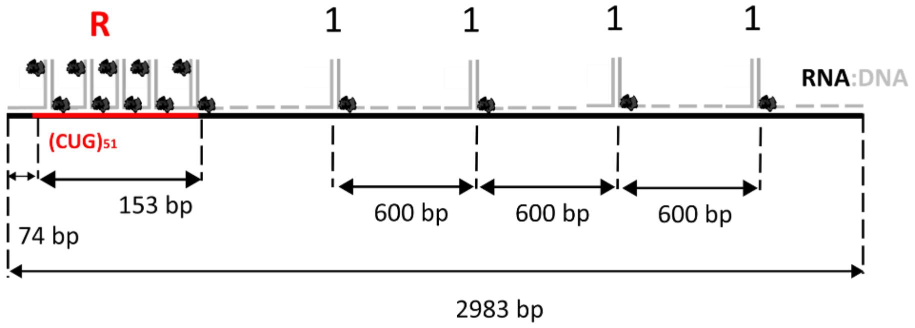


b

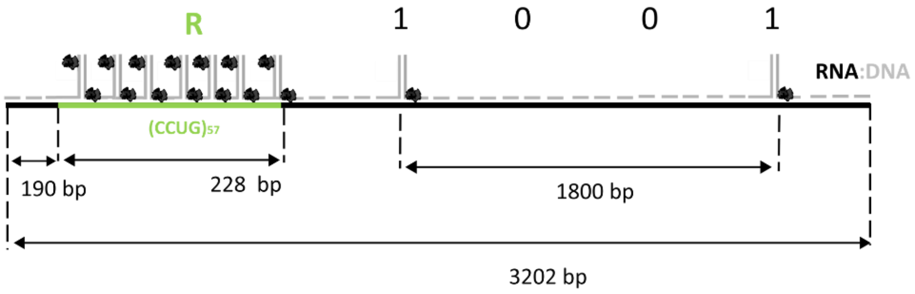


**Figure S6.** Detailed design of RNA:DNA nanostructures used for detection of tandem repeats. **a** The RNA:DNA nanostructure tagged with '1111' is used to detect RNA containing 51 CUG tandem repeats and **b** RNA:DNA nanostructure tagged with a '1001' barcode is used to identify RNA containing 57 CCUG.

**Figure S7.**

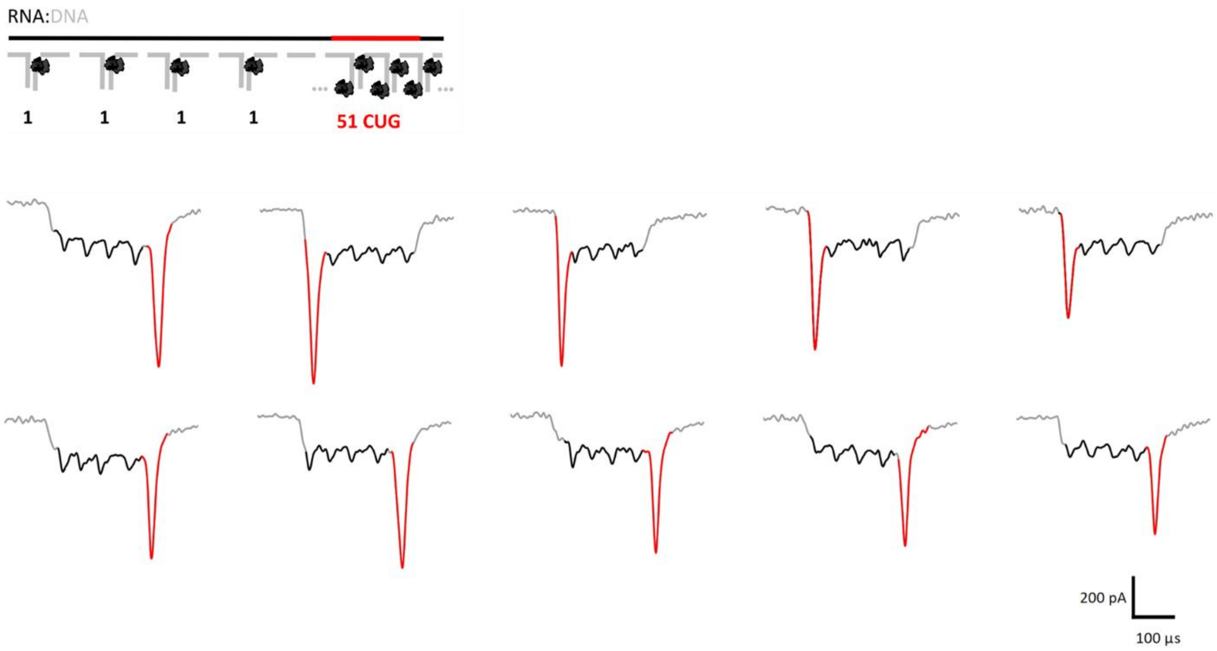


**Figure S7.** Nanopore translocation events of RNA:DNA nanostructures tagged with '1111' barcode, used to detect RNA containing 51 CUG tandem repeats. The current spike ascribed to the repeats can be found either at the beginning or at the end of the ionic current drop as the molecules can translocate in the 5'-to-3' direction or in the 3'-to-5' direction through the nanopore.

**Figure S8.**

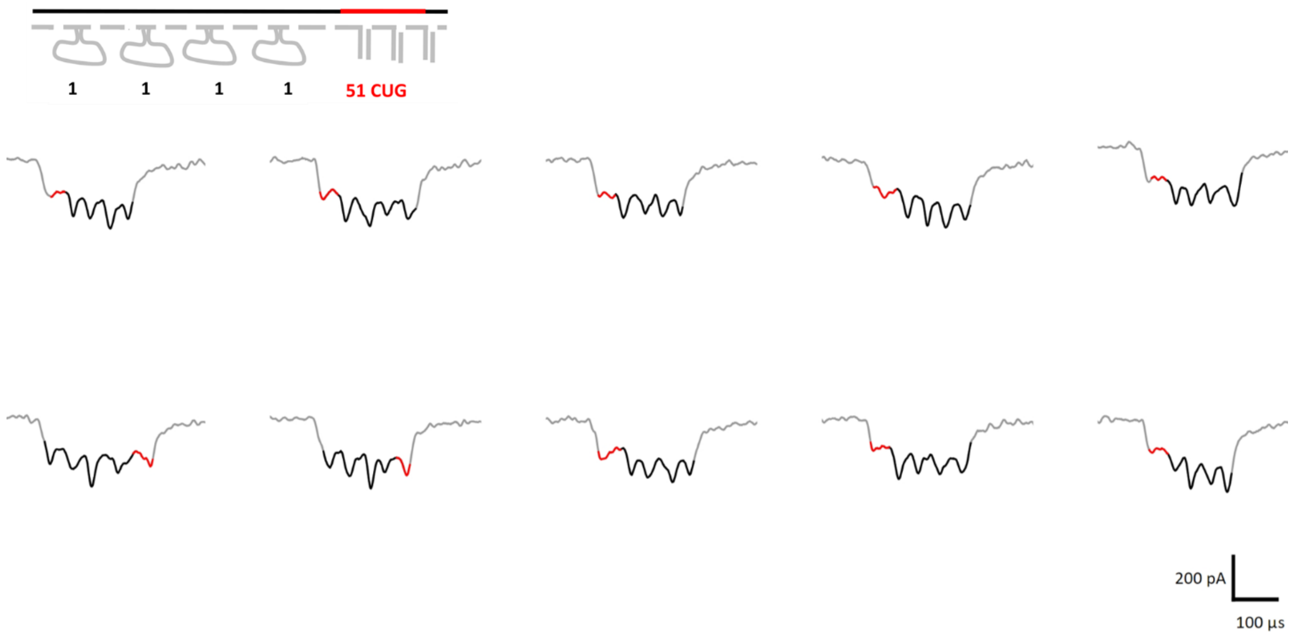


**Figure S8.** Nanopore translocation events of unfolded RNA:DNA nanostructures tagged with '1111' barcode, where each '1' is produced from 12 consecutive DNA dumbbells. The 51 CUG tandem repeats are not labelled with monovalent streptavidin.

**Figure S9.**

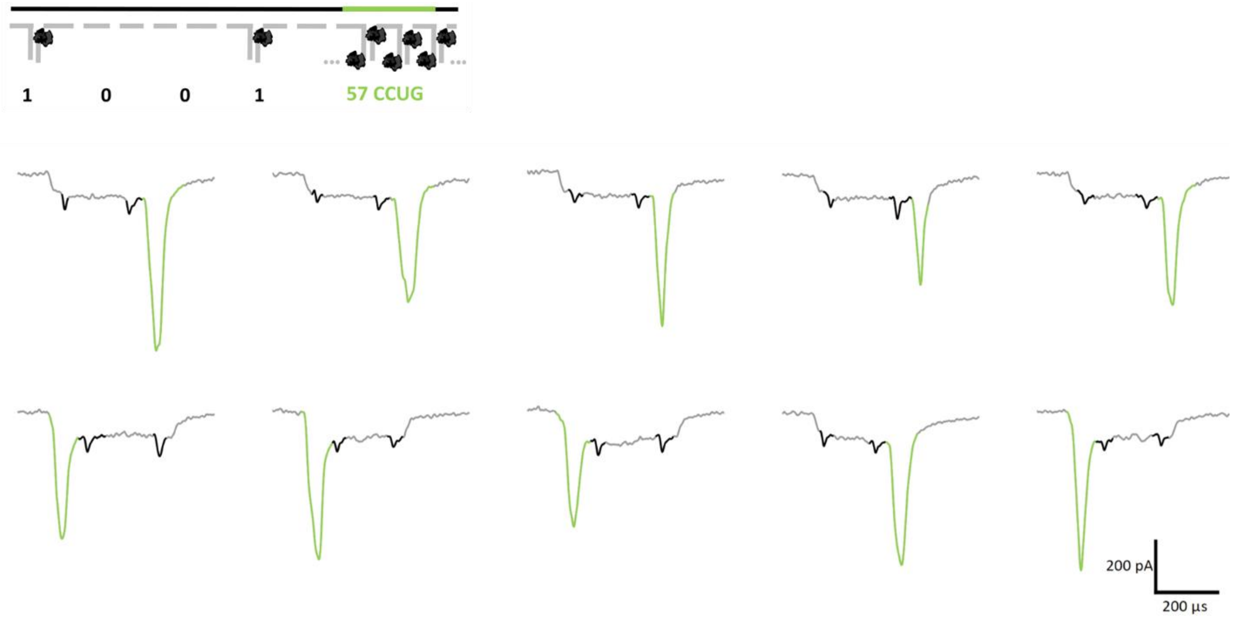


**Figure S9.** Nanopore translocation events of unfolded RNA:DNA nanostructures tagged with '1001' barcode, where the 57 CCUG tandem repeats are labelled with monovalent streptavidin. The current spike ascribed to the repeats can be found at the beginning or at the end of the ionic current drop as the molecules can translocate in the 5'-to-3' direction or in the 3'-to-5' direction through the nanopore.

**Figure S10.**

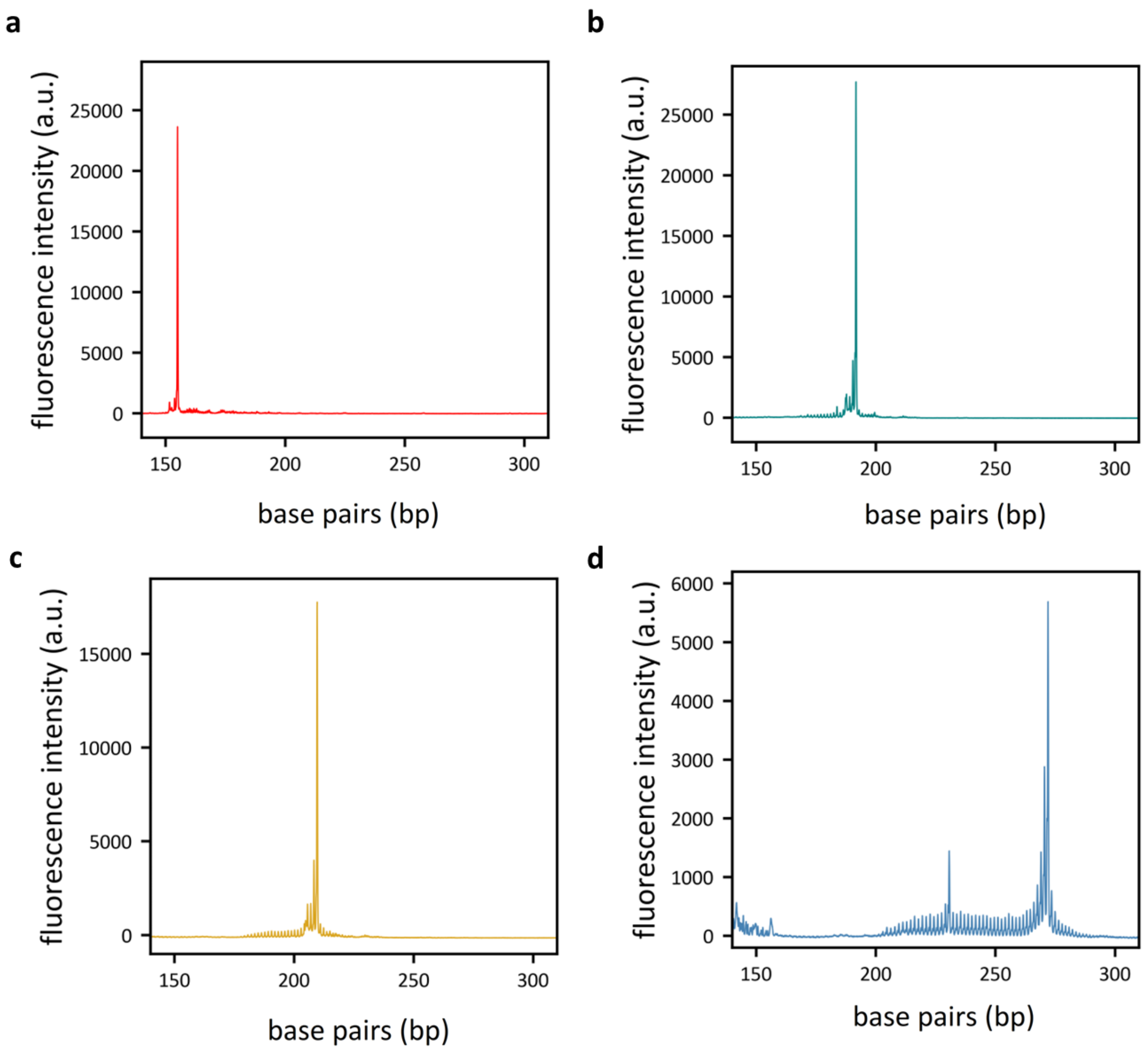


**Figure S10.** Size distribution of CTG tandem repeats in DNA used for *in vitro* transcription. The fragment analysis is presented for **a** 12, **b** 24, **c** 30, and **d** 51 repeats. Fragment analysis was performed using GeneScan 600 LIZ as a size standard.

**Figure S11.**

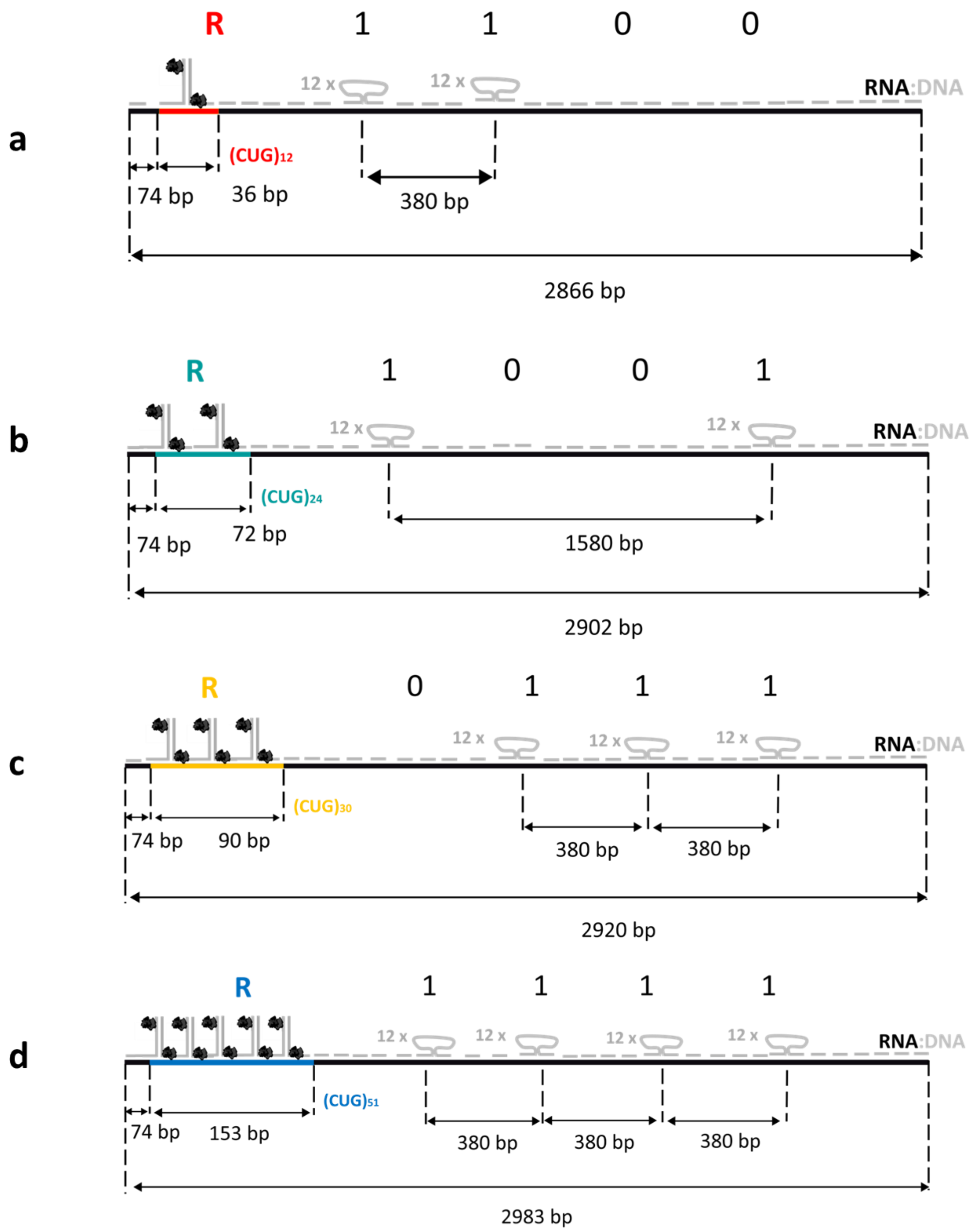


**Figure S11.** Detailed design of RNA:DNA nanostructures used for quantitative analysis of CTG tandem repeats. **a** The nanostructure tagged with a ‘0011’ is used to study 12 CUG tandem repeats, **b** RNA:DNA nanostructure tagged with a ‘1001’ barcode is used to study 24 CUG, **c** RNA:DNA nanostructure tagged with a ‘1110’ barcode is used to study 30 CUG, **d** RNA:DNA nanostructure tagged with a ‘1111’ barcode is used to study 51 CUG repeats.

**Figure S12.**

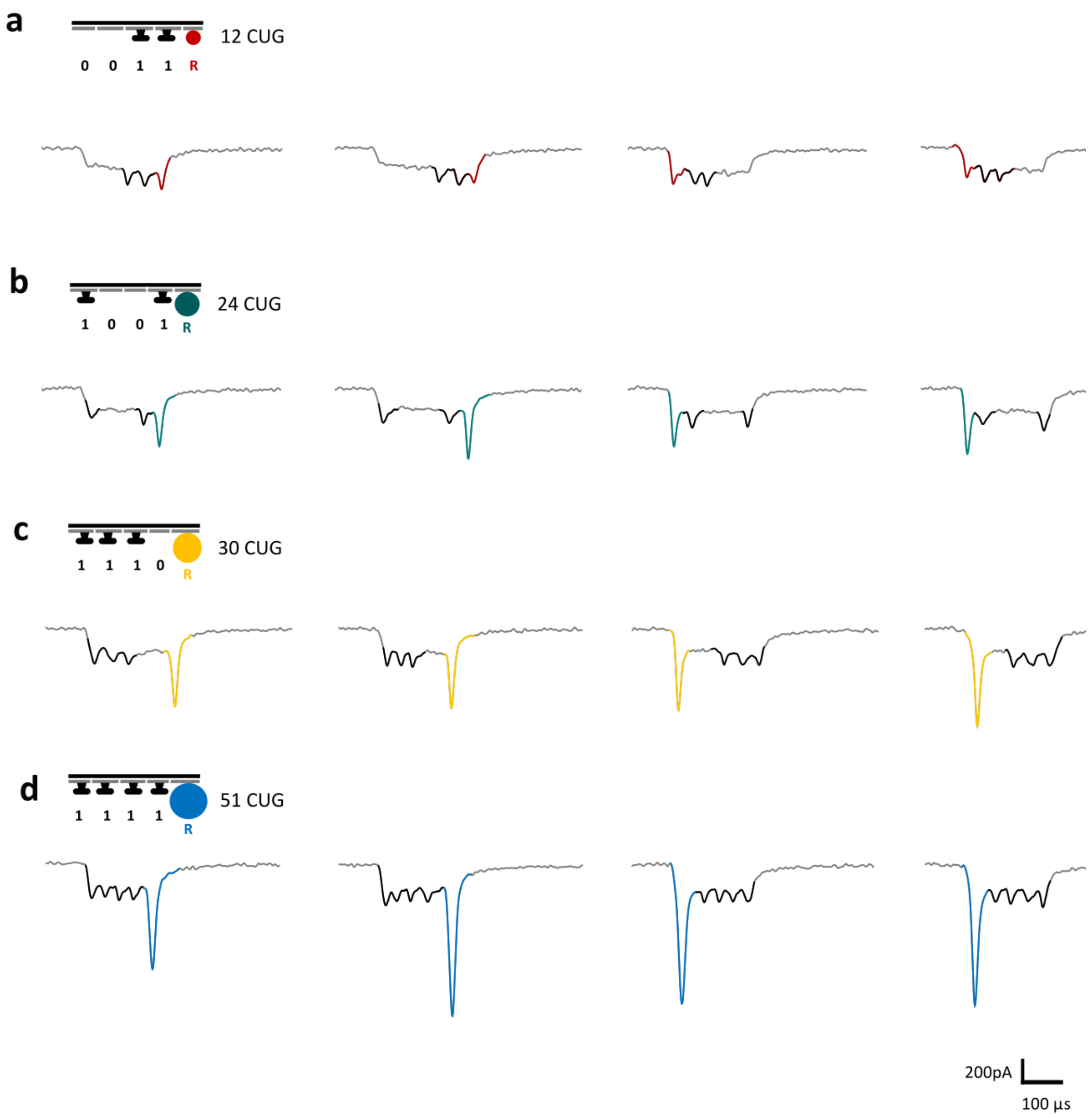


**Figure S12.** Nanopore translocation events of unfolded RNA:DNA nanostructures tagged with **a** '1111' for quantitative analysis of 12 CUG repeats, **b** '1001' for 24 CUG repeats, **c** '1110' for 30 CUG repeats, **d** '1111' for 51 CUG repeats.

**Figure S13.**

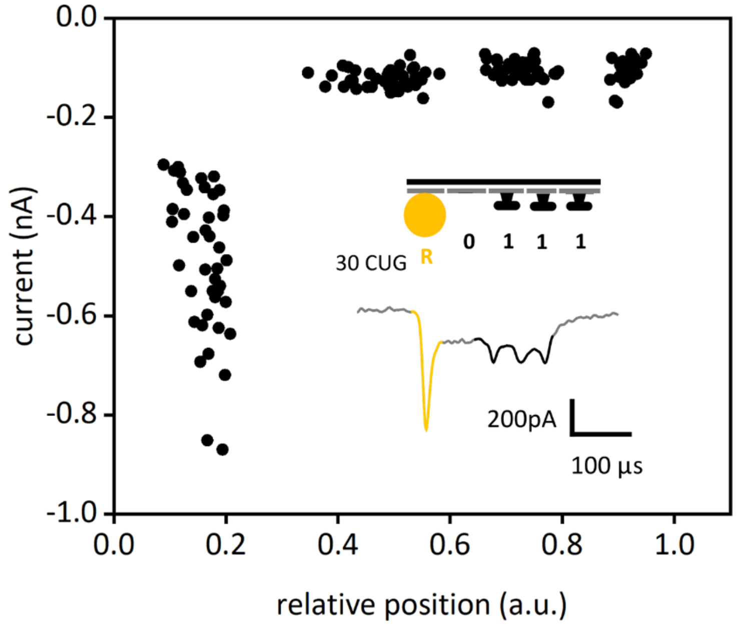


**Figure S13.** Quantitative description of the current modulation produced by the translocation of a '1110' tagged nanostructure with 30 CUG repeats. The spike depth and relative position of each current spike produced by either the dumbbells of the barcode or the labelling of CUG repeats are presented.

#### Figure S14.

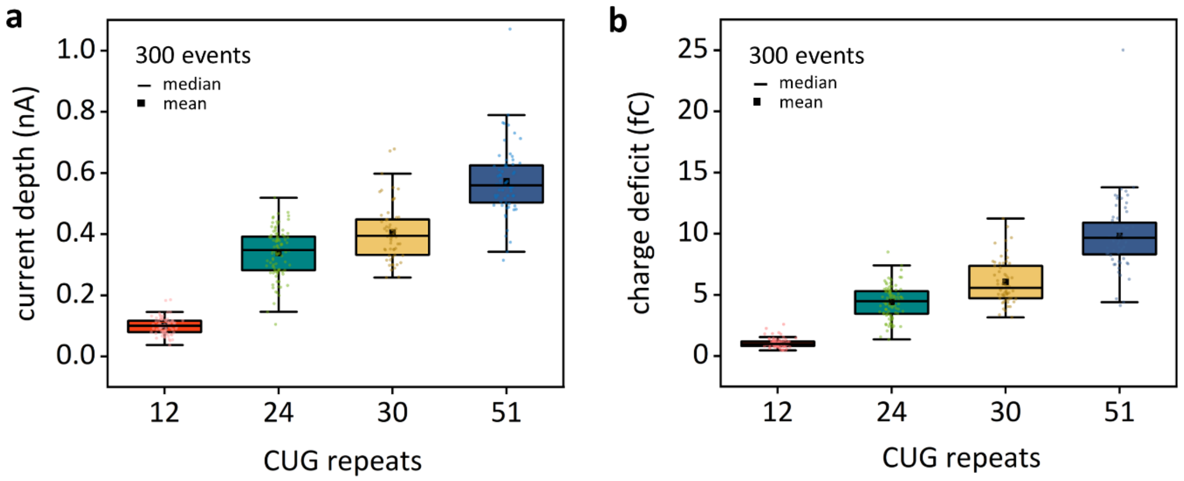


**Figure S14.** Additional dataset for quantitative analysis of tandem repeats. We provide supplementary dataset that includes detailed examination of the spike current's depth and area across four distinct CUG repeat array sizes. **a** The dataset shows median spike current depths of 0.10 nA, 0.34 nA, 0.39 nA, 0.55 nA and **b** spike areas of 0.98 fC, 4.49 fC. 5.57 fC, 9.66 fC. The dataset offers a comprehensive view of the variations in current characteristics associated with each repeat array size.

### Figure S15

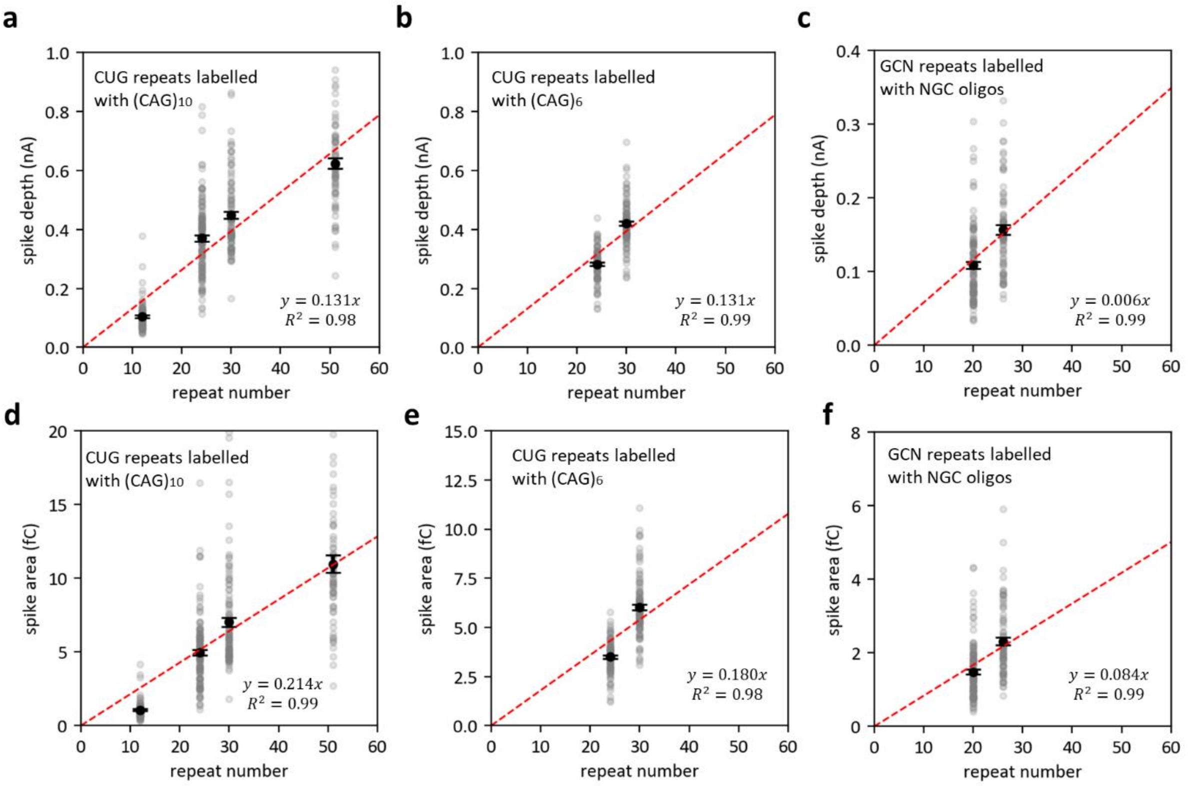


**Figure S15**. Correlation of repeat array length to current spike observables. The expected repeat array length based on the spike depth for characterization of **a** 12, 24, 30 and 51 CUG repeats with $(CAG)_{10}$ oligonucleotides. **b** 24 and 30 CUG repeat arrays with $(CAG)_6$ oligonucleotides. **c** 20 and 26 GCN repeats with NGC oligonucleotides. The expected repeat array length based on the spike area for characterization of the same CUG repeats with **d** $(CAG)_{10}$ and **e** $(CAG)_6$ oligonucleotides, and **f** the same GCN repeats with NGC oligonucleotides. The mean of each distribution is indicated. Error bars indicate standard error.

### Figure S16

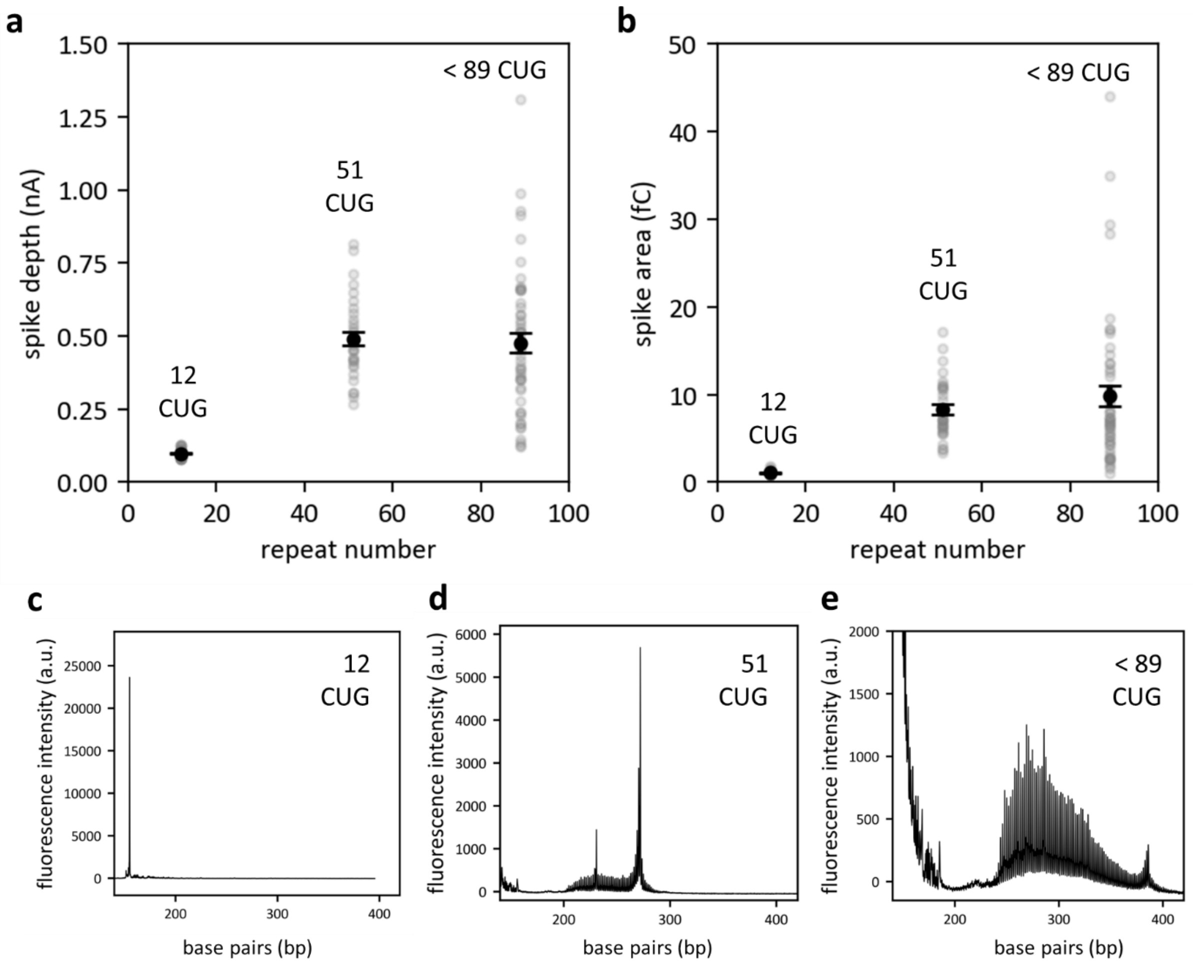


**Figure S16**. Correlation of repeat array length to current spike observables including the spike depth and spike area. We show the spike **a** depth and **b** area for a CUG repeat array of 12 repeats, 51 repeats and a mix of repeat lengths up to 89 repeats. The mean of each distribution is indicated. Error bars indicate standard error. Panels **c**, **d**, and **e** show the fragment analysis for each repeat array, which were performed using GeneScan 600 LIZ as the size standard.

**Figure S17.**

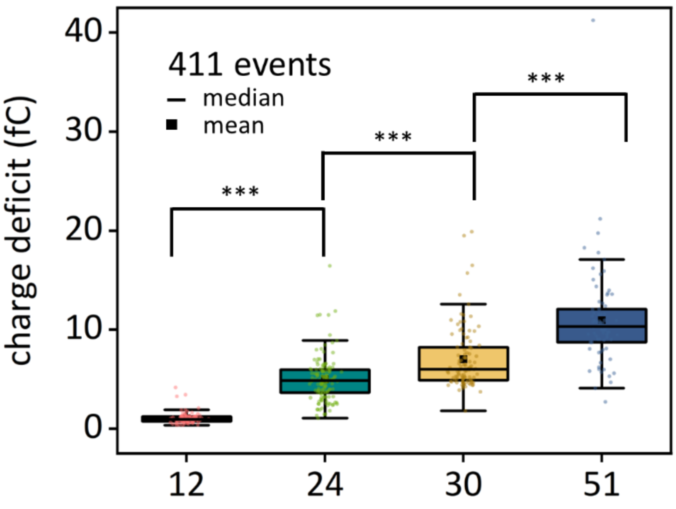


**Figure S17.** Quantitative profiling of tandem repeats was performed though the analysis of the current spike ascribed to the streptavidin bound to the repeats region while translocating through the nanopore. In Figure 2, we use the depth of this current spike ascribed to compare the repeat array sizes. Here, we show that the area of the spike increases its magnitude with a greater number of repeats. The spike areas are 0.94 fC, 4.86 fC, 6.00 fC, and 10.31 fC. Welch's t-test ***p-values < 0.00001: $7.1\times10^{-44}$, $6.39\times10^{-8}$, $2.88\times10^{-8}$. A one-way ANOVA test f-ratio value is 163.3 and the p-value is < 0.00001.

### Figure S18.

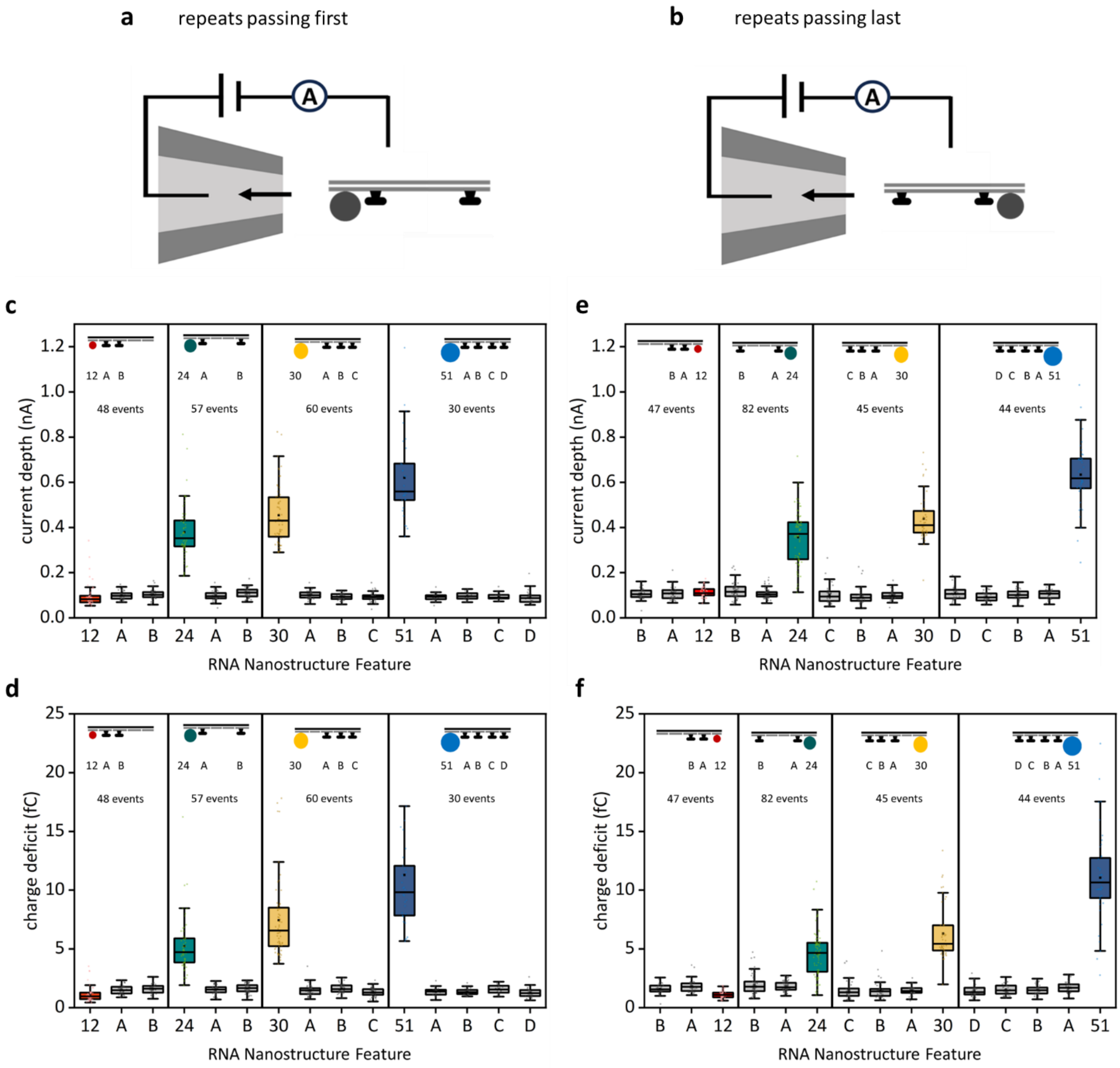


**Figure S18.** Tandem repeats can be quantified *via* nanopore sensing independently of the direction of the free translocation. The RNA:DNA nanostructure containing tandem repeats can pass through the nanopore either **a** in the 5′ to 3′ direction (repeats translocate first) or **b** in the 3′ to 5′ direction (repeats translocate at the end). We show that the current spike depth ascribed to the repeats and its area increase with the number of repeats independently of whether the translocation occurs in the 5′ to 3′ direction (**c**, **d**) or the 3′ to 5′ direction (**e**, **f**).

**Figure S19.**

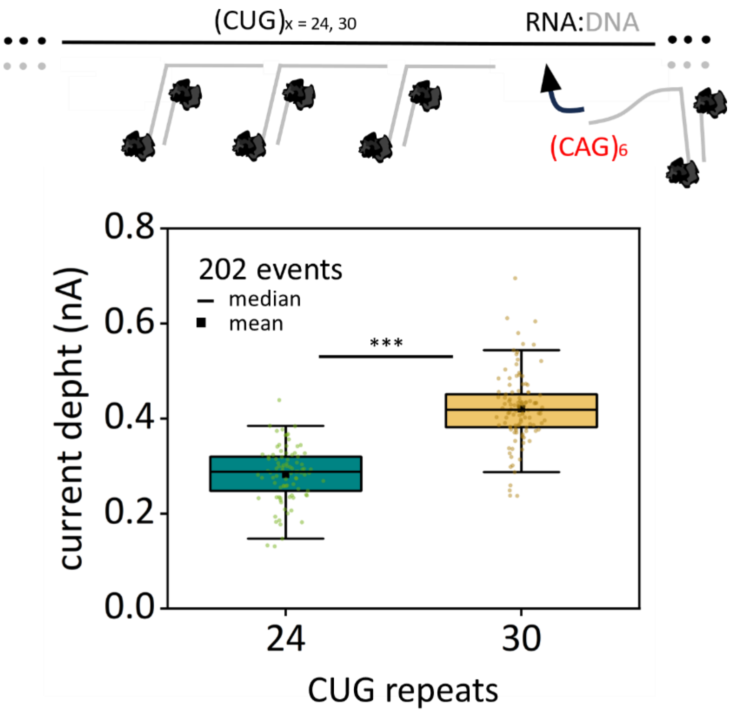


**Figure S19.** The oligonucleotides used for labelling tandem repeats can be adjusted to the repeat array sizes to enhance quantification precision. For clearer distinction between 24 and 30 CUG repeats, $(CAG)_6$ oligonucleotides can be implemented, where 4 and 5 oligonucleotides bind to the CUG repeats, respectively. Nanopore measurements yield downward current spikes with median values of 0.29 nA and 0.42 nA, for 24 and 30 repeats. p-value (Welsch test) was $2.38\times10^{-33}$, a higher difference compared to labelling with $(CAG)_{10}$ oligonucleotides, where a third oligonucleotide may or may not bind to the 24 CUG expansion.

#### Figure S20.

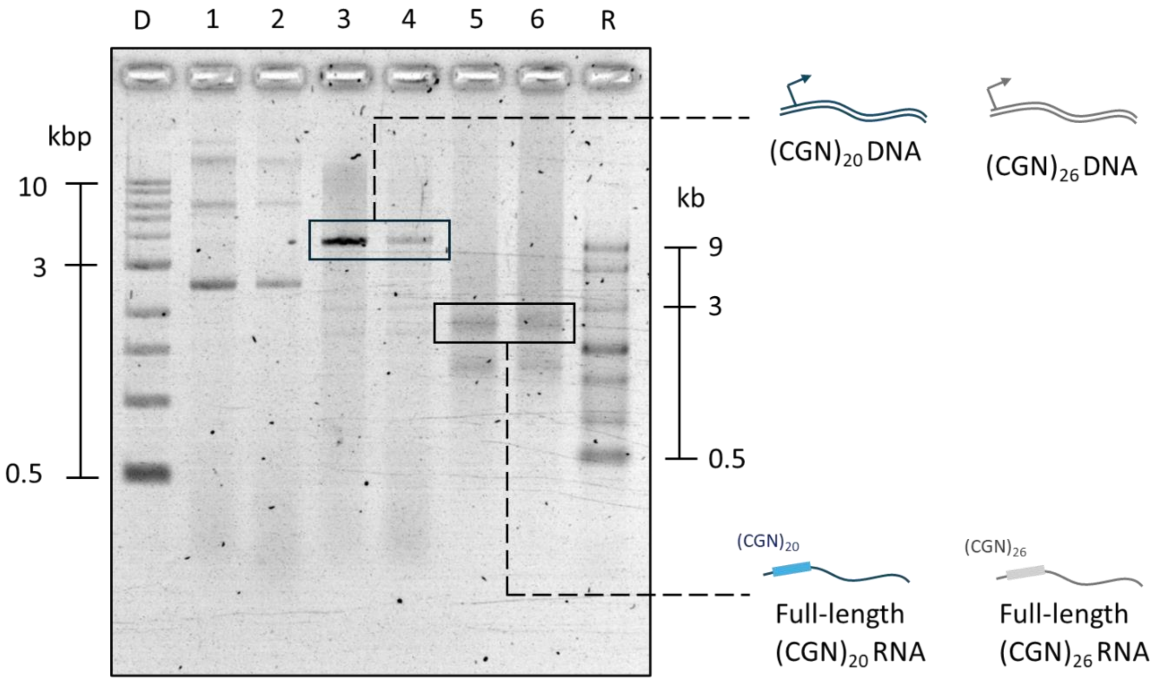


**Figure S20.** *In vitro* transcription of linear DNA construct with GCN repeats. DNA ladder and ssRNA ladder are included on both sides of the gel, in lanes D and R, respectively. Lane 1: 3.7 kbp circular DNA with 20 GCN repeats (sequence in Table S9). Lane 2: 3.7 kbp Circular DNA with 26 GCN repeats (sequence in Table S10). Lane 3: DNA construct from lane 1 linearized with DraIII restriction enzyme. Lane 4: DNA construct from lane 2 linearized with DraIII. Lane 5: RNA from *in vitro* transcription of DNA from lane 3. Lane 6: RNA from *in vitro* transcription of DNA from lane 4. Gel: 1 % (w/v) agarose, 1 × TBE, 0.02% sodium hypochlorite.

#### Figure S21

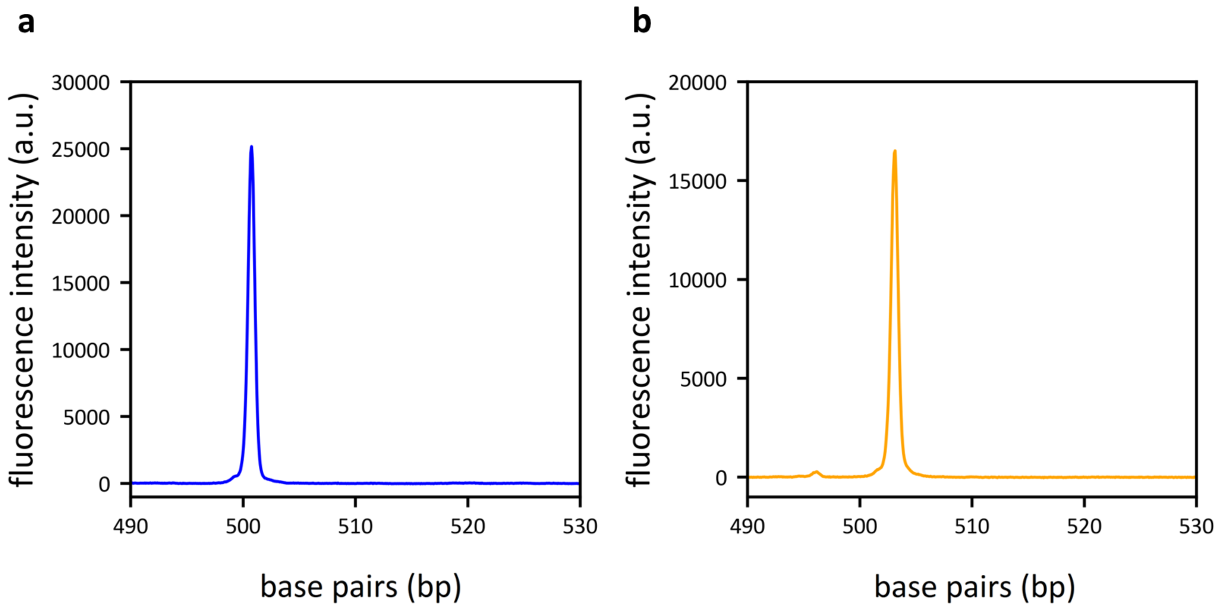


**Figure S21.** Size distribution of GCN repeats in DNA used for *in vitro* transcription. The fragment analysis is presented for **a** 20 and **b** 26 GCN repeats. Fragment analysis was performed using GeneScan 600 LIZ as a size standard.

**Figure S22.**

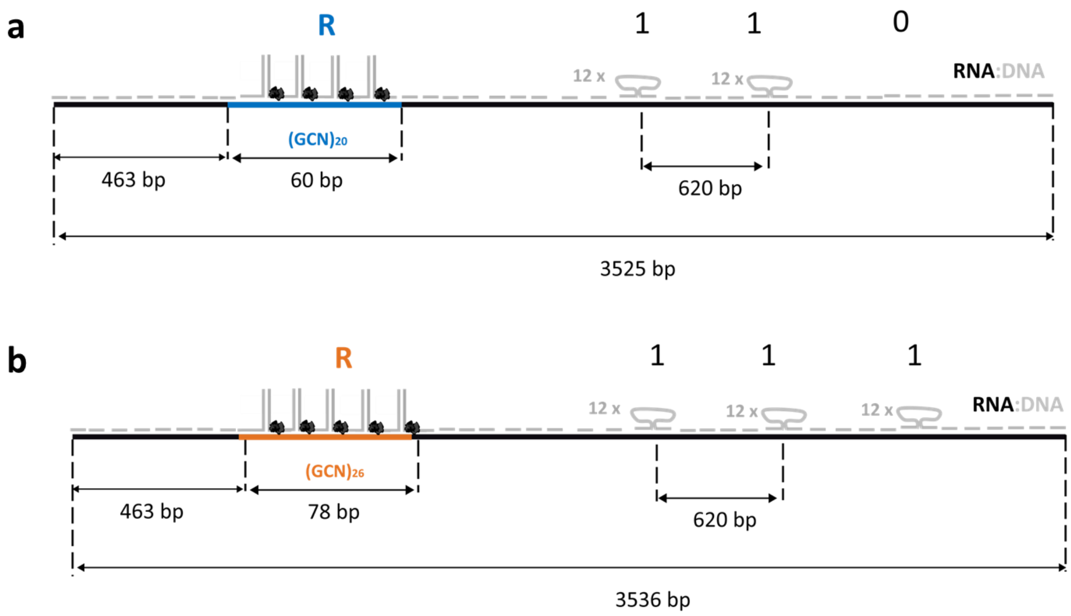


**Figure S22.** Detailed design of RNA:DNA nanostructures used for quantitative analysis of GCN repeats. **a** The RNA:DNA nanostructure tagged with '011' is used to study 20 GCN repeats, **b** RNA:DNA nanostructure tagged with a '111' barcode is used to study 26 GCN repeats in Figure 3.

**Figure S23.**

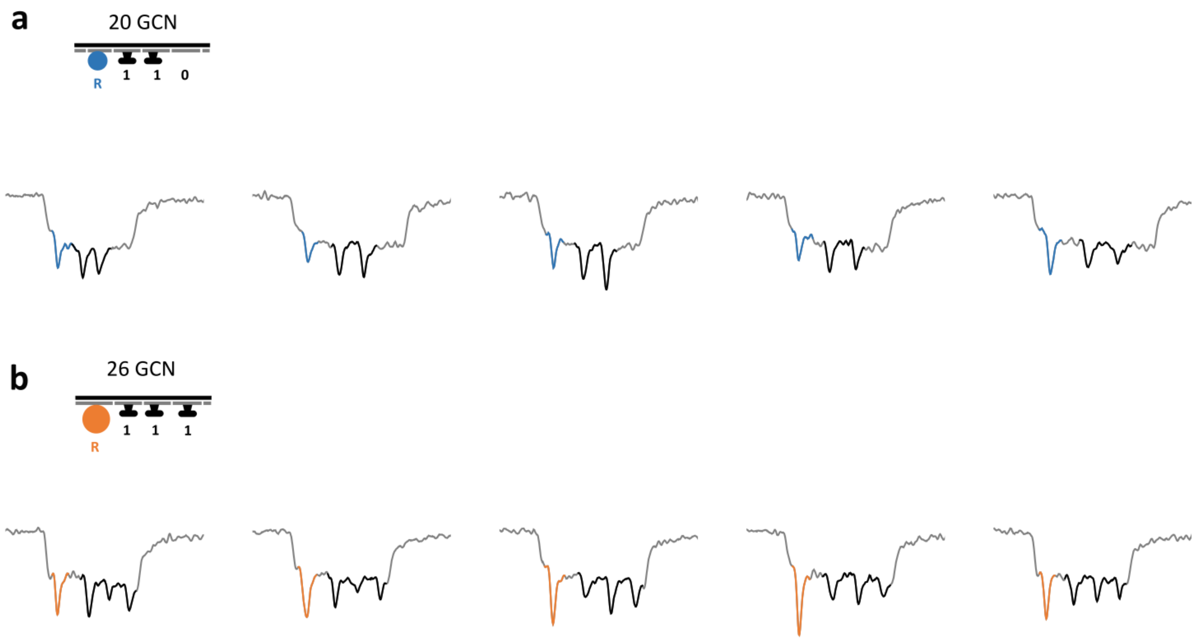


**Figure S23.** Nanopore translocation events of unfolded **a** RNA:DNA nanostructures tagged with ‘011’ barcode, where the 20 GCN repeats are labelled with monovalent streptavidin and **b** RNA:DNA nanostructures tagged with ‘111’ barcode, where the 26 GCN repeats are labelled with monovalent streptavidin.

**Figure S24.**

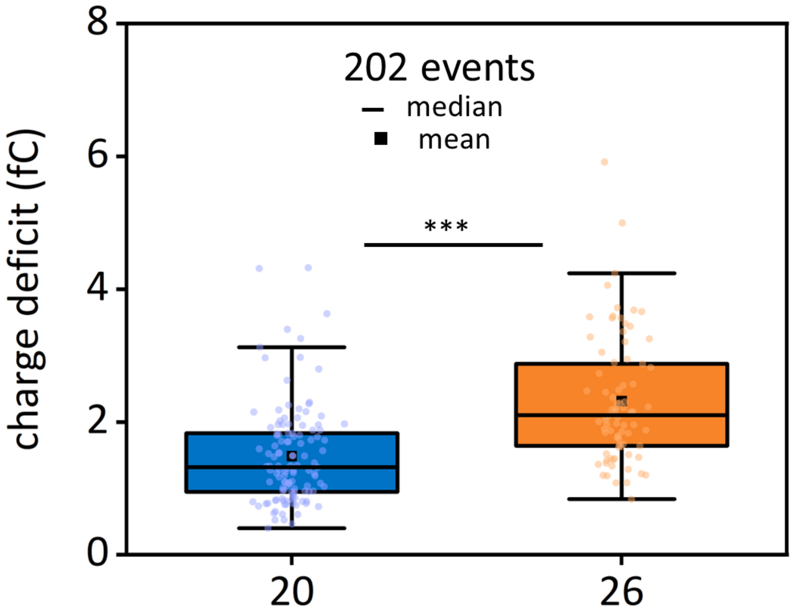


**Figure S24.** The current spike area associated to the repeats labelling shows a higher median for constructs with 26 CGN repeats than 20 GCN repeats (2.2 and 1.7 fC, respectively), and a p-value between the distributions of $7.2\times10^{-10}$ for Welsch t-test. This distinction between the two populations enables clear differentiation of pathogenic repeat expansion sizes, holding clinical relevance for detection of Congenital Central Hypoventilation Syndrome 1.

#### Figure S25.

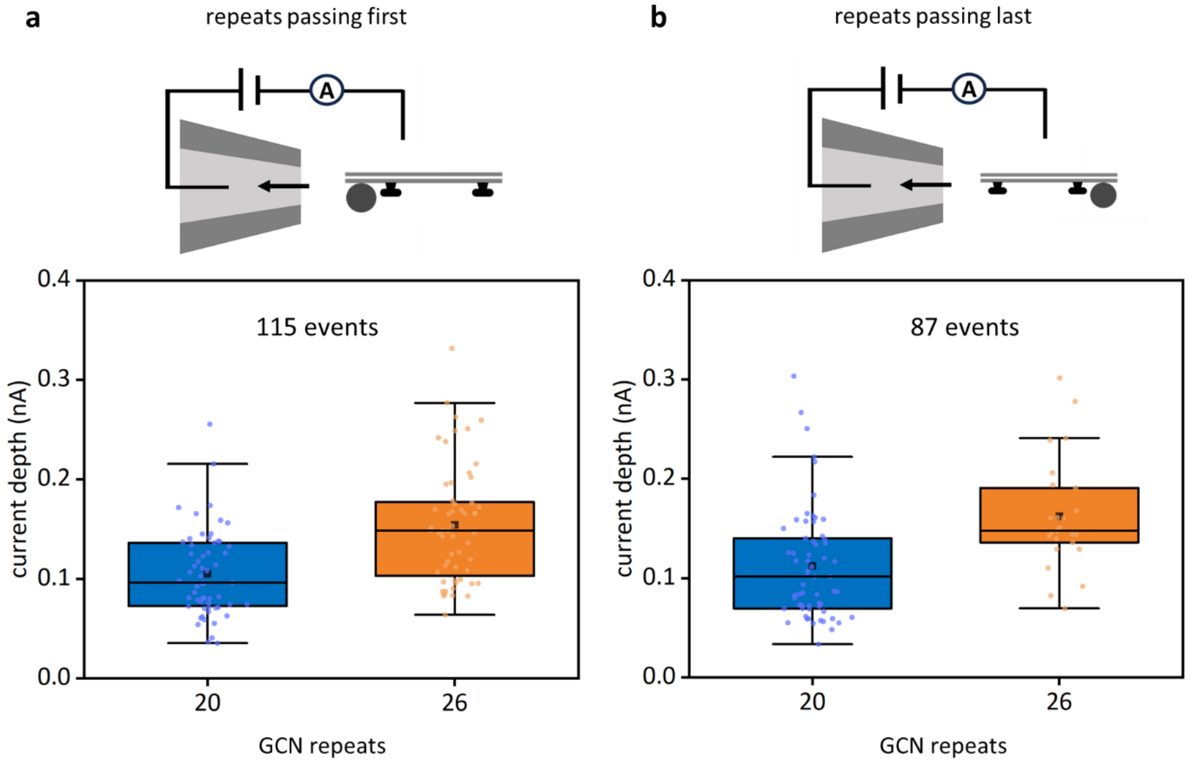


**Figure S25.** GCN repeats can be quantified *via* nanopore sensing independently of the direction of the free translocation. The RNA:DNA nanostructure containing GCN repeats can pass through the nanopore **a** in the 5′ to 3′ direction (repeats translocate first) or **b** in the 3′ to 5′ direction (repeats translocate at the end). The current spike depth ascribed to the repeats increases with the number of repeats independently of the direction of translocation events.

### Figure S26.

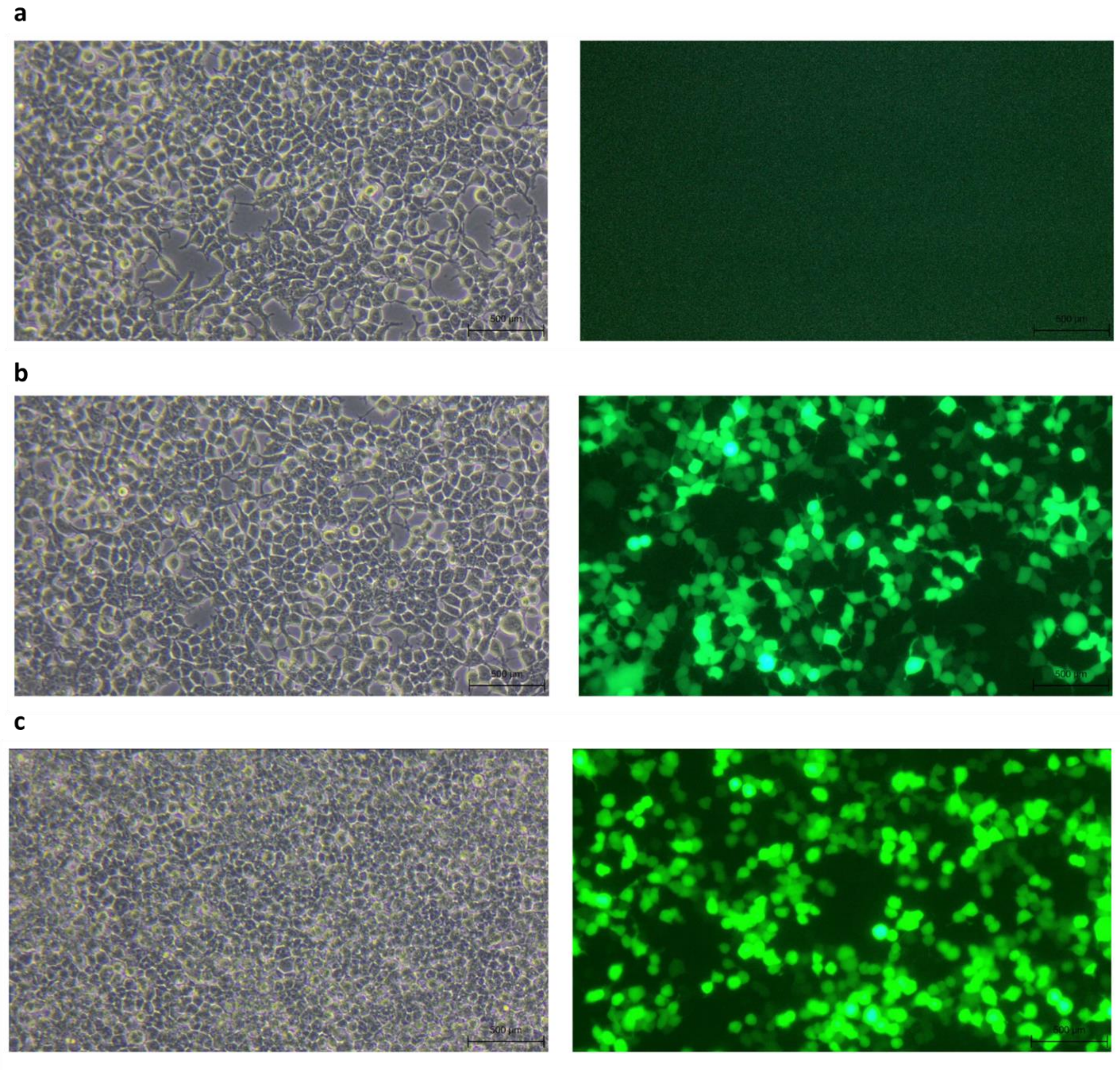


**Figure S26.** Brightfield and fluorescence microscopy images of HEK293 cells from Figure 4b. Cells transfected with **a** lipofectamine only, no plasmid, **b** plasmid coding for EGFP with no repeats, **c** plasmid coding for EGFP and containing 34 CTG tandem repeats. Scale bars are 500 μm.

#### Figure S27.

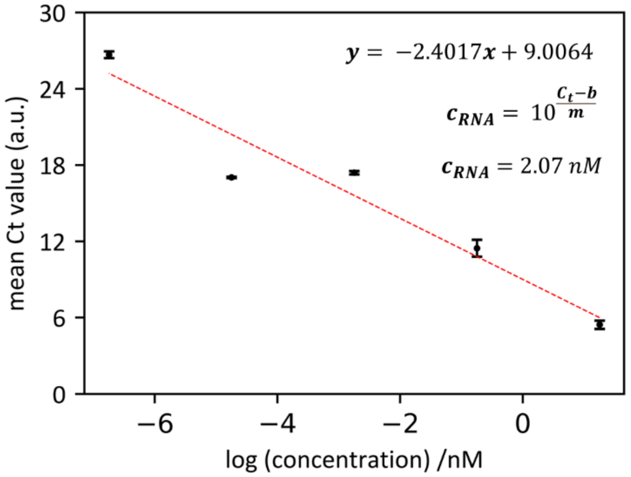


**Figure S27.** Concentration of mRNA with 34 CUG repeats via absolute quantification qPCR. We show the standard curve of pEGPF-N, where we plot the mean $C_t$ value against the logarithm of known concentrations. The error bars correspond to the standard deviation. The concentration of mRNA with 34 CUG repeats ($c_{RNA}$) was derived from the experimental $C_t = 8.2452$. We used the slope ($m$) and intercept ($b$) values of the linear regression performed to the standard curve of pEGPF-N.

**Figure S28.**

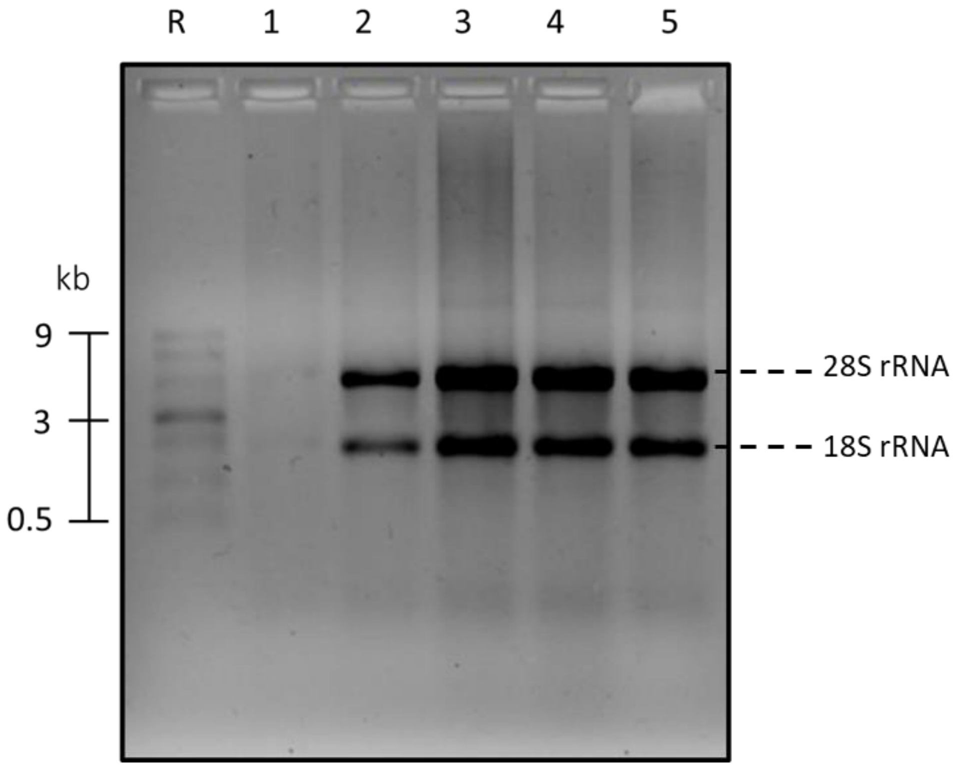


**Figure S28.** Total RNA isolated from HEK293T cells transiently transfected with the expression plasmid pEGFP-N1. Agarose gel electrophoresis shows total RNA containing ~1.3 kb mRNA expressed from pEGFP-N1 plasmid. Lane R: ssRNA ladder (Catalog number N0362S). Lane 1: Blank sample, cells transfected only with lipofectamine, no plasmid (concentration of total RNA: 217 ng/µL). Lane 2: Total RNA with EGFP-coding mRNA, cells transfected with plasmid containing no repeats (concentration of total RNA: 635 ng/µL). Lanes 3-5: Total RNA with EGFP-coding mRNA containing CTG tandem repeats, cells transfected with plasmid containing repeats (concentration of total RNA: 1761 ng/µL, 1239 ng/µL and 959 ng/µL, respectively). Gel: 1 % (w/v) agarose, 1 × TBE.

**Figure S29.**

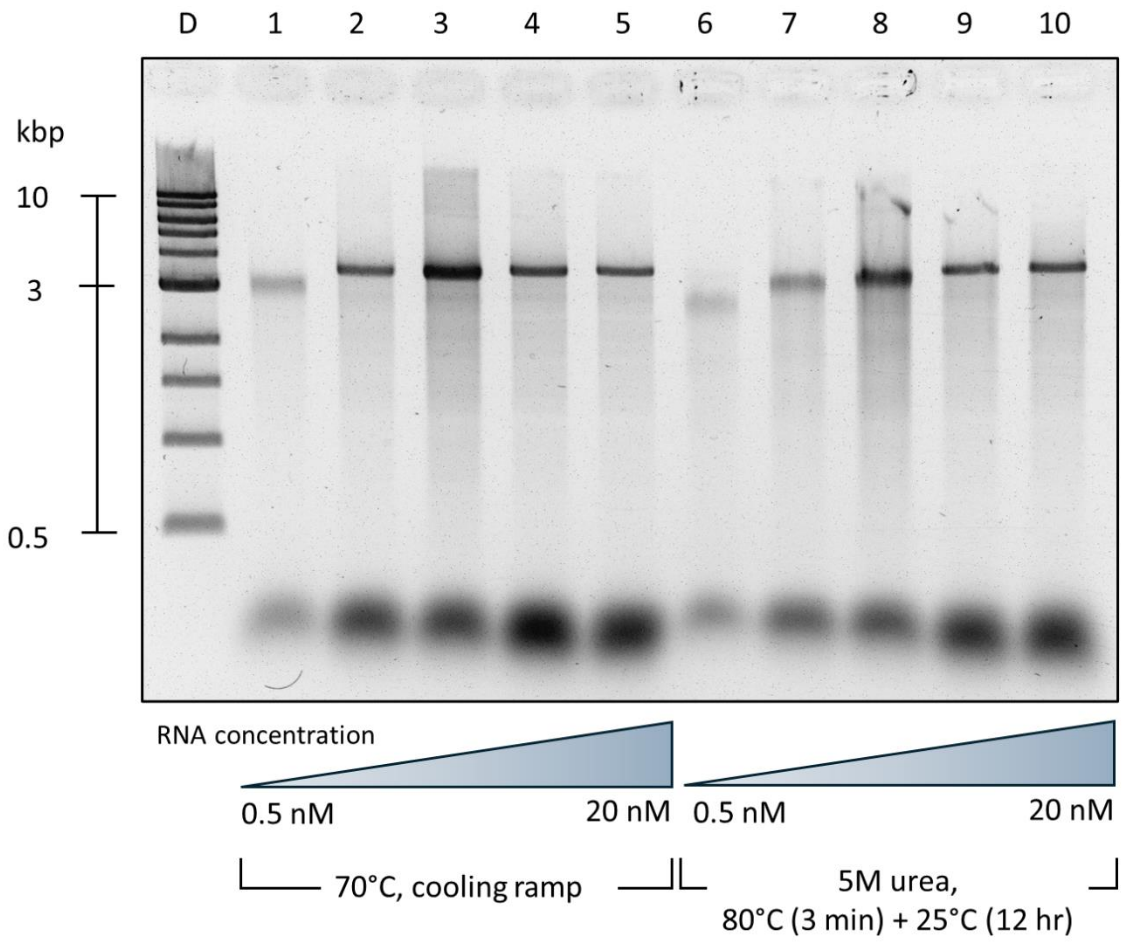


**Figure S29.** Assembly of RNA:DNA nanostructures at varying concentrations. RNA:DNA nanostructures are constructed using 3.6 kb MS2 RNA as a scaffold, with RNA concentrations ranging from 0.5 nM to 20 nM. The DNA oligonucleotides used for duplex assembly are added in 10 times excess for each RNA concentration (sequence of oligonucleotides in Table S15) and two annealing protocols are used. In the first protocol (lanes 1 to 5), the assembly is done by heating to 70°C for 30 seconds, followed by gradual cooling to room temperature over 45 minutes. In the second annealing protocol (lanes 6 to 10), assembly is done in (5M urea), heating at 80°C for 3 minutes followed by incubation at 25°C for 12 hours[2]. For both assembly protocols, it is observed that at low RNA concentration, a 10 times excess of complementary DNA oligonucleotides is insufficient for full RNA:DNA duplex formation. Lane D: 1kbp DNA ladder. Lanes 1 and 6: 0.5 nM RNA:DNA nanostructure. Lanes 2 and 7: 1.5 nM RNA:DNA nanostructure. Lanes 3 and 8: 5 nM RNA:DNA nanostructure. Lanes 4 and 9: 10 nM RNA:DNA nanostructure. Lanes 5 and 10: 20 nM RNA:DNA nanostructure. Gel: 1 % (w/v) agarose, 1 × TBE, 0.02% sodium hypochlorite.

#### Figure S30.

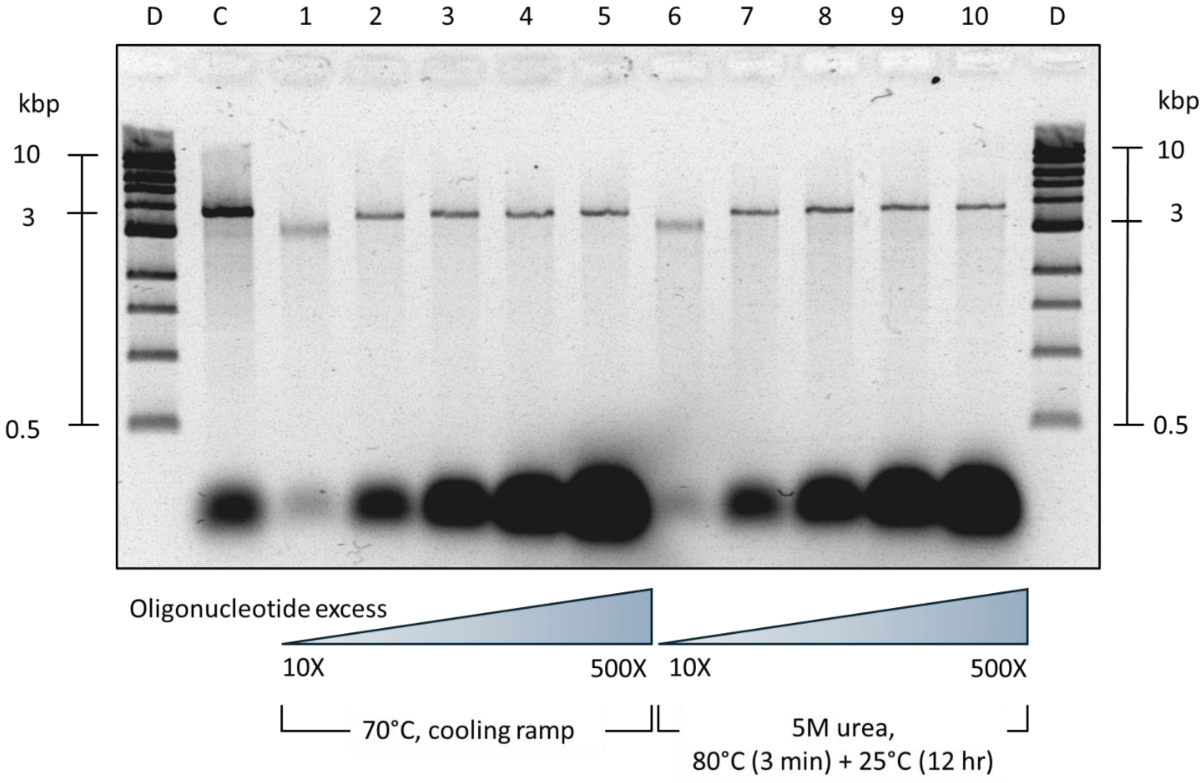


**Figure S30.** Assembly of RNA:DNA nanostructures at low concentration (0.5 nM). RNA:DNA nanostructures are constructed using 3.6 kb MS2 RNA as a scaffold, with RNA concentration of 0.5 nM. The DNA oligonucleotides used for duplex assembly are added in an excess ranging from 10 to 500 times excess and two annealing protocols are used. In the first protocol (lanes 1 to 5), the assembly is done by heating to 70°C for 30 seconds, followed by gradual cooling to room temperature over 45 minutes. In the second annealing protocol (lanes 6 to 10), assembly is done in (5 M urea), heating at 80°C for 3 minutes followed by incubation at 25°C for 12 hours. For both assembly protocols, it is observed that at low RNA concentration (0.5 nM), a 10 times excess of complementary DNA oligonucleotides is insufficient for full RNA:DNA duplex formation. An excess of at least 50 times of DNA oligonucleotides enables RNA:DNA nanostructure assembly at low concentrations. Lane D: 1kbp DNA ladder. Lane C: MS2 control nanostructure assembled at 20 nM concentration with a 10 times DNA excess. Lanes 1 and 6: 0.5 nM RNA:DNA nanostructure with 10 times excess of DNA oligonucleotides. Lanes 2 and 7: 50 times excess. Lanes 3 and 8: 100 times excess. Lanes 4 and 9: 200 times excess. Lanes 5 and 10: 500 times excess. Gel: 1 % (w/v) agarose, 1 × TBE, 0.02% sodium hypochlorite.

#### Figure S31.

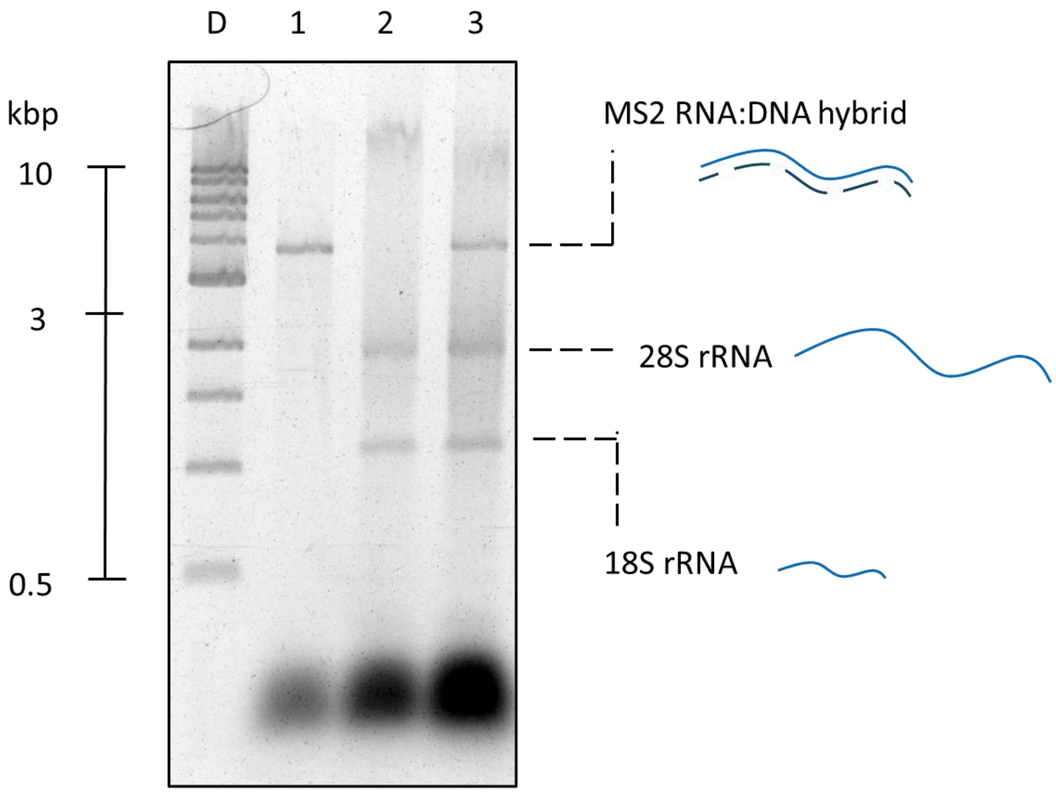


**Figure S31.** Assembly of RNA:DNA nanostructures at low concentration (1 nM) in the presence of total RNA background. Lane D: 1 kbp DNA ladder. Lane 1: MS2 nanostructure assembled at a concentration of 1 nM with 200 times excess of DNA oligonucleotides. As demonstrated previously, this excess is suitable for successful nanostructure assembly. Lane 2: total RNA with DNA oligonucleotides. Lane 3: MS2 nanostructure assembled at a concentration of 1 nM with 200 times excess of DNA oligonucleotides in 100 ng/µL total RNA background, in 100 mM LiCl. Gel: 1 % (w/v) agarose, 1 × TBE, 0.02% sodium hypochlorite.

**Figure S32.**

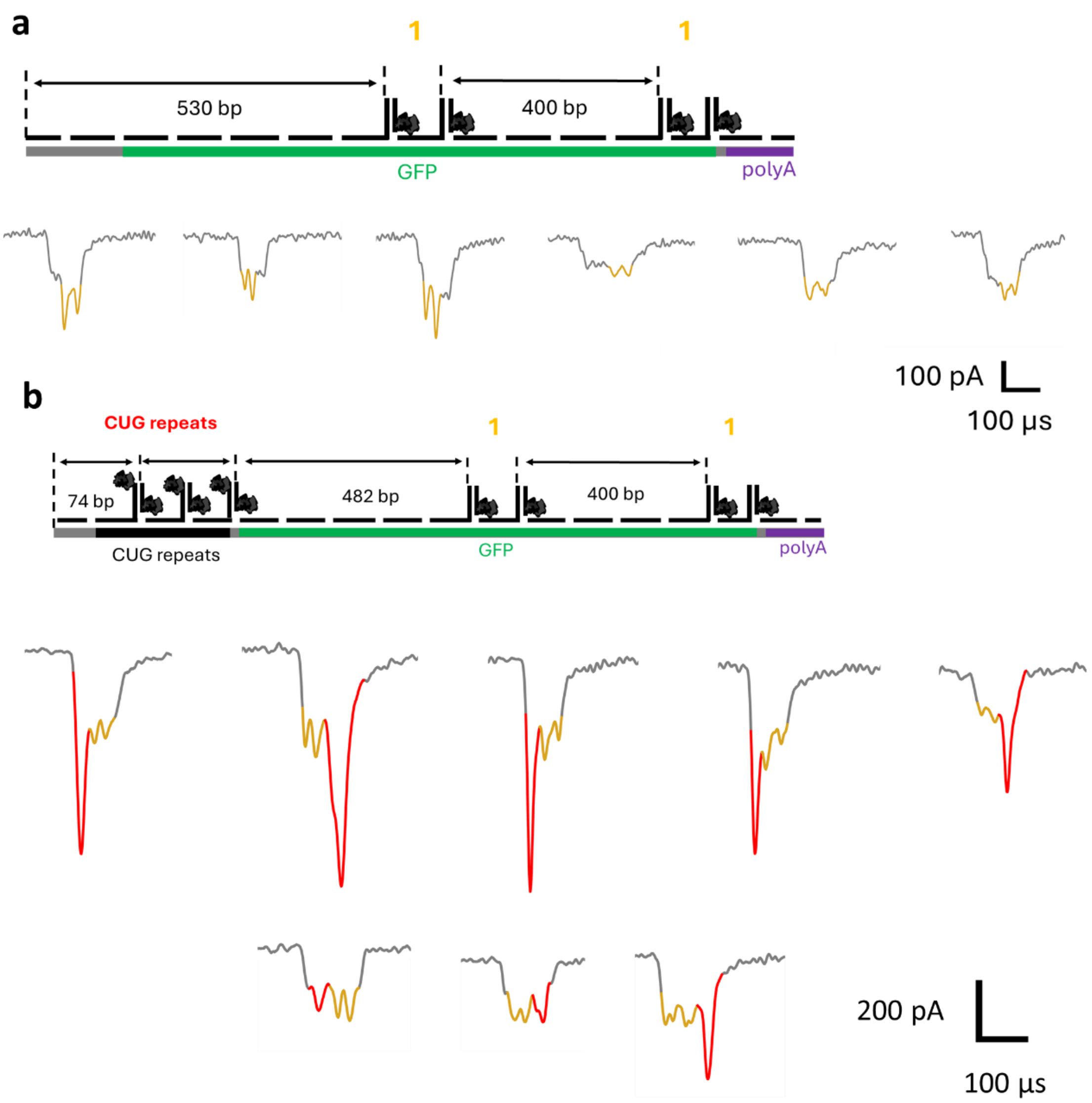


**Figure S32.** Design and nanopore translocation events of unfolded **a** RNA:DNA nanostructures tagged with '11' barcode (gold), where no CUG repeats are present in the RNA transcript used as scaffold. **b** RNA:DNA nanostructures tagged with '11' barcode (gold), where the CUG repeats (red) are labelled with monovalent streptavidin.

**Figure S33.**

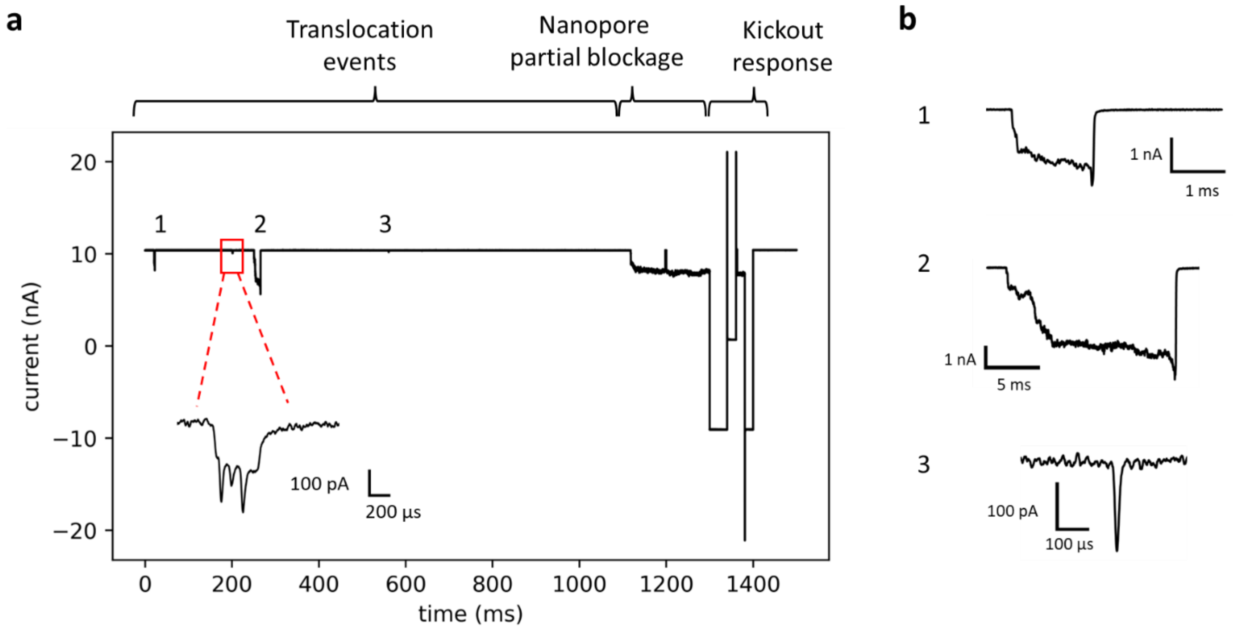


**Figure S33.** MS2 RNA nanostructure detected in total RNA background using nanopore sensing. RNA:DNA nanostructure with a barcode 'PPP' was assembled at 0.5 nM (0.6 ng/µL) concentration in 100 ng/µL total RNA background and sensed using solid state nanopores. **a** The presence of total RNA leads to non-specific translocations and partial blockages of the nanopore. These partial blockages are attributed to RNA adhering to the nanopore surface. This adherence is resolved through a "kickout" response, triggered by an increase in the ionic current noise. The barcode of the nanostructure induces drops in the ionic current with specificity to detect MS2 despite the total RNA background. **b** Non-specific translocation events that produce drops in ionic current are presented.

#### Figure S34.

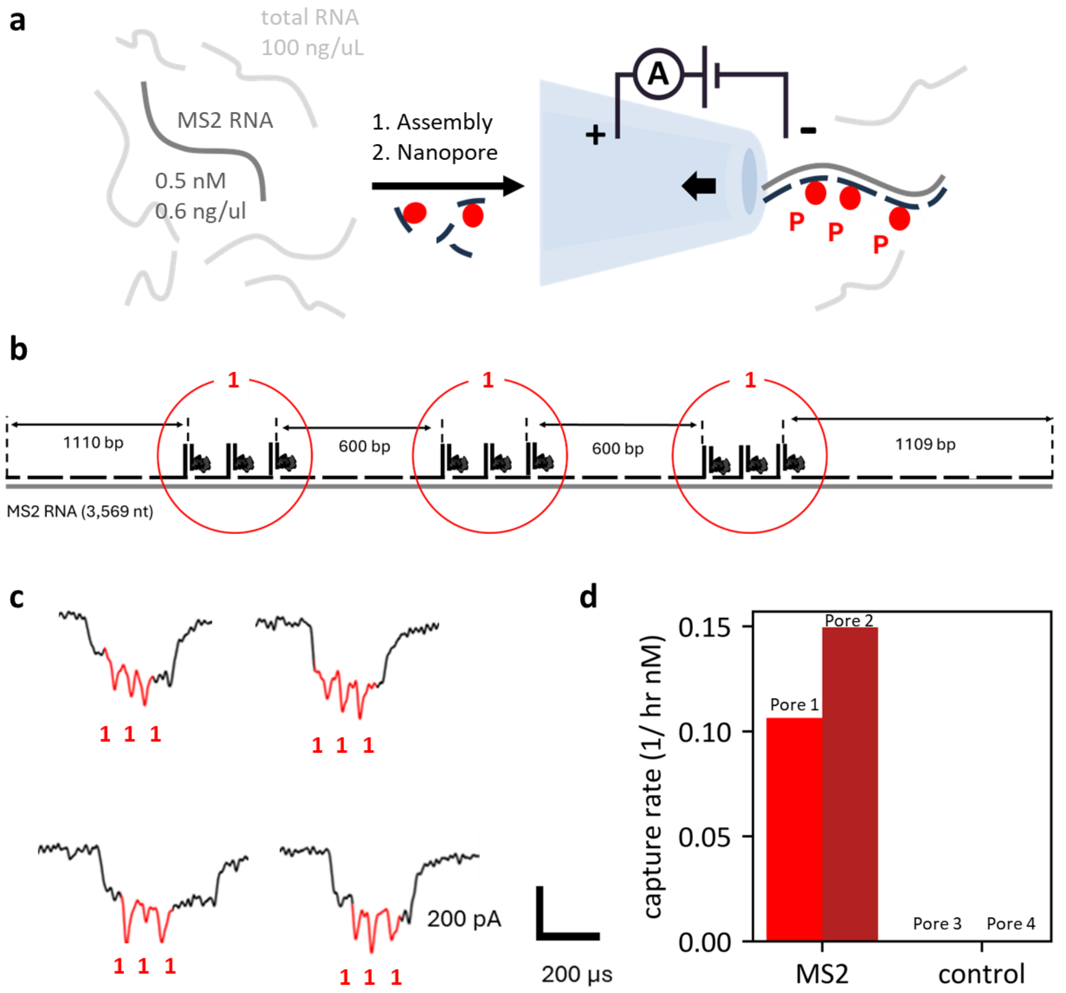


**Figure S34.** Detection of MS2 RNA nanostructures within a total RNA background using nanopore sensing. **a** RNA:DNA nanostructures containing a 'PPP' barcode are assembled using MS2 RNA as scaffold at a concentration of 0.5 nM (0.6 ng/ μL) in a total RNA background of 100 ng/μL and are subsequently detected using solid-state nanopores. **b** The RNA:DNA nanostructure design features a 'PPP' barcode, where each 'P' consists of three consecutive monovalent streptavidins conjugated to biotinylated DNA oligonucleotides. The sequence of the DNA oligonucleotides is presented in Table S16. **c** Nanopore events ascribed to the nanostructure show 3 downward current spikes. **d** Capture rate of 'PPP' MS2 nanostructures agrees with previous reports on the capture rate amplitude of nucleic acids in solid-state nanopores[3,4]. The measurements were performed in the absence of MS2 RNA and DNA oligonucleotides, where we found no translocations that could be ascribed to the nanostructure design, indicating high specificity of the nanostructure produced.

**Figure S35.**

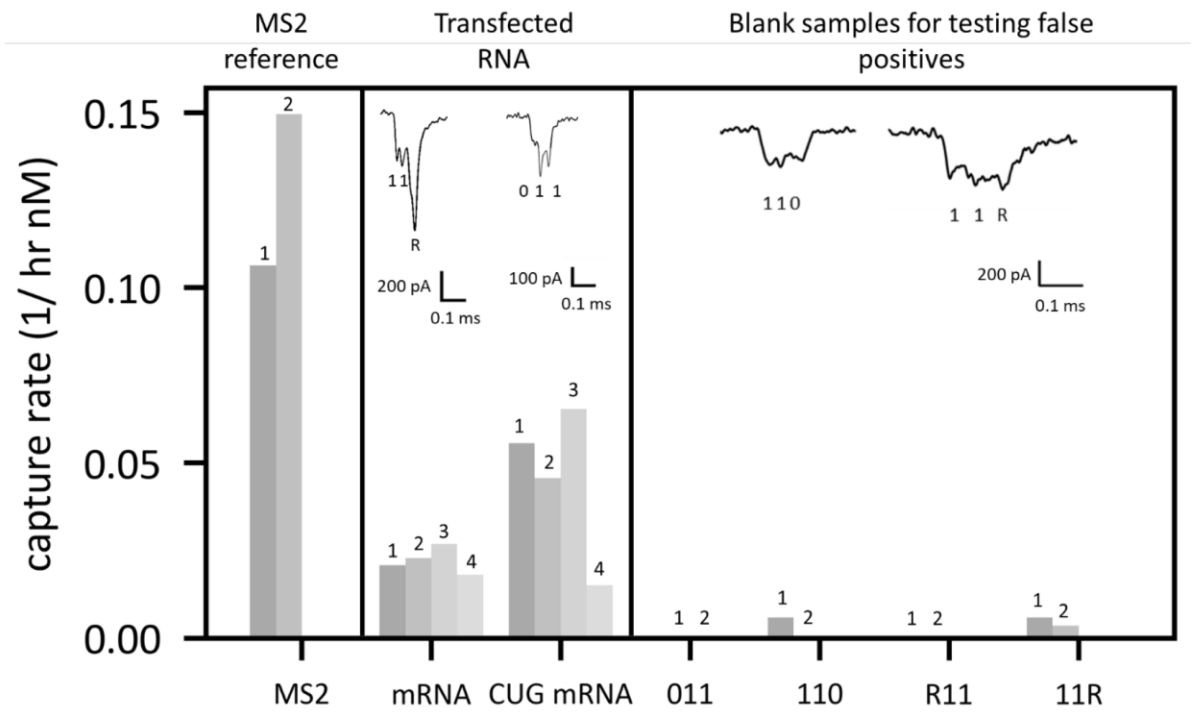


**Figure S35.** MS2, mRNA, and CUG mRNA nanostructures are prepared and detected *via* nanopore sensing in the presence of total RNA background. The capture rate of translocations that can be ascribed to unfolded nanostructures passing through the nanopore is presented for both RNA constructs. The translocation frequency agrees with previous reports on the capture rate amplitude of nucleic acids in solid-state nanopores[3,4], further validating the assembly of the nanostructures and the specificity of nanostructure detection *via* nanopore sensing. In order to detect false positives, nanopore sensing was performed for the mRNA when this was not hybridized with DNA plus oligonucleotides, and for the total RNA where no plasmid transfection was performed. We identified "110" and "11R" translocation events which could be potentially ascribed to the mRNA and CUG mRNA nanostructures, meaning that the event charge deficit, translocation time, and mean current value, was within expected for a 1.2 kb RNA:DNA nanostructure (with and without repeats), and the shape of the current trace matched a similar shape to what would be expected for each RNA:DNA nanostructure design. We attribute these translocations to non-specific molecules from the total RNA background. The capture rate of these random translocations is lower than for the engineered nanostructures, further validating the specificity of the assay.

**Figure S36.**

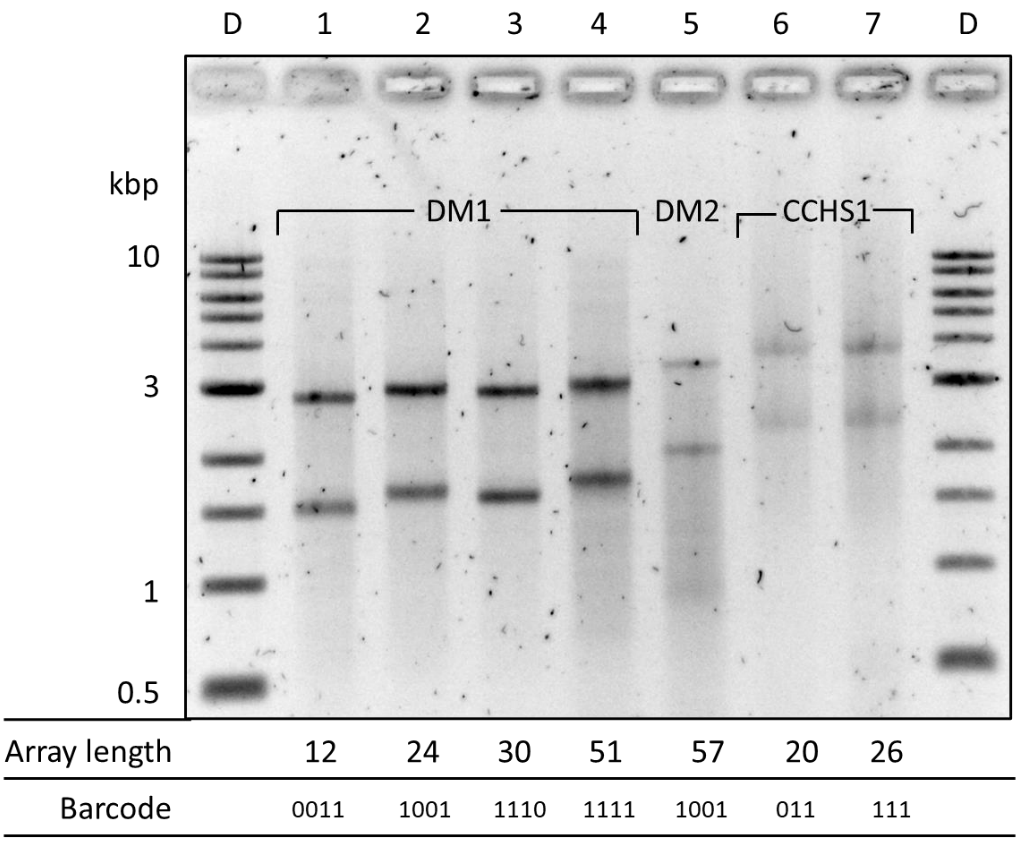


**Figure S36.** Electrophoretic mobility assay (1% agarose gel) of RNA:DNA nanostructures implemented for characterization of tandem repeats. Lanes 'D' correspond to DNA ladders, which are used as reference, ranging from 0.5 to 10 kbp. Lanes 1 to 4 show nanostructures from transcripts holding 12, 24, 30 and 51 CUG repeats, respectively. Each nanostructure holds a four-digit barcode, where '1' corresponds to 12 consecutive DNA dumbbells and '0' is assigned to RNA:DNA heteroduplex. Lane 5 is an RNA:DNA nanostructure from a transcript with 57 CCUG repeats. Lanes 6 and 7 show nanostructures for 20 and 26 GCN repeats, respectively. Each nanostructure holds a three-digit barcode, where '1' corresponds to 12 consecutive DNA dumbbells and '0' is assigned to RNA:DNA heteroduplex. For each nanostructure, two bands are depicted. The upper band corresponds to the full-length RNA:DNA nanostructure and the lower band originates from the nanostructure of prematurely terminated transcripts.

**Figure S37.**

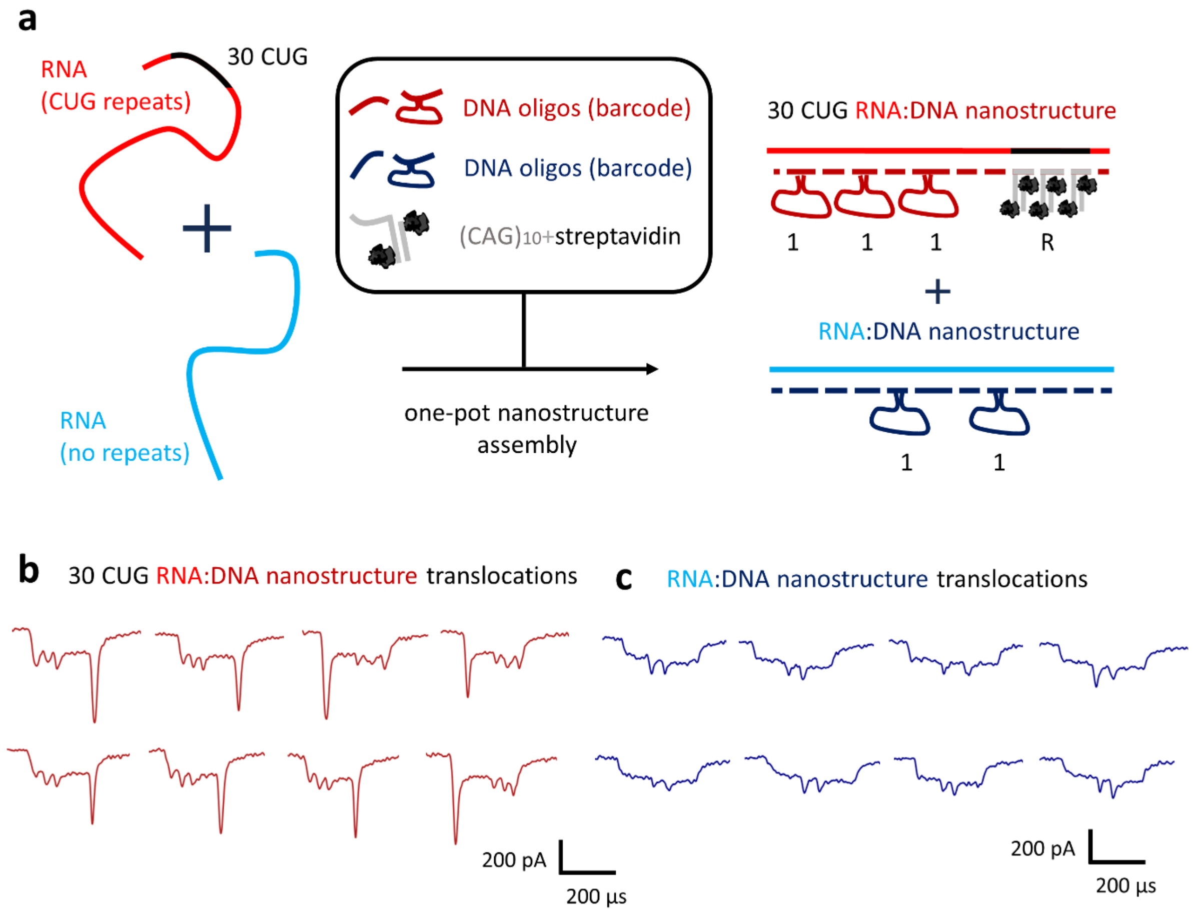


**Figure S37.** Multiplexed assembly and nanopore characterization of RNA:DNA nanostructures. **a** A 3 kb RNA containing 30 CUG repeats (red) was mixed with a 3.6 kb RNA containing no tandem repeats (blue) in a single reaction tube. DNA oligonucleotides were added to label the CUG-containing RNA with a '111' barcode and the repeat-free RNA with a '11' barcode. Biotinylated $(CAG)_{10}$ oligonucleotides were also included to hybridize to the CUG repeats. **b** Nanopore characterization revealed translocations with current spikes attributed to the '111' barcode and an additional spike associated with the CUG repeat region. **c** Nanopore readout using the same nanopore also showed translocations featuring the '11' barcode, corresponding to the transcript without repeats. These results demonstrate multiplexed assembly and nanopore detection of distinct RNA:DNA nanostructures in a single mixture.

**Figure S38.**

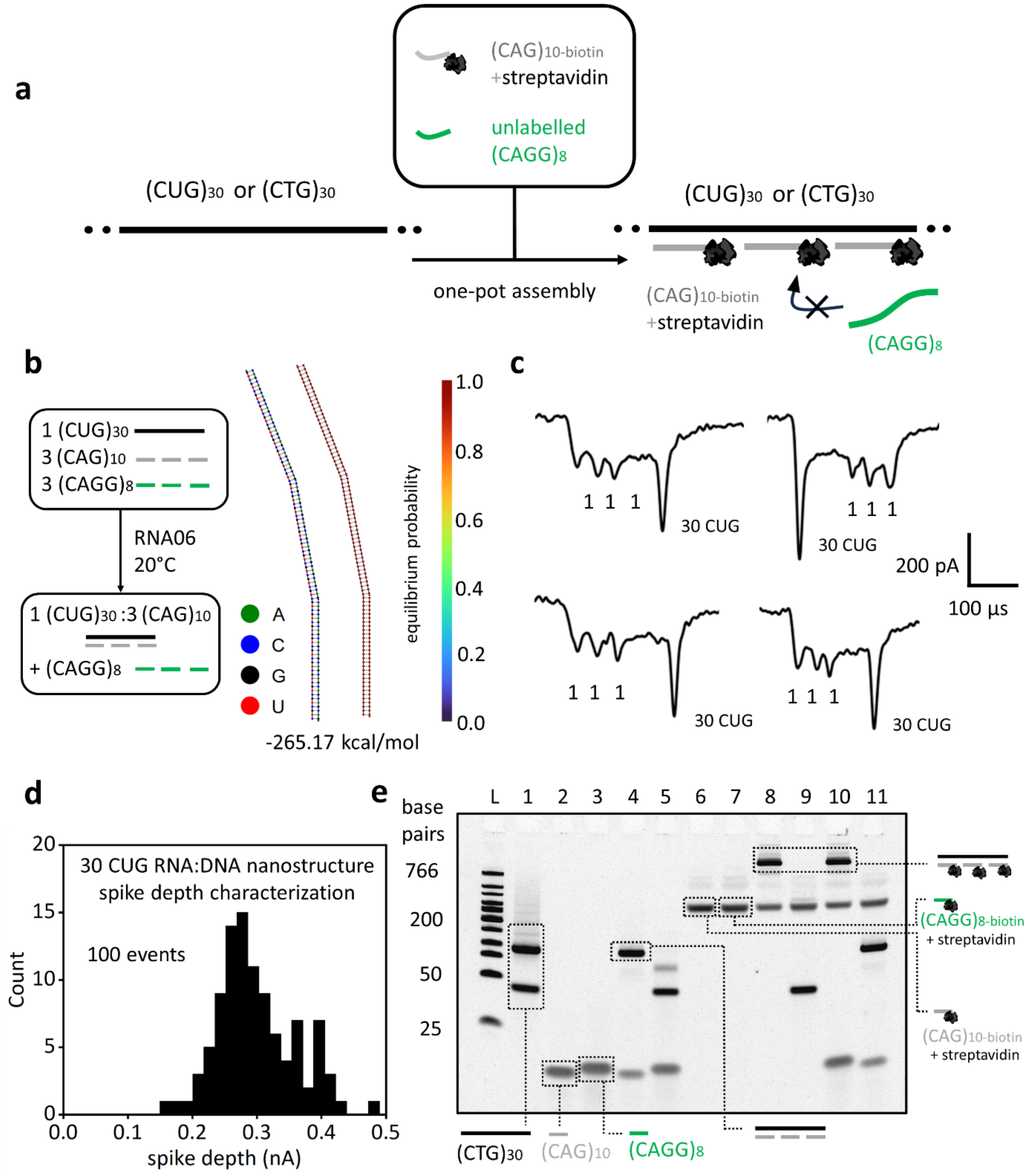


**Figure S38.** CAG oligonucleotides selectively hybridize to CUG repeats outcompeting CAGG oligonucleotides. **a** We evaluated potential cross-hybridization by testing binding of $(CAG)_{10}$ and $(CAGG)_8$ oligonucleotides to $(CUG)_{30}$ repeats in full-length transcripts with nanopore sensing. We also test hybridization of the oligonucleotides to $(CTG)_{30}$ repeats with native polyacrylamide gel electrophoresis. **b** NUPACK3 with the RNA06 model (including full stacking parameterization) was used to predict complexes in a mixture of 1:3:3 µM =

$(CUG)_{30}$:$(CAG)_{10}$:$(CAGG)_8$ at 20°C. The model predicts 100% of $(CUG)_{30}$ binding to three equivalents of $(CAG)_{10}$. The free energy of the secondary structure is -265.17 kcal/mol. No $(CAGG)_8$ cross-hybridization with $(CUG)_{30}$ is predicted. **c** An RNA:DNA nanostructure was assembled from the transcript containing 30 CUG repeats in the presence of biotinylated $(CAG)_{10}$ oligonucleotides, which recruit streptavidin, together with non-biotinylated $(CAGG)_8$ oligonucleotides, which do not bind streptavidin. Nanopore recordings show translocations with the '111' barcode spikes and an additional spike corresponding to the repeat region. **d** The spike-depth distribution attributed to the repeats indicates streptavidin binding mediated by hybridization of biotinylated $(CAG)_{10}$ oligonucleotides, demonstrating that $(CAG)_{10}$ oligonucleotides outcompete $(CAGG)_8$ oligonucleotides and prevent repeat mislabelling. **e** PAGE further supports selective $(CAG)_{10}$ hybridization specificity outcompeting $(CAGG)_8$ oligonucleotides to $(CUG)_{30}$ repeats. Lane L: Low range DNA ladder. Lane 1: $(CTG)_{30}$. Lane 2: $(CAG)_{10}$. Lane 3: $(CAGG)_8$. Lane 4: $(CTG)_{30}$ + $(CAG)_{10}$. Lane 5: $(CTG)_{30}$ + $(CAGG)_8$. Lane 6: $(CAG)_{10\text{-biotin}}$ + streptavidin. Lane 7: $(CAGG)_{8\text{-biotin}}$ + streptavidin. Lane 8: $(CTG)_{30}$ + $(CAG)_{10\text{-biotin}}$ + streptavidin. Lane 9: $(CTG)_{30}$ + $(CAGG)_{8\text{-biotin}}$ + streptavidin. Lane 10: $(CTG)_{30}$ + $(CAG)_{10\text{-biotin}}$ + $(CAGG)_8$ + streptavidin. Lane 11: $(CTG)_{30}$ + $(CAG)_{10}$ + $(CAGG)_{8\text{-biotin}}$ + streptavidin.

**Figure S39.**

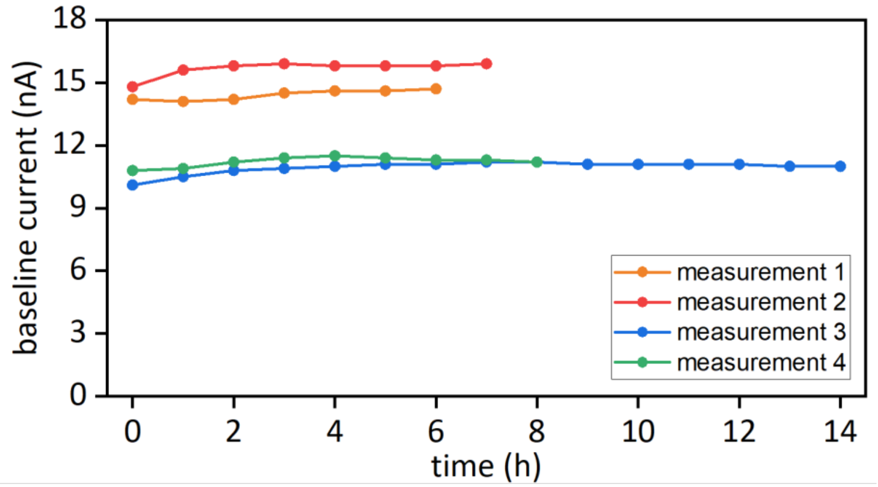


**Figure S39.** Ionic current baseline for nanopore measurements in 4M LiCl with durations of 6, 7, 8 and 14 hours.

### Figure S40.

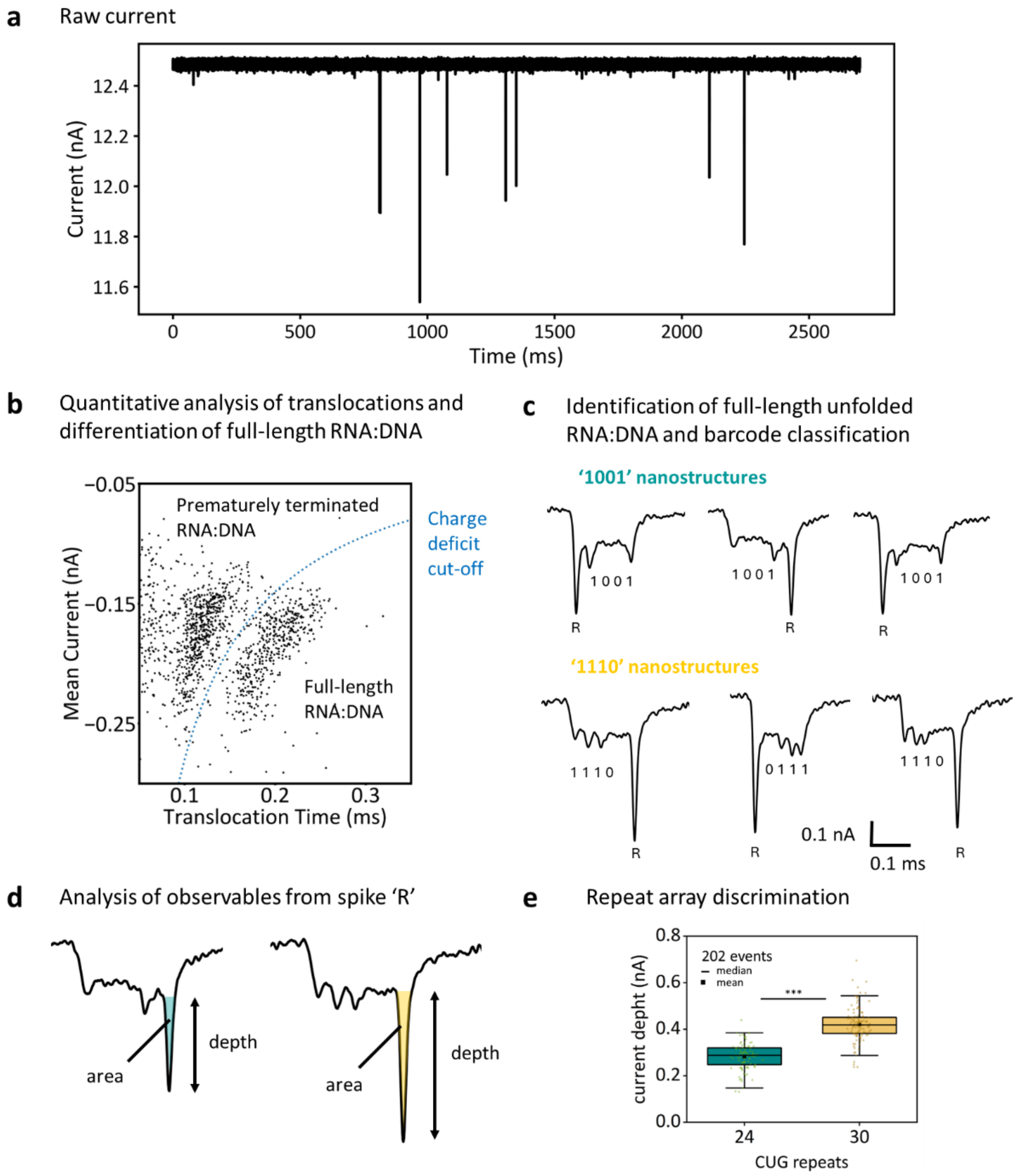


**Figure S40.** Step-by-step nanopore data analysis of tandem repeat arrays. **a** An example segment of the raw ionic current trace is shown for multiplexed characterization of transcripts containing 24 and 30 tandem repeats. Translocation events are identified using custom LabVIEW software based on simple thresholding. Typical parameters include a translocation duration >0.05 ms, a charge deficit of 10–400 fC, and a mean event current below −100 pA.

These criteria enable reliable detection of events corresponding to RNA:DNA nanostructures. **b** Further quantitative analysis of event features (translocation time, charge deficit, mean current) is used to distinguish full-length nanostructures from prematurely terminated or fragmented constructs. **c** Using the nanostructure design and translocation time, unfolded constructs are classified by barcode. In this example, the 24-repeat nanostructure carries a '1001' barcode, whereas the 30-repeat nanostructure carries a '1110' barcode. **d** Finally, the spike depth and area associated with the tandem repeat region are quantified, **e** enabling discrimination of repeat array length.

#### Figure S41.

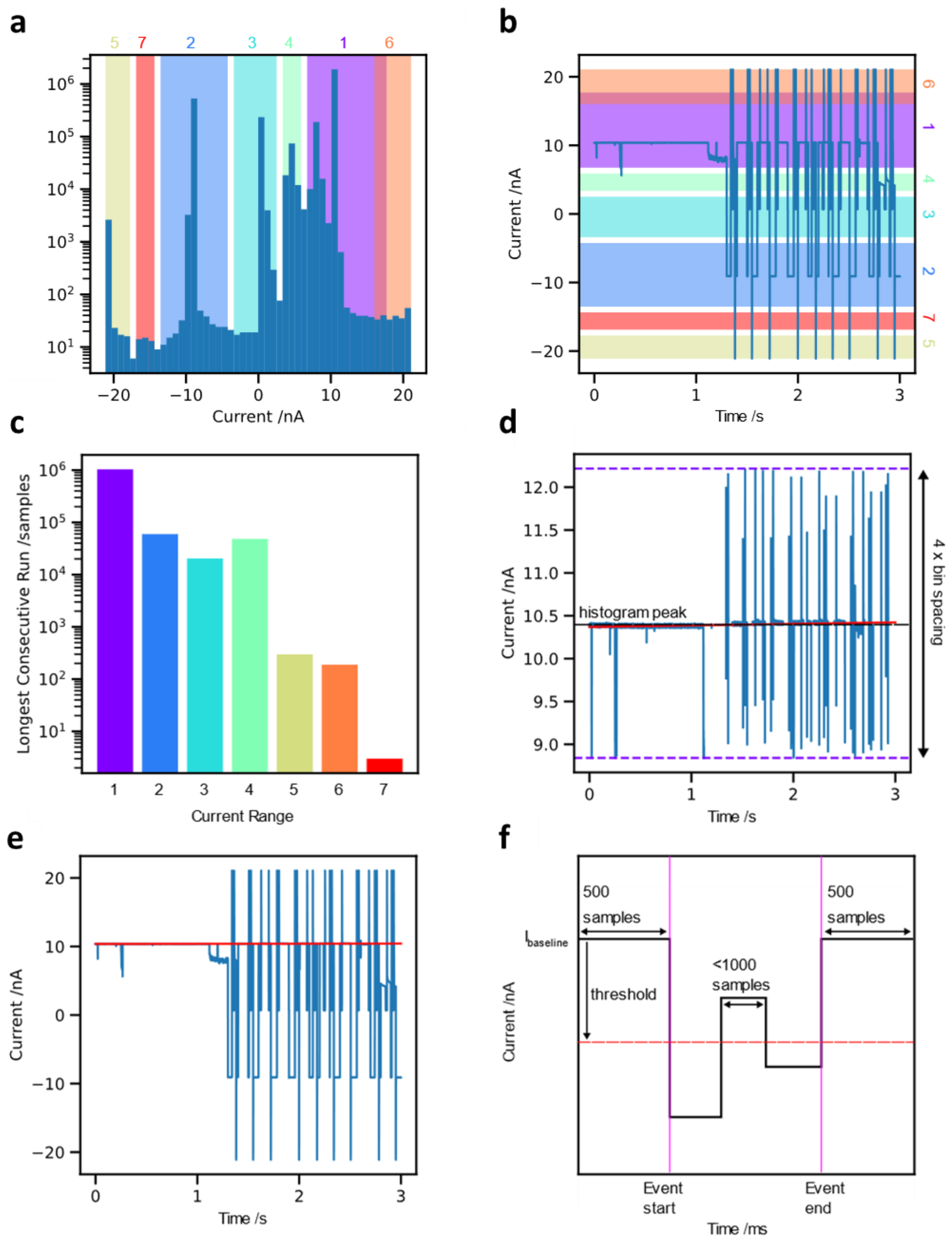


**Figure S41.** Extraction of mRNA nanostructure translocation events from ionic current trace. **a** A histogram is generated with all current values from the file. Peaks are detected and peak boundaries found. **b** The peak boundaries from the histogram displayed overlayed on the raw current trace. **c** For each pair of boundaries found from the histogram, the longest consecutive run of samples within the range is found. The current range with the longest consecutive run is used to calculate the baseline. **d** Points more than 2 histogram bin widths away from the selected peak are removed and a line is fitted to the remaining data to approximate the baseline. **e** The full current trace with the calculated baseline overlayed. **f** An event is extract by finding a start point where the current drops from the baseline by more than the threshold value. The end of the event is determined by finding the first time when the blockade drops below the threshold value and does not recover within 1000 samples. A berth of 500 points either side of the event start and end are included when the event is saved.

#### Table S1.

Sequence of modified pJET1.2/blunt cloning vector (CloneJET PCR Cloning Kit, Thermo Fisher, Catalog number: K1231): Circular DNA (3179 bp) with inserted $(CTG)_{51}$ tandem repeats. Map of the plasmid is included at the end of the table.

| Sequence (5′→ 3′) |
|---|
| T7 promoter $(CTG)_{51}$ repeats DraIII Cutting |
| GCCCCTGCAGCCGAATTATATTATTTTTGCCAAATAATTTTTAACAAAAGCTCTGAAGTCTTCTTC<br>ATTTAAATTCTTAGATGATACTTCATCTGGAAAATTGTCCCAATTAGTAGCAT**CACGCTGTG**AGTA<br>AGTTCTAAACCATTTTTTTATTGTTGTATTATCTCTAATCTTACTACTCGATGAGTTTTCGGTATT<br>ATCTCTATTTTTAACTTGGAGCAGGTTCCATTCATTGTTTTTTTCATCATAGTGAATAAAATCAAC<br>TGCTTTAACACTTGTGCCTGAACACCATATCCATCCGGCG**TAATACGACTCACTATAGG**GAGAGCG<br>GCCGCCAGATCTTCCGGATGGCTCGAGTTTTTCAGCAAGATCATGGTGCAGTGTAGCCGGGAATG**C**<br>**TGCTGCTGCTGCTGCTGCTGCTGCTGCTGCTGCTGCTGCTGCTGCTGCTGCTGCTGCTGCTGCTGC**<br>**TGCTGCTGCTGCTGCTGCTGCTGCTGCTGCTGCTGCTGCTGCTGCTGCTGCTGCTGCTGCTGCTGC**<br>**TGCTGCTGCTGCTGCTGCTG**GGGGGATCACAGACCATTCTCGGCTTAAATCTTTCTAGAAGATCTC<br>CTACAATATTCTCAGCTGCCATGGAAAATCGATGTTCTTCTTTTATTCTCTCAAGATTTTCAGGCT<br>GTATATTAAAACTTATATTAAGAACTATGCTAACCACCTCATCAGGAACCGTTGTAGGTGGCGTGG<br>GTTTTCTTGGCAATCGACTCTCATGAAAACTACGAGCTAAATATTCAATATGTTCCTCTTGACCAA<br>CTTTATTCTGCATTTTTTTTGAACGAGGTTTAGAGCAAGCTTCAGGAAACTGAGACAGGAATTTTA<br>TTAAAAATTTAAATTTTGAAGAAAGTTCAGGGTTAATAGCATCCATTTTTTGCTTTGCAAGTTCCT<br>CAGCATTCTTAACAAAAGACGTCTCTTTTGACATGTTTAAAGTTTAAACCTCCTGTGTGAAATTAT<br>TATCCGCTCATAATTCCACACATTATACGAGCCGGAAGCATAAAGTGTAAAGCCTGGGGTGCCTAA<br>TGAGTGAGCTAACTCACATTAATTGCGTTGCGCTCACTGCCAATTGCTTTCCAGTCGGGAAACCTG<br>TCGTGCCAGCTGCATTAATGAATCGGCCAACGCGCGGGGAGAGGCGGTTTGCGTATTGGGCGCTCT<br>TCCGCTTCCTCGCTCACTGACTCGCTGCGCTCGGTCGTTCGGCTGCGGCGAGCGGTATCAGCTCAC<br>TCAAAGGCGGTAATACGGTTATCCACAGAATCAGGGGATAACGCAGGAAAGAACATGTGAGCAAAA<br>GGCCAGCAAAAGGCCAGGAACCGTAAAAAGGCCGCGTTGCTGGCGTTTTTCCATAGGCTCCGCCCC<br>CCTGACGAGCATCACAAAAATCGACGCTCAAGTCAGAGGTGGCGAAACCCGACAGGACTATAAAGA<br>TACCAGGCGTTTCCCCCTGGAAGCTCCCTCGTGCGCTCTCCTGTTCCGACCCTGCCGCTTACCGGA<br>TACCTGTCCGCCTTTCTCCCTTCGGGAAGCGTGGCGCTTTCTCATAGCTCACGCTGTAGGTATCTC<br>AGTTCGGTGTAGGTCGTTCGCTCCAAGCTGGGCTGTGTGCACGAACCCCCCGTTCAGCCCGACCGC<br>TGCGCCTTATCCGGTAACTATCGTCTTGAGTCCAACCCGGTAAGACACGACTTATCGCCACTGGCA<br>GCAGCCACTGGTAACAGGATTAGCAGAGCGAGGTATGTAGGCGGTGCTACAGAGTTCTTGAAGTGG<br>TGGCCTAACTACGGCTACACTAGAAGGACAGTATTTGGTATCTGCGCTCTGCTGAAGCCAGTTACC<br>TTCGGAAAAAGAGTTGGTAGCTCTTGATCCGGCAAACAAACCACCGCTGGTAGCGGTGGTTTTTTT<br>GTTTGCAAGCAGCAGATTACGCGCAGAAAAAAAGGATCTCAAGAAGATCCTTTGATCTTTTCTACG<br>GGGTCTGACGCTCAGTGGAACGAAAACTCACGTTAAGGGATTTTGGTCATGAGATTATCAAAAAGG<br>ATCTTCACCTAGATCCTTTTAAATTAAAAATGAAGTTTTAAATCAATCTAAAGTATATATGAGTAA<br>ACTTGGTCTGACAGTTACCAATGCTTAATCAGTGAGGCACCTATCTCAGCGATCTGTCTATTTCGT<br>TCATCCATAGTTGCCTGACTCCCCGTCGTGTAGATAACTACGATACGGGAGGGCTTACCATCTGGC<br>CCCAGTGCTGCAATGATACCGCGAGACCCACGCTCACCGGCTCCAGATTTATCAGCAATAAACCAG<br>CCAGCCGGAAGGGCCGAGCGCAGAAGTGGTCCTGCAACTTTATCCGCCTCCATCCAGTCTATTAAT |

```
TGTTGCCGGGAAGCTAGAGTAAGTAGTTCGCCAGTTAATAGTTTGCGCAACGTTGTTGCCATTGCT
ACAGGCATCGTGGTGTCACGCTCGTCGTTTGGTATGGCTTCATTCAGCTCCGGTTCCCAACGATCA
AGGCGAGTTACATGATCCCCCATGTTGTGCAAAAAAGCGGTTAGCTCCTTCGGTCCTCCGATCGTT
GTCAGAAGTAAGTTGGCCGCAGTGTTATCACTCATGGTTATGGCAGCACTGCATAATTCTCTTACT
GTCATGCCATCCGTAAGATGCTTTTCTGTGACTGGTGAGTACTCAACCAAGTCATTCTGAGAATAG
TGTATGCGGCGACCGAGTTGCTCTTGCCCGGCGTCAATACGGGATAATACCGCGCCACATAGCAGA
ACTTTAAAAGTGCTCATCATTGGAAAACGTTCTTCGGGGCGAAAACTCTCAAGGATCTTACCGCTG
TTGAGATCCAGTTCGATGTAACCCACTCGTGCACCCAACTGATCTTCAGCATCTTTTACTTTCACC
AGCGTTTCTGGGTGAGCAAAAACAGGAAGGCAAAATGCCGCAAAAAAGGGAATAAGGGCGACACGG
AAATGTTGAATACTCATACTCTTCCTTTTTCAATATTATTGAAGCATTTATCAGGGTTATTGTCTC
ATGAGCGGATACATATTTGAATGTATTTAGAAAAATAAACAAATAGGGGTTCCGCGCACATTTCCC
CGAAAAGTGCCACCTGACGTCTAAGAAACCATTATTATCATGACATTAACCTATAAAAATAGGCGT
ATCACGAGGCC
```

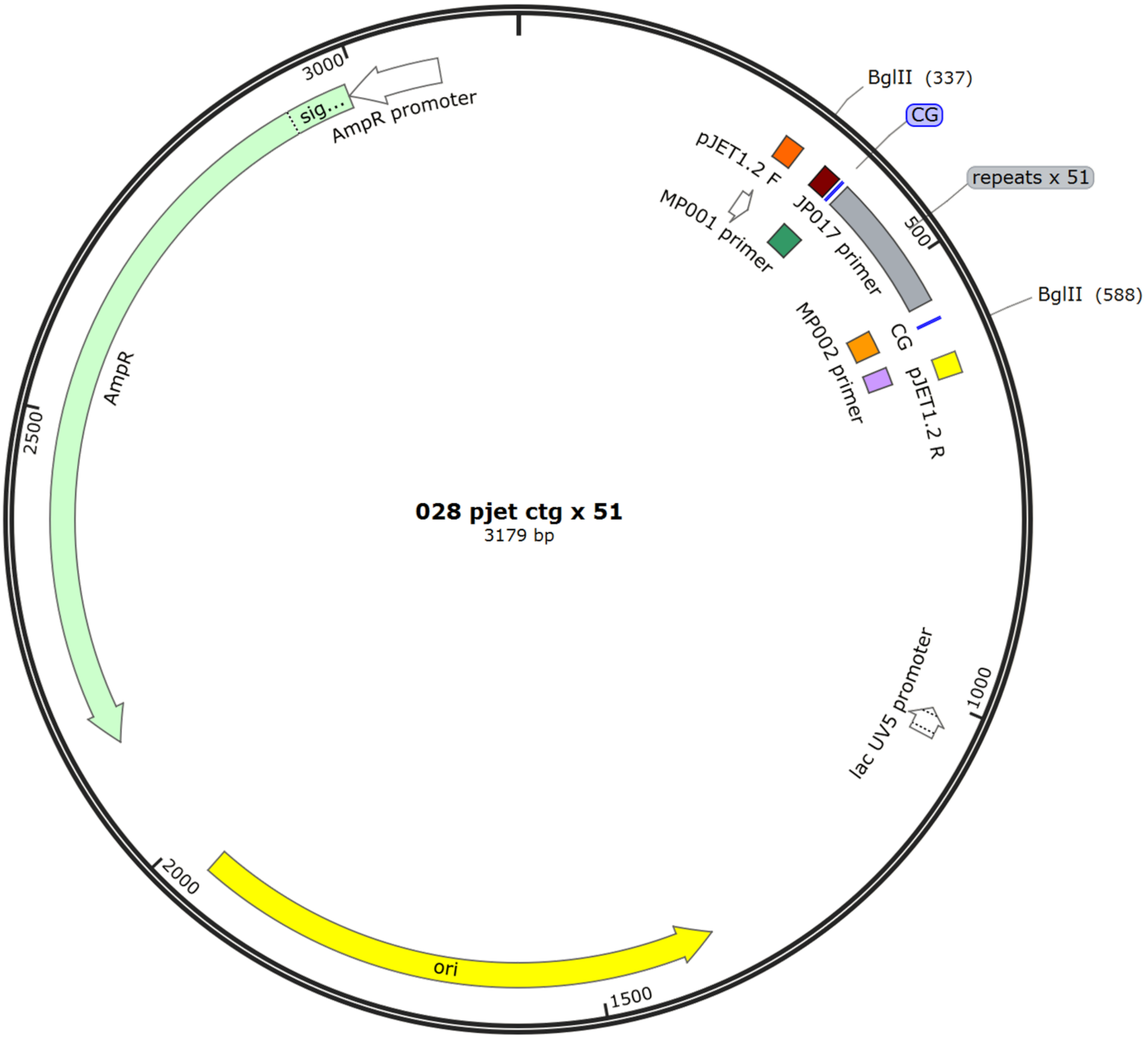

#### Table S2.

Sequence of modified pJET1.2/blunt cloning vector (CloneJET PCR Cloning Kit, Thermo Fisher, Catalog number: K1231): Circular DNA (3398 bp) with inserted $(CCTG)_{57}$ tandem repeats.

| Sequence (5′→ 3′) |
|---|
| **T7 promoter** **$(CCTG)_{57}$ repeats** **DraIII Cutting** |

GCCCCTGCAGCCGAATTATATTATTTTTGCCAAATAATTTTTAACAAAAGCTCTGAAGTCTTCTTC
ATTTAAATTCTTAGATGATACTTCATCTGGAAAATTGTCCCAATTAGTAGCAT**CACGCTGTG**AGTA
AGTTCTAAACCATTTTTTTATTGTTGTATTATCTCTAATCTTACTACTCGATGAGTTTTCGGTATT
ATCTCTATTTTTAACTTGGAGCAGGTTCCATTCATTGTTTTTTTCATCATAGTGAATAAAATCAAC
TGCTTTAACACTTGTGCCTGAACACCATATCCATCCGGCG**TAATACGACTCACTATAGG**GAGAGCG
GCCGCCAGATCTTCCGGATGGCTCGAGTTTTTCAGCAAGATGGCCTTATAACCATGCAAATGTGTC
CATTAAGTTGGACTTGGAATGAGTGAATGAGTATTACTGCCAGTGTGTGTGTGTGTGTGTGTGTGT
GTGTGTGTGTGTGTGTGTCTGTCTGTCTGTCTGTCTGTCTGTCTGTCTG**CCTGCCTGCCTGCCTGC**
**CTGCCTGCCTGCCTGCCTGCCTGCCTGCCTGCCTGCCTGCCTGCCTGCCTGCCTGCCTGCCTGCCT**
**GCCTGCCTGCCTGCCTGCCTGCCTGCCTGCCTGCCTGCCTGCCTGCCTGCCTGCCTGCCTGCCTGC**
**CTGCCTGCCTGCCTGCCTGCCTGCCTGCCTGCCTGCCTGCCTGCCTGCCTGCCTGCCTGCCTGCCT**
**GCCTGCCTGCCTG**TCTGTCTGTCTCACTTTGTCCCCTAGGCTGGAGTGCAGTGGTATGATCTCGGC
TCAATCTTTCTAGAAGATCTCCTACAATATTCTCAGCTGCCATGGAAAATCGATGTTCTTCTTTTA
TTCTCTCAAGATTTTCAGGCTGTATATTAAAACTTATATTAAGAACTATGCTAACCACCTCATCAG
GAACCGTTGTAGGTGGCGTGGGTTTTCTTGGCAATCGACTCTCATGAAAACTACGAGCTAAATATT
CAATATGTTCCTCTTGACCAACTTTATTCTGCATTTTTTTTGAACGAGGTTTAGAGCAAGCTTCAG
GAAACTGAGACAGGAATTTTATTAAAAATTTAAATTTTGAAGAAAGTTCAGGGTTAATAGCATCCA
TTTTTTGCTTTGCAAGTTCCTCAGCATTCTTAACAAAAGACGTCTCTTTTGACATGTTTAAAGTTT
AAACCTCCTGTGTGAAATTATTATCCGCTCATAATTCCACACATTATACGAGCCGGAAGCATAAAG
TGTAAAGCCTGGGGTGCCTAATGAGTGAGCTAACTCACATTAATTGCGTTGCGCTCACTGCCAATT
GCTTTCCAGTCGGGAAACCTGTCGTGCCAGCTGCATTAATGAATCGGCCAACGCGCGGGGAGAGGC
GGTTTGCGTATTGGGCGCTCTTCCGCTTCCTCGCTCACTGACTCGCTGCGCTCGGTCGTTCGGCTG
CGGCGAGCGGTATCAGCTCACTCAAAGGCGGTAATACGGTTATCCACAGAATCAGGGGATAACGCA
GGAAAGAACATGTGAGCAAAAGGCCAGCAAAAGGCCAGGAACCGTAAAAAGGCCGCGTTGCTGGCG
TTTTTCCATAGGCTCCGCCCCCCTGACGAGCATCACAAAAATCGACGCTCAAGTCAGAGGTGGCGA
AACCCGACAGGACTATAAAGATACCAGGCGTTTCCCCCTGGAAGCTCCCTCGTGCGCTCTCCTGTT
CCGACCCTGCCGCTTACCGGATACCTGTCCGCCTTTCTCCCTTCGGGAAGCGTGGCGCTTTCTCAT
AGCTCACGCTGTAGGTATCTCAGTTCGGTGTAGGTCGTTCGCTCCAAGCTGGGCTGTGTGCACGAA
CCCCCCGTTCAGCCCGACCGCTGCGCCTTATCCGGTAACTATCGTCTTGAGTCCAACCCGGTAAGA
CACGACTTATCGCCACTGGCAGCAGCCACTGGTAACAGGATTAGCAGAGCGAGGTATGTAGGCGGT
GCTACAGAGTTCTTGAAGTGGTGGCCTAACTACGGCTACACTAGAAGGACAGTATTTGGTATCTGC
GCTCTGCTGAAGCCAGTTACCTTCGGAAAAAGAGTTGGTAGCTCTTGATCCGGCAAACAAACCACC
GCTGGTAGCGGTGGTTTTTTTGTTTGCAAGCAGCAGATTACGCGCAGAAAAAAAGGATCTCAAGAA
GATCCTTTGATCTTTTCTACGGGGTCTGACGCTCAGTGGAACGAAAACTCACGTTAAGGGATTTTG
GTCATGAGATTATCAAAAAGGATCTTCACCTAGATCCTTTTAAATTAAAAATGAAGTTTTAAATCA
ATCTAAAGTATATATGAGTAAACTTGGTCTGACAGTTACCAATGCTTAATCAGTGAGGCACCTATC
TCAGCGATCTGTCTATTTCGTTCATCCATAGTTGCCTGACTCCCCGTCGTGTAGATAACTACGATA

```
CGGGAGGGCTTACCATCTGGCCCCAGTGCTGCAATGATACCGCGAGACCCACGCTCACCGGCTCCA
GATTTATCAGCAATAAACCAGCCAGCCGGAAGGGCCGAGCGCAGAAGTGGTCCTGCAACTTTATCC
GCCTCCATCCAGTCTATTAATTGTTGCCGGGAAGCTAGAGTAAGTAGTTCGCCAGTTAATAGTTTG
CGCAACGTTGTTGCCATTGCTACAGGCATCGTGGTGTCACGCTCGTCGTTTGGTATGGCTTCATTC
AGCTCCGGTTCCCAACGATCAAGGCGAGTTACATGATCCCCCATGTTGTGCAAAAAAGCGGTTAGC
TCCTTCGGTCCTCCGATCGTTGTCAGAAGTAAGTTGGCCGCAGTGTTATCACTCATGGTTATGGCA
GCACTGCATAATTCTCTTACTGTCATGCCATCCGTAAGATGCTTTTCTGTGACTGGTGAGTACTCA
ACCAAGTCATTCTGAGAATAGTGTATGCGGCGACCGAGTTGCTCTTGCCCGGCGTCAATACGGGAT
AATACCGCGCCACATAGCAGAACTTTAAAAGTGCTCATCATTGGAAAACGTTCTTCGGGGCGAAAA
CTCTCAAGGATCTTACCGCTGTTGAGATCCAGTTCGATGTAACCCACTCGTGCACCCAACTGATCT
TCAGCATCTTTTACTTTCACCAGCGTTTCTGGGTGAGCAAAAACAGGAAGGCAAAATGCCGCAAAA
AAGGGAATAAGGGCGACACGGAAATGTTGAATACTCATACTCTTCCTTTTTCAATATTATTGAAGC
ATTTATCAGGGTTATTGTCTCATGAGCGGATACATATTTGAATGTATTTAGAAAAATAAACAAATA
GGGGTTCCGCGCACATTTCCCCGAAAAGTGCCACCTGACGTCTAAGAAACCATTATTATCATGACA
TTAACCTATAAAAATAGGCGTATCACGAGGCC
```

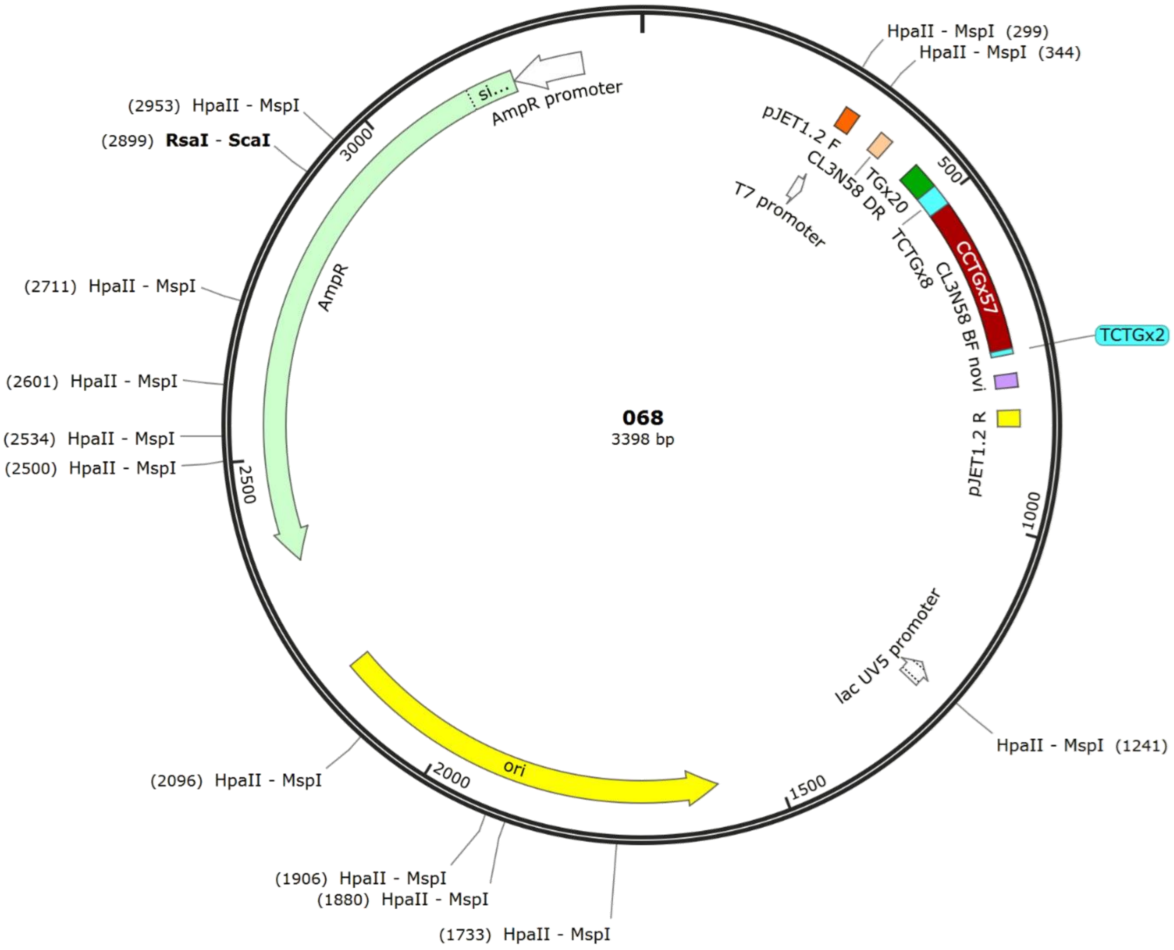

#### Table S3.

Sequence of DNA oligonucleotides used for assembly of '1111' RNA:DNA nanostructure used for the study of RNA containing 51 CUG repeats (Figure 1d).

▲ Complementary DNA oligonucleotide ▲ Overhang for '1' generation

| Oligo number | Sequence (5′→ 3′) |
|---|---|
| 1 | AGTTGGGTGCACGAGTGGGTTACATCG |
| 2 | AACTGGATCTCAACAGCGGTAAGATTTTGGATATCACTCATTAGTGGT |
| 3 | CCTTGAGAGTTTTCGCCCCGAAGAACGTTTTCCAATGATG |
| 4 | AGCACTTTTAAAGTTCTGCTATGTGGCGCGGTATTATCCC |
| 5 | GTATTGACGCCGGGCAAGAGCAACTCGGTCGCCGCATACA |
| 6 | CTATTCTCAGAATGACTTGGTTGAGTACTCACCAGTCACA |
| 7 | GAAAAGCATCTTACGGATGGCATGACAGTAAGAGAATTAT |
| 8 | GCAGTGCTGCCATAACCATGAGTGATAACACTGCGGCCAA |
| 9 | CTTACTTCTGACAACGATCGGAGGACCGAAGGAGCTAACC |
| 10 | GCTTTTTTGCACAACATGGGGGATCATGTAACTCGCCTTG |
| 11 | ATCGTTGGGAACCGGAGCTGAATGAAGCCATACCAAACGA |
| 12 | CGAGCGTGACACCACGATGCCTGTAGCAATGGCAACAACG |
| 13 | TTGCGCAAACTATTAACTGGCGAACTACTTACTCTAGCTT |
| 14 | CCCGGCAACAATTAATAGACTGGATGGAGGCGGATAAAGT |
| 15 | TGCAGGACCACTTCTGCGCTCGGCCCTTCCGGCTGGCTGG |
| 16 | TTTATTGCTGATAAATCTGGAGCCGGT |
| 17 | GAGCGTGGGTCTCGCGGTATCATTGCAG |
| **18** | CACTGGGGCCAGATGGTAAGCCCTCTTTGGATATCACTCATTAGTGGT |
| 19 | CCGTATCGTAGTTATCTACACGACGGGGAGTCAGGCAACT |
| 20 | ATGGATGAACGAAATAGACAGATCGCTGAGATAGGTGCCT |
| 21 | CACTGATTAAGCATTGGTAACTGTCAGACCAAGTTTACTC |
| 22 | ATATATACTTTAGATTGATTTAAAACTTCATTTTTAATTT |
| 23 | AAAAGGATCTAGGTGAAGATCCTTTTTGATAATCTCATGA |
| 24 | CCAAAATCCCTTAACGTGAGTTTTCGTTCCACTGAGCGTC |
| 25 | AGACCCCGTAGAAAAGATCAAAGGATCTTCTTGAGATCCT |
| 26 | TTTTTTCTGCGCGTAATCTGCTGCTTGCAAACAAAAAAAC |
| 27 | CACCGCTACCAGCGGTGGTTTGTTTGCCGGATCAAGAGCT |
| 28 | ACCAACTCTTTTTCCGAAGGTAACTGGCTTCAGCAGAGCG |
| 29 | CAGATACCAAATACTGTCCTTCTAGTGTAGCCGTAGTTAG |
| 30 | GCCACCACTTCAAGAACTCTGTAGCACCGCCTACATACCT |
| 31 | CGCTCTGCTAATCCTGTTACCAGTGGCTGCTGCCAGTGGC |
| 32 | GATAAGTCGTGTCTTACCGGGTTGGAC |
| 33 | TCAAGACGATAGTTACCGGATAAGGCGC |
| **34** | AGCGGTCGGGCTGAACGGGGGGTTCTTTGGATATCACTCATTAGTGGT |
| 35 | GTGCACACAGCCCAGCTTGGAGCGAACGACCTACACCGAA |
| 36 | CTGAGATACCTACAGCGTGAGCTATGAGAAAGCGCCACGC |

| | |
|---|---|
| 37 | TTCCCGAAGGGAGAAAGGCGGACAGGTATCCGGTAAGCGG |
| 38 | CAGGGTCGGAACAGGAGAGCGCACGAGGGAGCTTCCAGGG |
| 39 | GGAAACGCCTGGTATCTTTATAGTCCTGTCGGGTTTCGCC |
| 40 | ACCTCTGACTTGAGCGTCGATTTTTGTGATGCTCGTCAGG |
| 41 | GGGGCGGAGCCTATGGAAAAACGCCAGCAACGCGGCCTTT |
| 42 | TTACGGTTCCTGGCCTTTTGCTGGCCTTTTGCTCACATGT |
| 43 | TCTTTCCTGCGTTATCCCCTGATTCTGTGGATAACCGTAT |
| 44 | TACCGCCTTTGAGTGAGCTGATACCGCTCGCCGCAGCCGA |
| 45 | ACGACCGAGCGCAGCGAGTCAGTGAGCGAGGAAGCGGAAG |
| 46 | AGCGCCCAATACGCAAACCGCCTCTCCCCGCGCGTTGGCC |
| 47 | GATTCATTAATGCAGCTGGCACGACAGGTTTCCCGACTGG |
| 48 | AAAGCAATTGGCAGTGAGCGCAACGCA |
| 49 | ATTAATGTGAGTTAGCTCACTCATTAGG |
| 50 | CACCCCAGGCTTTACACTTTATGCTTTTGGATATCACTCATTAGTGGT |
| 51 | TCCGGCTCGTATAATGTGTGGAATTATGAGCGGATAATAA |
| 52 | TTTCACACAGGAGGTTTAAACTTTAAACATGTCAAAAGAG |
| 53 | ACGTCTTTTGTTAAGAATGCTGAGGAACTTGCAAAGCAAA |
| 54 | AAATGGATGCTATTAACCCTGAACTTTCTTCAAAATTTAA |
| 55 | ATTTTTAATAAAATTCCTGTCTCAGTTTCCTGAAGCTTGC |
| 56 | TCTAAACCTCGTTCAAAAAAAATGCAGAATAAAGTTGGTC |
| 57 | AAGAGGAACATATTGAATATTTAGCTCGTAGTTTTCATGA |
| 58 | GAGTCGATTGCCAAGAAAACCCACGCCACCTACAACGGTT |
| 59 | CCTGATGAGGTGGTTAGCATAGTTCTTAATATAAGTTTTA |
| 60 | ATATACAGCCTGAAAATCTTGAGAGAATAAAAGAAGAACA |
| 61 | TCGATTTTCCATGGCAGCTGAGAATATTGTAGGAGATCTT |
| 62 | CTAGAAAGATTTAAGCCGAGAATGGTCTGTGATCCCCC |
| 63 | CATTCCCGGCTACACTGCACCATGATCTTGCTGAAAAACT |
| 64 | CGAGCCATCCGGAAGATCTGGCGGCCGCTCTCCC |
| 65 | AGTTGGGTGCACGAGTGGGTTACATCG |
| 66 | AACTGGATCTCAACAGCGGTAAGATTTTGGATATCACTCATTAGTGGT |
| 67 | CCTTGAGAGTTTTCGCCCCGAAGAACGTTTTCCAATGATG |
| 68 | AGCACTTTTAAAGTTCTGCTATGTGGCGCGGTATTATCCC |
| 69 | GTATTGACGCCGGGCAAGAGCAACTCGGTCGCCGCATACA |
| 70 | CTATTCTCAGAATGACTTGGTTGAGTACTCACCAGTCACA |
| 71 | GAAAAGCATCTTACGGATGGCATGACAGTAAGAGAATTAT |
| 72 | GCAGTGCTGCCATAACCATGAGTGATAACACTGCGGCCAA |
| 73 | CTTACTTCTGACAACGATCGGAGGACCGAAGGAGCTAACC |
| 74 | GCTTTTTTGCACAACATGGGGGATCATGTAACTCGCCTTG |
| 75 | ATCGTTGGGAACCGGAGCTGAATGAAGCCATACCAAACGA |
| 76 | CGAGCGTGACACCACGATGCCTGTAGCAATGGCAACAACG |

#### Table S4.

Sequence of DNA oligonucleotides used for assembly of '1001' RNA:DNA nanostructure used for the study of RNA containing 57 CCUG repeats (Figure 1e).

▲ Complementary DNA oligonucleotide ▲ Overhang for '1' generation

| Oligo number | Sequence (5'→ 3') |
|---|---|
| 1 | AGCGTGATGCTACTAATTGGGACAATTTTCCAGATGAAGT |
| 2 | ATCATCTAAGAATTTAAATGAAGAAGACTTCAGAGCTTTT |
| 3 | GTTAAAAATTATTTGGCAAAAATAATATAATTCGGCTGCA |
| 4 | GGGGCGGCCTCGTGATACGCCTATTTTTATAGGTTAATGT |
| 5 | CATGATAATAATGGTTTCTTAGACGTCAGGTGGCACTTTT |
| 6 | CGGGGAAATGTGCGCGGAACCCCTATTTGTTTATTTTTCT |
| 7 | AAATACATTCAAATATGTATCCGCTCATGAGACAATAACC |
| 8 | CTGATAAATGCTTCAATAATATTGAAAAAGGAAGAGTATG |
| 9 | AGTATTCAACATTTCCGTGTCGCCCTTATTCCCTTTTTTG |
| 10 | CGGCATTTTGCCTTCCTGTTTTTGCTCACCCAGAAACGCT |
| 11 | GGTGAAAGTAAAAGATGCTGAAGATCAGTTGGGTGCACGA |
| 12 | GTGGGTTACATCGAACTGGATCTCAAC |
| 13 | AGCGGTAAGATCCTTGAGAGTTTTCTTTGGATATCACTCATTAGTGGT |
| 14 | GCCCCGAAGAACGTTTTCCAATGATGAGCACTTTTAAAGT |
| 15 | TCTGCTATGTGGCGCGGTATTATCCCGTATTGACGCCGGG |
| 16 | CAAGAGCAACTCGGTCGCCGCATACACTATTCTCAGAATG |
| 17 | ACTTGGTTGAGTACTCACCAGTCACAGAAAAGCATCTTAC |
| 18 | GGATGGCATGACAGTAAGAGAATTATGCAGTGCTGCCATA |
| 19 | ACCATGAGTGATAACACTGCGGCCAACTTACTTCTGACAA |
| 20 | CGATCGGAGGACCGAAGGAGCTAACCGCTTTTTTGCACAA |
| 21 | CATGGGGGATCATGTAACTCGCCTTGATCGTTGGGAACCG |
| 22 | GAGCTGAATGAAGCCATACCAAACGACGAGCGTGACACCA |
| 23 | CGATGCCTGTAGCAATGGCAACAACGTTGCGCAAACTATT |
| 24 | AACTGGCGAACTACTTACTCTAGCTTCCCGGCAACAATTA |
| 25 | ATAGACTGGATGGAGGCGGATAAAGTTGCAGGACCACTTC |
| 26 | TGCGCTCGGCCCTTCCGGCTGGCTGGTTTATTGCTGATAA |
| 27 | ATCTGGAGCCGGTGAGCGTGGGTCTCG |
| 28 | CGGTATCATTGCAGCACTGGGGCCAGAT |
| 29 | GGTAAGCCCTCCCGTATCGTAGTTA |
| 30 | TCTACACGACGGGGAGTCAGGCAACTATGGATGAACGAAA |
| 31 | TAGACAGATCGCTGAGATAGGTGCCTCACTGATTAAGCAT |
| 32 | TGGTAACTGTCAGACCAAGTTTACTCATATATACTTTAGA |
| 33 | TTGATTTAAAACTTCATTTTTAATTTAAAAGGATCTAGGT |
| 34 | GAAGATCCTTTTTGATAATCTCATGACCAAAATCCCTTAA |
| 35 | CGTGAGTTTTCGTTCCACTGAGCGTCAGACCCCGTAGAAA |
| 36 | AGATCAAAGGATCTTCTTGAGATCCTTTTTTTCTGCGCGT |

| | |
|---|---|
| **37** | AATCTGCTGCTTGCAAACAAAAAAACCACCCTACCAGCG |
| **38** | GTGGTTTGTTTGCCGGATCAAGAGCTACCAACTCTTTTTC |
| **39** | CGAAGGTAACTGGCTTCAGCAGAGCGCAGATACCAAATAC |
| **40** | TGTCCTTCTAGTGTAGCCGTAGTTAGGCCACCACTTCAAG |
| **41** | AACTCTGTAGCACCGCCTACATACCTCGCTCTGCTAATCC |
| **42** | TGTTACCAGTGGCTGCTGCCAGTGGCGATAAGTCGTGTCT |
| **43** | TACCGGGTTGGACTCAAGACGATAGTT |
| **44** | ACCGGATAAGGCGCAGCGGTCGGGCTGA |
| **45** | ACGGGGGGTTCGTGCACACAGCCCA |
| **46** | GCTTGGAGCGAACGACCTACACCGAACTGAGATACCTACA |
| **47** | GCGTGAGCTATGAGAAAGCGCCACGCTTCCCGAAGGGAGA |
| **48** | AAGGCGGACAGGTATCCGGTAAGCGGCAGGGTCGGAACAG |
| **49** | GAGAGCGCACGAGGGAGCTTCCAGGGGGAAACGCCTGGTA |
| **50** | TCTTTATAGTCCTGTCGGGTTTCGCCACCTCTGACTTGAG |
| **51** | CGTCGATTTTTGTGATGCTCGTCAGGGGGGCGGAGCCTAT |
| **52** | GGAAAAACGCCAGCAACGCGGCCTTTTTACGGTTCCTGGC |
| **53** | CTTTTGCTGGCCTTTTGCTCACATGTTCTTTCCTGCGTTA |
| **54** | TCCCCTGATTCTGTGGATAACCGTATTACCGCCTTTGAGT |
| **55** | GAGCTGATACCGCTCGCCGCAGCCGAACGACCGAGCGCAG |
| **56** | CGAGTCAGTGAGCGAGGAAGCGGAAGAGCGCCCAATACGC |
| **57** | AAACCGCCTCTCCCCGCGCGTTGGCCGATTCATTAATGCA |
| **58** | GCTGGCACGACAGGTTTCCCGACTGGAAAGCAATTGGCAG |
| **59** | TGAGCGCAACGCAATTAATGTGAGTTA |
| **60** | GCTCACTCATTAGGCACCCCAGGCTTTA |
| **61** | CACTTTATGCTTCCGGCTCGTATAATTTGGATATCACTCATTAGTGGT |
| **62** | TGTGTGGAATTATGAGCGGATAATAATTTCACACAGGAGG |
| **63** | TTTAAACTTTAAACATGTCAAAAGAGACGTCTTTTGTTAA |
| **64** | GAATGCTGAGGAACTTGCAAAGCAAAAAATGGATGCTATT |
| **65** | AACCCTGAACTTTCTTCAAAATTTAAATTTTTAATAAAAT |
| **66** | TCCTGTCTCAGTTTCCTGAAGCTTGCTCTAAACCTCGTTC |
| **67** | AAAAAAAATGCAGAATAAAGTTGGTCAAGAGGAACATATT |
| **68** | GAATATTTAGCTCGTAGTTTTCATGAGAGTCGATTGCCAA |
| **69** | GAAAACCCACGCCACCTACAACGGTTCCTGATGAGGTGGT |
| **70** | TAGCATAGTTCTTAATATAAGTTTTAATATACAGCCTGAA |
| **71** | AATCTTGAGAGAATAAAAGAAGAACATCGATTTTCCATGG |
| **72** | CAGCTGAGAATATTGTAGGAGATCTTCTAGAAAGATTGAG |
| **73** | CCGAGATCATACCACTGCACTCCAGC |
| **74** | CTAGGGGACAAAGTGAGACAGACAGA |
| **75** | CAGACAGACAGACAGACAGACAGACAGACAGACACACACA |
| **76** | CACACACACACACACACACACACACACACACACTGGCAGT |

#### Table S5.

Sequence of DNA oligonucleotides used for assembly of ‘0011’ RNA:DNA nanostructure used for the quantitative characterization of RNA containing 12 CUG repeats (Figure 2).

▲ Complementary DNA oligonucleotide ▲ DNA Dumbbells

| Oligo number | Sequence (5′→ 3′) |
|---|---|
| 1 | AGCGTGATGCTACTAATTGGGACAATTTTCCAGATGAAGT |
| 2 | ATCATCTAAGAATTTAAATGAAGAAGACTTCAGAGCTTTT |
| 3 | GTTAAAAATTATTTGGCAAAAATAATATAATTCGGCTGCA |
| 4 | GGGGCGGCCTCGTGATACGCCTATTTTTATAGGTTAATGT |
| 5 | CATGATAATAATGGTTTCTTAGACGTCAGGTGGCACTTTT |
| 6 | CGGGGAAATGTGCGCGGAACCCCTATTTGTTTATTTTTCT |
| 7 | AAATACATTCAAATATGTATCCGCTCATGAGACAATAACC |
| 8 | CTGATAAATGCTTCAATAATATTGAAAAAGGAAGAGTATG |
| 9 | AGTATTCAACATTTCCGTGTCGCCCTTATTCCCTTTTTTG |
| 10 | CGGCATTTTGCCTTCCTGTTTTTGCTCACCCAGAAACGCT |
| 11 | GGTGAAAGTAAAAGATGCTGAAGATCAGTTGGGTGCACGA |
| 12 | GTGGGTTACATCGAACTGGATCTCAACAGCGGTAAGATCC |
| 13 | TTGAGAGTTTTCGCCCCGAAGAACGTTTTCCAATGATGAG |
| 14 | CACTTTTAAAGTTCTGCTATGTGGCGCGGTATTATCCCGT |
| 15 | ATTGACGCCGGGCAAGAGCAACTCGGTCGCCGCATACACT |
| 16 | ATTCTCAGAATGACTTGGTTGAGTACTCACCAGTCACAGA |
| 17 | AAAGCATCTTACGGATGGCATGACAGTAAGAGAATTATGC |
| 18 | AGTGCTGCCATAACCATGAGTGATAACACTGCGGCCAACT |
| 19 | TACTTCTGACAACGATCGGAGGACCGAAGGAGCTAACCGC |
| 20 | TTTTTTGCACAACATGGGGGATCATGTAACTCGCCTTGAT |
| 21 | CGTTGGGAACCGGAGCTGAATGAAGCCATACCAAACGACG |
| 22 | AGCGTGACACCACGATGCCTGTAGCAATGGCAACAACGTT |
| 23 | GCGCAAACTATTAACTGGCGAACTACTTACTCTAGCTTCC |
| 24 | CGGCAACAATTAATAGACTGGATGGAGGCGGATAAAGTTG |
| 25 | CAGGACCACTTCTGCGCTCGGCCCTTCCGGCTGGCTGGTT |
| 26 | TATTGCTGATAAATCTGGAGCCGGTGAGCGTGGGTCTCGC |
| 27 | GGTATCATTGCAGCACTGGGGCCAGATGGTAAGCCCTCCC |
| 28 | GTATCGTAGTTATCTACACGACGGGGAGTCAGGCAACTAT |
| 29 | GGATGAACGAAATAGACAGATCGCTGAGATAGGTGCCTCA |
| 30 | CTGATTAAGCATTGGTAACTGTCAGACCAAGTTTACTCAT |
| 31 | ATATACTTTAGATTGATTTAAAACTTCATTTTTAATTTAA |
| 32 | AAGGATCTAGGTGAAGATCCTTTTTGATAATCTCATGACC |
| 33 | AAAATCCCTTAACGTGAGTTTTCGTTCCACTGAGCGTCAG |
| 34 | ACCCCGTAGAAAAGATCAAAGGATCTTCTTGAGATCCTTT |
| 35 | TTTTCTGCGCGTAATCTGCTGCTTGCAAACAAAAAAACCA |
| 36 | CCGCTACCAGTCCTCTTTTGAGGAACAAGTTTTCTTGTCGGTGGTTTG |
| 37 | TTTGCCGGATTCCTCTTTTGAGGAACAAGTTTTCTTGTCAAGAGCTAC |

| | |
|---|---|
| **38** | CAACTCTTTTTCCTCTTTTGAGGAACAAGTTTTCTTGTTCCGAAGGTA |
| **39** | ACTGGCTTCATCCTCTTTTGAGGAACAAGTTTTCTTGTGCAGAGCGCA |
| **40** | GATACCAAATTCCTCTTTTGAGGAACAAGTTTTCTTGTACTGTCCTTC |
| **41** | TAGTGTAGCCTCCTCTTTTGAGGAACAAGTTTTCTTGTGTAGTTAGGC |
| **42** | CACCACTTCATCCTCTTTTGAGGAACAAGTTTTCTTGTAGAACTCTGT |
| **43** | AGCACCGCCTTCCTCTTTTGAGGAACAAGTTTTCTTGTACATACCTCG |
| **44** | CTCTGCTAATTCCTCTTTTGAGGAACAAGTTTTCTTGTCCTGTTACCA |
| **45** | GTGGCTGCTGTCCTCTTTTGAGGAACAAGTTTTCTTGTCCAGTGGCGA |
| **46** | TAAGTCGTGTTCCTCTTTTGAGGAACAAGTTTTCTTGTCTTACCGGGT |
| **47** | TGGACTCAAGTCCTCTTTTGAGGAACAAGTTTTCTTGTACGATAGTTA |
| **48** | CCGGATAAGGCGCAGCGGTCGGGCTGAACGGGGGGTTCGT |
| **49** | GCACACAGCCCAGCTTGGAGCGAACGACCTACACCGAACT |
| **50** | GAGATACCTACAGCGTGAGCTATGAGAAAGCGCCACGCTT |
| **51** | CCCGAAGGGAGAAAGGCGGACAGGTATCCGGTAAGCGGCA |
| **52** | GGGTCGGAACAGGAGAGCGCACGAGGGAGCTTCCAGGGGG |
| **53** | AAACGCCTGGTATCTTTATAGTCCTGTCGGGTTTCGCCAC |
| **54** | CTCTGACTTGAGCGTCGATTTTTGTGATGCTCGTCAGGGG |
| **55** | GGCGGAGCCTATGGAAAAACGCCAGCAACGCGGCCTTTTT |
| **56** | ACGGTTCCTGGCCTTTTGCTGGCCTTTTGCTCACATGTTC |
| **57** | TTTCCTGCGTTCCTCTTTTGAGGAACAAGTTTTCTTGTTATCCCCTGA |
| **58** | TTCTGTGGATTCCTCTTTTGAGGAACAAGTTTTCTTGTAACCGTATTA |
| **59** | CCGCCTTTGATCCTCTTTTGAGGAACAAGTTTTCTTGTGTGAGCTGAT |
| **60** | ACCGCTCGCCTCCTCTTTTGAGGAACAAGTTTTCTTGTGCAGCCGAAC |
| **61** | GACCGAGCGCTCCTCTTTTGAGGAACAAGTTTTCTTGTAGCGAGTCAG |
| **62** | TGAGCGAGGATCCTCTTTTGAGGAACAAGTTTTCTTGTAGCGGAAGAG |
| **63** | CGCCCAATACTCCTCTTTTGAGGAACAAGTTTTCTTGTGCAAACCGCC |
| **64** | TCTCCCCGCGTCCTCTTTTGAGGAACAAGTTTTCTTGTCGTTGGCCGA |
| **65** | TTCATTAATGTCCTCTTTTGAGGAACAAGTTTTCTTGTCAGCTGGCAC |
| **66** | GACAGGTTTCTCCTCTTTTGAGGAACAAGTTTTCTTGTCCGACTGGAA |
| **67** | AGCAATTGGCTCCTCTTTTGAGGAACAAGTTTTCTTGTAGTGAGCGCA |
| **68** | ACGCAATTAATCCTCTTTTGAGGAACAAGTTTTCTTGTTGTGAGTTAG |
| **69** | CTCACTCATTAGGCACCCCAGGCTTTACACTTTATGCTTC |
| **70** | CGGCTCGTATAATGTGTGGAATTATGAGCGGATAATAATT |
| **71** | TCACACAGGAGGTTTAAACTTTAAACATGTCAAAAGAGAC |
| **72** | GTCTTTTGTTAAGAATGCTGAGGAACTTGCAAAGCAAAAA |
| **73** | ATGGATGCTATTAACCCTGAACTTTCTTCAAAATTTAAAT |
| **74** | TTTTAATAAAATTCCTGTCTCAGTTTCCTGAAGCTTGCTC |
| **75** | TAAACCTCGTTCAAAAAAAATGCAGAATAAAGTTGGTCAA |
| **76** | GAGGAACATATTGAATATTTAGCTCGTAGTTTTCATGAGA |
| **77** | GTCGATTGCCAAGAAAACCCACGCCACCTACAACGGTTCC |
| **78** | TGATGAGGTGGTTAGCATAGTTCTTAATATAAGTTTTAAT |
| **79** | ATACAGCCTGAAAATCTTGAGAGAATAAAAGAAGAACATC |
| **80** | GATTTTCCATGGCAGCTGAGAATATTGTAGGAGATCTTCT |
| **81** | AGAAAGATTTAAGCCGAGAATGGTCTGTGATCCCCC |
| **82** | CATTCCCGGCTACACTGCACCATGATCTTGCTGAAAAACT |
| **83** | CGAGCCATCCGGAAGATCTGGCGGCCGCTCTCCC |

#### Table S6.

Sequence of DNA oligonucleotide used for assembly of '1001' RNA:DNA nanostructure used for the quantitative characterization of RNA containing 24 CUG repeats (Figure 2).

▲ Complementary DNA oligonucleotide ▲ DNA Dumbbells

| Oligo number | Sequence (5′→ 3′) |
|---|---|
| 1 | AGCGTGATGCTACTAATTGGGACAATTTTCCAGATGAAGT |
| 2 | ATCATCTAAGAATTTAAATGAAGAAGACTTCAGAGCTTTT |
| 3 | GTTAAAAATTATTTGGCAAAAATAATATAATTCGGCTGCA |
| 4 | GGGGCGGCCTCGTGATACGCCTATTTTTATAGGTTAATGT |
| 5 | CATGATAATAATGGTTTCTTAGACGTCAGGTGGCACTTTT |
| **6** | CGGGGAAATGTCCTCTTTTGAGGAACAAGTTTTCTTGTTGCGCGGAAC |
| **7** | CCCTATTTGTTCCTCTTTTGAGGAACAAGTTTTCTTGTTTATTTTTCT |
| **8** | AAATACATTCTCCTCTTTTGAGGAACAAGTTTTCTTGTAAATATGTAT |
| **9** | CCGCTCATGATCCTCTTTTGAGGAACAAGTTTTCTTGTGACAATAACC |
| **10** | CTGATAAATGTCCTCTTTTGAGGAACAAGTTTTCTTGTCTTCAATAAT |
| **11** | ATTGAAAAAGTCCTCTTTTGAGGAACAAGTTTTCTTGTGAAGAGTATG |
| **12** | AGTATTCAACTCCTCTTTTGAGGAACAAGTTTTCTTGTATTTCCGTGT |
| **13** | CGCCCTTATTTCCTCTTTTGAGGAACAAGTTTTCTTGTCCCTTTTTTG |
| **14** | CGGCATTTTGTCCTCTTTTGAGGAACAAGTTTTCTTGTCCTTCCTGTT |
| **15** | TTTGCTCACCTCCTCTTTTGAGGAACAAGTTTTCTTGTCAGAAACGCT |
| **16** | GGTGAAAGTATCCTCTTTTGAGGAACAAGTTTTCTTGTAAAGATGCTG |
| **17** | AAGATCAGTTTCCTCTTTTGAGGAACAAGTTTTCTTGTGGGTGCACGA |
| 18 | GTGGGTTACATCGAACTGGATCTCAACAGCGGTAAGATCC |
| 19 | TTGAGAGTTTTCGCCCCGAAGAACGTTTTCCAATGATGAG |
| 20 | CACTTTTAAAGTTCTGCTATGTGGCGCGGTATTATCCCGT |
| 21 | ATTGACGCCGGGCAAGAGCAACTCGGTCGCCGCATACACT |
| 22 | ATTCTCAGAATGACTTGGTTGAGTACTCACCAGTCACAGA |
| 23 | AAAGCATCTTACGGATGGCATGACAGTAAGAGAATTATGC |
| 24 | AGTGCTGCCATAACCATGAGTGATAACACTGCGGCCAACT |
| 25 | TACTTCTGACAACGATCGGAGGACCGAAGGAGCTAACCGC |
| 26 | TTTTTTGCACAACATGGGGGATCATGTAACTCGCCTTGAT |
| 27 | CGTTGGGAACCGGAGCTGAATGAAGCCATACCAAACGACG |
| 28 | AGCGTGACACCACGATGCCTGTAGCAATGGCAACAACGTT |
| 29 | GCGCAAACTATTAACTGGCGAACTACTTACTCTAGCTTCC |
| 30 | CGGCAACAATTAATAGACTGGATGGAGGCGGATAAAGTTG |
| 31 | CAGGACCACTTCTGCGCTCGGCCCTTCCGGCTGGCTGGTT |
| 32 | TATTGCTGATAAATCTGGAGCCGGTGAGCGTGGGTCTCGC |
| 33 | GGTATCATTGCAGCACTGGGGCCAGATGGTAAGCCCTCCC |
| 34 | GTATCGTAGTTATCTACACGACGGGGAGTCAGGCAACTAT |
| 35 | GGATGAACGAAATAGACAGATCGCTGAGATAGGTGCCTCA |
| 36 | CTGATTAAGCATTGGTAACTGTCAGACCAAGTTTACTCAT |
| 37 | ATATACTTTAGATTGATTTAAAACTTCATTTTTAATTTAA |

| | |
|---|---|
| 38 | AAGGATCTAGGTGAAGATCCTTTTTGATAATCTCATGACC |
| 39 | AAAATCCCTTAACGTGAGTTTTCGTTCCACTGAGCGTCAG |
| 40 | ACCCCGTAGAAAAGATCAAAGGATCTTCTTGAGATCCTTT |
| 41 | TTTTCTGCGCGTAATCTGCTGCTTGCAAACAAAAAAACCA |
| 42 | CCGCTACCAGCGGTGGTTTGTTTGCCGGATCAAGAGCTAC |
| 43 | CAACTCTTTTTCCGAAGGTAACTGGCTTCAGCAGAGCGCA |
| 44 | GATACCAAATACTGTCCTTCTAGTGTAGCCGTAGTTAGGC |
| 45 | CACCACTTCAAGAACTCTGTAGCACCGCCTACATACCTCG |
| 46 | CTCTGCTAATCCTGTTACCAGTGGCTGCTGCCAGTGGCGA |
| 47 | TAAGTCGTGTCTTACCGGGTTGGACTCAAGACGATAGTTA |
| 48 | CCGGATAAGGCGCAGCGGTCGGGCTGAACGGGGGGTTCGT |
| 49 | GCACACAGCCCAGCTTGGAGCGAACGACCTACACCGAACT |
| 50 | GAGATACCTACAGCGTGAGCTATGAGAAAGCGCCACGCTT |
| 51 | CCCGAAGGGAGAAAGGCGGACAGGTATCCGGTAAGCGGCA |
| 52 | GGGTCGGAACAGGAGAGCGCACGAGGGAGCTTCCAGGGGG |
| 53 | AAACGCCTGGTATCTTTATAGTCCTGTCGGGTTTCGCCAC |
| 54 | CTCTGACTTGAGCGTCGATTTTTGTGATGCTCGTCAGGGG |
| 55 | GGCGGAGCCTATGGAAAAACGCCAGCAACGCGGCCTTTTT |
| 56 | ACGGTTCCTGGCCTTTTGCTGGCCTTTTGCTCACATGTTC |
| 57 | TTTCCTGCGTTCCTCTTTTGAGGAACAAGTTTTCTTGTTATCCCCTGA |
| 58 | TTCTGTGGATTCCTCTTTTGAGGAACAAGTTTTCTTGTAACCGTATTA |
| 59 | CCGCCTTTGATCCTCTTTTGAGGAACAAGTTTTCTTGTGTGAGCTGAT |
| 60 | ACCGCTCGCCTCCTCTTTTGAGGAACAAGTTTTCTTGTGCAGCCGAAC |
| 61 | GACCGAGCGCTCCTCTTTTGAGGAACAAGTTTTCTTGTAGCGAGTCAG |
| 62 | TGAGCGAGGATCCTCTTTTGAGGAACAAGTTTTCTTGTAGCGGAAGAG |
| 63 | CGCCCAATACTCCTCTTTTGAGGAACAAGTTTTCTTGTGCAAACCGCC |
| 64 | TCTCCCCGCGTCCTCTTTTGAGGAACAAGTTTTCTTGTCGTTGGCCGA |
| 65 | TTCATTAATGTCCTCTTTTGAGGAACAAGTTTTCTTGTCAGCTGGCAC |
| 66 | GACAGGTTTCTCCTCTTTTGAGGAACAAGTTTTCTTGTCCGACTGGAA |
| 67 | AGCAATTGGCTCCTCTTTTGAGGAACAAGTTTTCTTGTAGTGAGCGCA |
| 68 | ACGCAATTAATCCTCTTTTGAGGAACAAGTTTTCTTGTTGTGAGTTAG |
| 69 | CTCACTCATTAGGCACCCCAGGCTTTACACTTTATGCTTC |
| 70 | CGGCTCGTATAATGTGTGGAATTATGAGCGGATAATAATT |
| 71 | TCACACAGGAGGTTTAAACTTTAAACATGTCAAAAGAGAC |
| 72 | GTCTTTTGTTAAGAATGCTGAGGAACTTGCAAAGCAAAAA |
| 73 | ATGGATGCTATTAACCCTGAACTTTCTTCAAAATTTAAAT |
| 74 | TTTTAATAAAATTCCTGTCTCAGTTTCCTGAAGCTTGCTC |
| 75 | TAAACCTCGTTCAAAAAAAATGCAGAATAAAGTTGGTCAA |
| 76 | GAGGAACATATTGAATATTTAGCTCGTAGTTTTCATGAGA |
| 77 | GTCGATTGCCAAGAAAACCCACGCCACCTACAACGGTTCC |
| 78 | TGATGAGGTGGTTAGCATAGTTCTTAATATAAGTTTTAAT |
| 79 | ATACAGCCTGAAAATCTTGAGAGAATAAAAGAAGAACATC |
| 80 | GATTTTCCATGGCAGCTGAGAATATTGTAGGAGATCTTCT |
| 81 | AGAAAGATTTAAGCCGAGAATGGTCTGTGATCCCCC |
| 82 | CATTCCCGGCTACACTGCACCATGATCTTGCTGAAAACT |
| 83 | CGAGCCATCCGGAAGATCTGGCGGCCGCTCTCCC |

#### Table S7.

Sequence of DNA oligonucleotides used for assembly of '1110' RNA:DNA nanostructure used for the quantitative characterization of RNA containing 30 CUG repeats (Figure 2).

▲ Complementary DNA oligonucleotides ▲ DNA Dumbbells

| Oligo number | Sequence (5′→ 3′) |
|---|---|
| 1 | AGCGTGATGCTACTAATTGGGACAATTTTCCAGATGAAGT |
| 2 | ATCATCTAAGAATTTAAATGAAGAAGACTTCAGAGCTTTT |
| 3 | GTTAAAAATTATTTGGCAAAAATAATATAATTCGGCTGCA |
| 4 | GGGGCGGCCTCGTGATACGCCTATTTTTATAGGTTAATGT |
| 5 | CATGATAATAATGGTTTCTTAGACGTCAGGTGGCACTTTT |
| 6 | CGGGGAAATGTCCTCTTTTGAGGAACAAGTTTTCTTGTTGCGCGGAAC |
| 7 | CCCTATTTGTTCCTCTTTTGAGGAACAAGTTTTCTTGTTTATTTTTCT |
| 8 | AAATACATTCTCCTCTTTTGAGGAACAAGTTTTCTTGTAAATATGTAT |
| 9 | CCGCTCATGATCCTCTTTTGAGGAACAAGTTTTCTTGTGACAATAACC |
| 10 | CTGATAAATGTCCTCTTTTGAGGAACAAGTTTTCTTGTCTTCAATAAT |
| 11 | ATTGAAAAAGTCCTCTTTTGAGGAACAAGTTTTCTTGTGAAGAGTATG |
| 12 | AGTATTCAACTCCTCTTTTGAGGAACAAGTTTTCTTGTATTTCCGTGT |
| 13 | CGCCCTTATTTCCTCTTTTGAGGAACAAGTTTTCTTGTCCCTTTTTTG |
| 14 | CGGCATTTTGTCCTCTTTTGAGGAACAAGTTTTCTTGTCCTTCCTGTT |
| 15 | TTTGCTCACCTCCTCTTTTGAGGAACAAGTTTTCTTGTCAGAAACGCT |
| 16 | GGTGAAAGTATCCTCTTTTGAGGAACAAGTTTTCTTGTAAAGATGCTG |
| 17 | AAGATCAGTTTCCTCTTTTGAGGAACAAGTTTTCTTGTGGGTGCACGA |
| 18 | GTGGGTTACATCGAACTGGATCTCAACAGCGGTAAGATCC |
| 19 | TTGAGAGTTTTCGCCCCGAAGAACGTTTTCCAATGATGAG |
| 20 | CACTTTTAAAGTTCTGCTATGTGGCGCGGTATTATCCCGT |
| 21 | ATTGACGCCGGGCAAGAGCAACTCGGTCGCCGCATACACT |
| 22 | ATTCTCAGAATGACTTGGTTGAGTACTCACCAGTCACAGA |
| 23 | AAAGCATCTTACGGATGGCATGACAGTAAGAGAATTATGC |
| 24 | AGTGCTGCCATAACCATGAGTGATAACACTGCGGCCAACT |
| 25 | TACTTCTGACAACGATCGGAGGACCGAAGGAGCTAACCGC |
| 26 | TTTTTTGCACAACATGGGGGATCATGTAACTCGCCTTGAT |
| 27 | CGTTGGGAACTCCTCTTTTGAGGAACAAGTTTTCTTGTCGGAGCTGAA |
| 28 | TGAAGCCATATCCTCTTTTGAGGAACAAGTTTTCTTGTCCAAACGACG |
| 29 | AGCGTGACACTCCTCTTTTGAGGAACAAGTTTTCTTGTCACGATGCCT |
| 30 | GTAGCAATGGTCCTCTTTTGAGGAACAAGTTTTCTTGTCAACAACGTT |
| 31 | GCGCAAACTATCCTCTTTTGAGGAACAAGTTTTCTTGTTTAACTGGCG |
| 32 | AACTACTTACTCCTCTTTTGAGGAACAAGTTTTCTTGTTCTAGCTTCC |
| 33 | CGGCAACAATTCCTCTTTTGAGGAACAAGTTTTCTTGTTAATAGACTG |
| 34 | GATGGAGGCGTCCTCTTTTGAGGAACAAGTTTTCTTGTGATAAAGTTG |
| 35 | CAGGACCACTTCCTCTTTTGAGGAACAAGTTTTCTTGTTCTGCGCTCG |
| 36 | GCCCTTCCGGTCCTCTTTTGAGGAACAAGTTTTCTTGTCTGGCTGGTT |
| 37 | TATTGCTGATTCCTCTTTTGAGGAACAAGTTTTCTTGTAAATCTGGAG |

| | |
|---|---|
| **38** | CCGGTGAGCGTCCTCTTTTGAGGAACAAGTTTTCTTGTTGGGTCTCGC |
| 39 | GGTATCATTGCAGCACTGGGGCCAGATGGTAAGCCCTCCC |
| 40 | GTATCGTAGTTATCTACACGACGGGGAGTCAGGCAACTAT |
| 41 | GGATGAACGAAATAGACAGATCGCTGAGATAGGTGCCTCA |
| 42 | CTGATTAAGCATTGGTAACTGTCAGACCAAGTTTACTCAT |
| 43 | ATATACTTTAGATTGATTTAAAACTTCATTTTTAATTTAA |
| 44 | AAGGATCTAGGTGAAGATCCTTTTTGATAATCTCATGACC |
| 45 | AAAATCCCTTAACGTGAGTTTTCGTTCCACTGAGCGTCAG |
| 46 | ACCCCGTAGAAAAGATCAAAGGATCTTCTTGAGATCCTTT |
| 47 | TTTTCTGCGCGTAATCTGCTGCTTGCAAACAAAAAAACCA |
| **48** | CCGCTACCAGTCCTCTTTTGAGGAACAAGTTTTCTTGTCGGTGGTTTG |
| **49** | TTTGCCGGATTCCTCTTTTGAGGAACAAGTTTTCTTGTCAAGAGCTAC |
| **50** | CAACTCTTTTTCCTCTTTTGAGGAACAAGTTTTCTTGTTCCGAAGGTA |
| **51** | ACTGGCTTCATCCTCTTTTGAGGAACAAGTTTTCTTGTGCAGAGCGCA |
| **52** | GATACCAAATTCCTCTTTTGAGGAACAAGTTTTCTTGTACTGTCCTTC |
| **53** | TAGTGTAGCCTCCTCTTTTGAGGAACAAGTTTTCTTGTGTAGTTAGGC |
| **54** | CACCACTTCATCCTCTTTTGAGGAACAAGTTTTCTTGTAGAACTCTGT |
| **55** | AGCACCGCCTTCCTCTTTTGAGGAACAAGTTTTCTTGTACATACCTCG |
| **56** | CTCTGCTAATTCCTCTTTTGAGGAACAAGTTTTCTTGTCCTGTTACCA |
| **57** | GTGGCTGCTGTCCTCTTTTGAGGAACAAGTTTTCTTGTCCAGTGGCGA |
| **58** | TAAGTCGTGTTCCTCTTTTGAGGAACAAGTTTTCTTGTCTTACCGGGT |
| **59** | TGGACTCAAGTCCTCTTTTGAGGAACAAGTTTTCTTGTACGATAGTTA |
| 60 | CCGGATAAGGCGCAGCGGTCGGGCTGAACGGGGGGTTCGT |
| 61 | GCACACAGCCCAGCTTGGAGCGAACGACCTACACCGAACT |
| 62 | GAGATACCTACAGCGTGAGCTATGAGAAAGCGCCACGCTT |
| 63 | CCCGAAGGGAGAAAGGCGGACAGGTATCCGGTAAGCGGCA |
| 64 | GGGTCGGAACAGGAGAGCGCACGAGGGAGCTTCCAGGGGG |
| 65 | AAACGCCTGGTATCTTTATAGTCCTGTCGGGTTTCGCCAC |
| 66 | CTCTGACTTGAGCGTCGATTTTTGTGATGCTCGTCAGGGG |
| 67 | GGCGGAGCCTATGGAAAAACGCCAGCAACGCGGCCTTTTT |
| 68 | ACGGTTCCTGGCCTTTTGCTGGCCTTTTGCTCACATGTTC |
| 69 | TTTCCTGCGTTATCCCCTGATTCTGTGGATAACCGTATTA |
| 70 | CCGCCTTTGAGTGAGCTGATACCGCTCGCCGCAGCCGAAC |
| 71 | GACCGAGCGCAGCGAGTCAGTGAGCGAGGAAGCGGAAGAG |
| 72 | CGCCCAATACGCAAACCGCCTCTCCCCGCGCGTTGGCCGA |
| 73 | TTCATTAATGCAGCTGGCACGACAGGTTTCCCGACTGGAA |
| 74 | AGCAATTGGCAGTGAGCGCAACGCAATTAATGTGAGTTAG |
| 75 | CTCACTCATTAGGCACCCCAGGCTTTACACTTTATGCTTC |
| 76 | CGGCTCGTATAATGTGTGGAATTATGAGCGGATAATAATT |
| 77 | TCACACAGGAGGTTTAAACTTTAAACATGTCAAAAGAGAC |
| 78 | GTCTTTTGTTAAGAATGCTGAGGAACTTGCAAAGCAAAAA |
| 79 | ATGGATGCTATTAACCCTGAACTTTCTTCAAAATTTAAAT |
| 80 | TTTTAATAAAATTCCTGTCTCAGTTTCCTGAAGCTTGCTC |
| 81 | TAAACCTCGTTCAAAAAAAATGCAGAATAAAGTTGGTCAA |
| 82 | GAGGAACATATTGAATATTTAGCTCGTAGTTTTCATGAGA |
| 83 | GTCGATTGCCAAGAAAACCCACGCCACCTACAACGGTTCC |

| | |
|---|---|
| **84** | TGATGAGGTGGTTAGCATAGTTCTTAATATAAGTTTTAAT |
| **85** | ATACAGCCTGAAAATCTTGAGAGAATAAAAGAAGAACATC |
| **86** | GATTTTCCATGGCAGCTGAGAATATTGTAGGAGATCTTCT |
| **87** | AGAAAGATTTAAGCCGAGAATGGTCTGTGATCCCCC |
| **88** | CATTCCCGGCTACACTGCACCATGATCTTGCTGAAAAACT |
| **89** | CGAGCCATCCGGAAGATCTGGCGGCCGCTCTCCC |

#### Table S8.

Sequence of DNA oligonucleotides used for assembly of '1111' RNA:DNA nanostructure used for the quantitative characterization of RNA containing 51 CUG repeats (Figure 2).

▲ Complementary DNA oligonucleotide ▲ DNA Dumbbells

| Oligo number | Sequence (5'→ 3') |
|---|---|
| 1 | AGCGTGATGCTACTAATTGGGACAATTTTCCAGATGAAGT |
| 2 | ATCATCTAAGAATTTAAATGAAGAAGACTTCAGAGCTTTT |
| 3 | GTTAAAAATTATTTGGCAAAAATAATATAATTCGGCTGCA |
| 4 | GGGGCGGCCTCGTGATACGCCTATTTTTATAGGTTAATGT |
| 5 | CATGATAATAATGGTTTCTTAGACGTCAGGTGGCACTTTT |
| 6 | CGGGGAAATGTCCTCTTTTGAGGAACAAGTTTTCTTGTTGCGCGGAAC |
| 7 | CCCTATTTGTTCCTCTTTTGAGGAACAAGTTTTCTTGTTTATTTTTCT |
| 8 | AAATACATTCTCCTCTTTTGAGGAACAAGTTTTCTTGTAAATATGTAT |
| 9 | CCGCTCATGATCCTCTTTTGAGGAACAAGTTTTCTTGTGACAATAACC |
| 10 | CTGATAAATGTCCTCTTTTGAGGAACAAGTTTTCTTGTCTTCAATAAT |
| 11 | ATTGAAAAAGTCCTCTTTTGAGGAACAAGTTTTCTTGTGAAGAGTATG |
| 12 | AGTATTCAACTCCTCTTTTGAGGAACAAGTTTTCTTGTATTTCCGTGT |
| 13 | CGCCCTTATTTCCTCTTTTGAGGAACAAGTTTTCTTGTCCCTTTTTTG |
| 14 | CGGCATTTTGTCCTCTTTTGAGGAACAAGTTTTCTTGTCCTTCCTGTT |
| 15 | TTTGCTCACCTCCTCTTTTGAGGAACAAGTTTTCTTGTCAGAAACGCT |
| 16 | GGTGAAAGTATCCTCTTTTGAGGAACAAGTTTTCTTGTAAAGATGCTG |
| 17 | AAGATCAGTTTCCTCTTTTGAGGAACAAGTTTTCTTGTGGGTGCACGA |
| 18 | GTGGGTTACATCGAACTGGATCTCAACAGCGGTAAGATCC |
| 19 | TTGAGAGTTTTCGCCCCGAAGAACGTTTTCCAATGATGAG |
| 20 | CACTTTTAAAGTTCTGCTATGTGGCGCGGTATTATCCCGT |
| 21 | ATTGACGCCGGGCAAGAGCAACTCGGTCGCCGCATACACT |
| 22 | ATTCTCAGAATGACTTGGTTGAGTACTCACCAGTCACAGA |
| 23 | AAAGCATCTTACGGATGGCATGACAGTAAGAGAATTATGC |
| 24 | AGTGCTGCCATAACCATGAGTGATAACACTGCGGCCAACT |
| 25 | TACTTCTGACAACGATCGGAGGACCGAAGGAGCTAACCGC |
| 26 | TTTTTTGCACAACATGGGGGATCATGTAACTCGCCTTGAT |
| 27 | CGTTGGGAACTCCTCTTTTGAGGAACAAGTTTTCTTGTCGGAGCTGAA |
| 28 | TGAAGCCATATCCTCTTTTGAGGAACAAGTTTTCTTGTCCAAACGACG |
| 29 | AGCGTGACACTCCTCTTTTGAGGAACAAGTTTTCTTGTCACGATGCCT |
| 30 | GTAGCAATGGTCCTCTTTTGAGGAACAAGTTTTCTTGTCAACAACGTT |
| 31 | GCGCAAACTATCCTCTTTTGAGGAACAAGTTTTCTTGTTTAACTGGCG |
| 32 | AACTACTTACTCCTCTTTTGAGGAACAAGTTTTCTTGTTCTAGCTTCC |
| 33 | CGGCAACAATTCCTCTTTTGAGGAACAAGTTTTCTTGTTAATAGACTG |
| 34 | GATGGAGGCGTCCTCTTTTGAGGAACAAGTTTTCTTGTGATAAAGTTG |
| 35 | CAGGACCACTTCCTCTTTTGAGGAACAAGTTTTCTTGTTCTGCGCTCG |
| 36 | GCCCTTCCGGTCCTCTTTTGAGGAACAAGTTTTCTTGTCTGGCTGGTT |
| 37 | TATTGCTGATTCCTCTTTTGAGGAACAAGTTTTCTTGTAAATCTGGAG |

| | |
|---|---|
| **38** | CCGGTGAGCGTCCTCTTTTGAGGAACAAGTTTTCTTGTTGGGTCTCGC |
| 39 | GGTATCATTGCAGCACTGGGGCCAGATGGTAAGCCCTCCC |
| 40 | GTATCGTAGTTATCTACACGACGGGGAGTCAGGCAACTAT |
| 41 | GGATGAACGAAATAGACAGATCGCTGAGATAGGTGCCTCA |
| 42 | CTGATTAAGCATTGGTAACTGTCAGACCAAGTTTACTCAT |
| 43 | ATATACTTTAGATTGATTTAAAACTTCATTTTTAATTTAA |
| 44 | AAGGATCTAGGTGAAGATCCTTTTTGATAATCTCATGACC |
| 45 | AAAATCCCTTAACGTGAGTTTTCGTTCCACTGAGCGTCAG |
| 46 | ACCCCGTAGAAAAGATCAAAGGATCTTCTTGAGATCCTTT |
| 47 | TTTTCTGCGCGTAATCTGCTGCTTGCAAACAAAAAAACCA |
| **48** | CCGCTACCAGTCCTCTTTTGAGGAACAAGTTTTCTTGTCGGTGGTTTG |
| **49** | TTTGCCGGATTCCTCTTTTGAGGAACAAGTTTTCTTGTCAAGAGCTAC |
| **50** | CAACTCTTTTTCCTCTTTTGAGGAACAAGTTTTCTTGTTCCGAAGGTA |
| **51** | ACTGGCTTCATCCTCTTTTGAGGAACAAGTTTTCTTGTGCAGAGCGCA |
| **52** | GATACCAAATTCCTCTTTTGAGGAACAAGTTTTCTTGTACTGTCCTTC |
| **53** | TAGTGTAGCCTCCTCTTTTGAGGAACAAGTTTTCTTGTGTAGTTAGGC |
| **54** | CACCACTTCATCCTCTTTTGAGGAACAAGTTTTCTTGTAGAACTCTGT |
| **55** | AGCACCGCCTTCCTCTTTTGAGGAACAAGTTTTCTTGTACATACCTCG |
| **56** | CTCTGCTAATTCCTCTTTTGAGGAACAAGTTTTCTTGTCCTGTTACCA |
| **57** | GTGGCTGCTGTCCTCTTTTGAGGAACAAGTTTTCTTGTCCAGTGGCGA |
| **58** | TAAGTCGTGTTCCTCTTTTGAGGAACAAGTTTTCTTGTCTTACCGGGT |
| **59** | TGGACTCAAGTCCTCTTTTGAGGAACAAGTTTTCTTGTACGATAGTTA |
| 60 | CCGGATAAGGCGCAGCGGTCGGGCTGAACGGGGGGTTCGT |
| 61 | GCACACAGCCCAGCTTGGAGCGAACGACCTACACCGAACT |
| 62 | GAGATACCTACAGCGTGAGCTATGAGAAAGCGCCACGCTT |
| 63 | CCCGAAGGGAGAAAGGCGGACAGGTATCCGGTAAGCGGCA |
| 64 | GGGTCGGAACAGGAGAGCGCACGAGGGAGCTTCCAGGGGG |
| 65 | AAACGCCTGGTATCTTTATAGTCCTGTCGGGTTTCGCCAC |
| 66 | CTCTGACTTGAGCGTCGATTTTTGTGATGCTCGTCAGGGG |
| 67 | GGCGGAGCCTATGGAAAAACGCCAGCAACGCGGCCTTTTT |
| 68 | ACGGTTCCTGGCCTTTTGCTGGCCTTTTGCTCACATGTTC |
| **69** | TTTCCTGCGTTCCTCTTTTGAGGAACAAGTTTTCTTGTTATCCCCTGA |
| **70** | TTCTGTGGATTCCTCTTTTGAGGAACAAGTTTTCTTGTAACCGTATTA |
| **71** | CCGCCTTTGATCCTCTTTTGAGGAACAAGTTTTCTTGTGTGAGCTGAT |
| **72** | ACCGCTCGCCTCCTCTTTTGAGGAACAAGTTTTCTTGTGCAGCCGAAC |
| **73** | GACCGAGCGCTCCTCTTTTGAGGAACAAGTTTTCTTGTAGCGAGTCAG |
| **74** | TGAGCGAGGATCCTCTTTTGAGGAACAAGTTTTCTTGTAGCGGAAGAG |
| **75** | CGCCCAATACTCCTCTTTTGAGGAACAAGTTTTCTTGTGCAAACCGCC |
| **76** | TCTCCCCGCGTCCTCTTTTGAGGAACAAGTTTTCTTGTCGTTGGCCGA |
| **77** | TTCATTAATGTCCTCTTTTGAGGAACAAGTTTTCTTGTCAGCTGGCAC |
| **78** | GACAGGTTTCTCCTCTTTTGAGGAACAAGTTTTCTTGTCCGACTGGAA |
| **79** | AGCAATTGGCTCCTCTTTTGAGGAACAAGTTTTCTTGTAGTGAGCGCA |
| **80** | ACGCAATTAATCCTCTTTTGAGGAACAAGTTTTCTTGTTGTGAGTTAG |
| 81 | CTCACTCATTAGGCACCCCAGGCTTTACACTTTATGCTTC |
| 82 | CGGCTCGTATAATGTGTGGAATTATGAGCGGATAATAATT |
| 83 | TCACACAGGAGGTTTAAACTTTAAACATGTCAAAAGAGAC |

| | |
|---|---|
| 84 | GTCTTTTGTTAAGAATGCTGAGGAACTTGCAAAGCAAAAA |
| 85 | ATGGATGCTATTAACCCTGAACTTTCTTCAAAATTTAAAT |
| 86 | TTTTAATAAAATTCCTGTCTCAGTTTCCTGAAGCTTGCTC |
| 87 | TAAACCTCGTTCAAAAAAAATGCAGAATAAAGTTGGTCAA |
| 88 | GAGGAACATATTGAATATTTAGCTCGTAGTTTTCATGAGA |
| 89 | GTCGATTGCCAAGAAAACCCACGCCACCTACAACGGTTCC |
| 90 | TGATGAGGTGGTTAGCATAGTTCTTAATATAAGTTTTAAT |
| 91 | ATACAGCCTGAAAATCTTGAGAGAATAAAAGAAGAACATC |
| 92 | GATTTTCCATGGCAGCTGAGAATATTGTAGGAGATCTTCT |
| 93 | AGAAAGATTTAAGCCGAGAATGGTCTGTGATCCCCC |
| 94 | CATTCCCGGCTACACTGCACCATGATCTTGCTGAAAAACT |
| 95 | CGAGCCATCCGGAAGATCTGGCGGCCGCTCTCCC |

#### Table S9.

Sequence of modified pJET1.2/blunt cloning vector (CloneJET PCR Cloning Kit, Thermo Fisher, Catalog number: K1231): Circular DNA (3721 bp) with inserted $(GCN)_{20}$ repeats.

| **Sequence (5′→ 3′)** |
|---|
| **T7 promoter** **$(GCN)_{20}$ repeats** **DraIII Cutting** |
| GCCCCTGCAGCCGAATTATATTATTTTTGCCAAATAATTTTTAACAAAAGCTCTGAAGTCTTCTTC<br>ATTTAAATTCTTAGATGATACTTCATCTGGAAAATTGTCCCAATTAGTAGCAT**CACGCTGTG**AGTA<br>AGTTCTAAACCATTTTTTTATTGTTGTATTATCTCTAATCTTACTACTCGATGAGTTTTCGGTATT<br>ATCTCTATTTTTAACTTGGAGCAGGTTCCATTCATTGTTTTTTTCATCATAGTGAATAAAATCAAC<br>TGCTTTAACACTTGTGCCTGAACACCATATCCATCCGGCG**TAATACGACTCACTATAGG**GAGAGCG<br>GCCGCCAGATCTTCCGGATGGCTCGAGTTTTTCAGCAAGATCATGGGGAGGAGGGTGTTAAAACAA<br>GCCGAAGTAGAACTTGGGCCACCCTAACCGGTGCTTTTCTTTCCCATTTTCTTCTTTCTCCCCCTG<br>CTTCACCGTCTCTCCTTCCGTCTTGGGCCAGGTGTGGTTCCAGAACCGCCGCGCCAAGTTTCGCAA<br>GCAGGAGCGCGCAGCGGCAGCCGCAGCGGCCGCGGCCAAGAACGGCTCCTCGGGCAAAAAGTCTGA<br>CTCTTCCAGGGACGACGAGAGCAAAGAGGCCAAGAGCACTGACCCGGACAGCACTGGGGGCCCAGG<br>TCCCAATCCCAACCCCACCCCCAGCTGCGGGGCGAATGGAGGCGGCGGCGGCGGGCCCAGCCCGGC<br>TGGAGCTCCGGGGGCGGCGGGGCCCGGGGGCCCGGGAGGCGAACCCGGCAAGGGCGGC**GCAGCAGC**<br>**AGCGGCGGCGGCCGCGGCAGCGGCGGCGGCGGCAGCGGCAGCGGCGGCAGCT**GGAGGCCTGGCTGC<br>GGCTGGGGGCCCTGGACAAGGCTGGGCTCCCGGCCCCGGCCCCATCACCTCCATCCCGGATTCGCT<br>TGGGGGTCCCTTCGCCAGCGTCCTATCTTCGCTCCAAAGACCCAACGGTGCCAAAGCCGCCTTAGT<br>GAAGAGCAGTATGTTCTGATCTGGAATCCTGCGGCGGCGGCGGCGGCGACAGCGGGCGAGCCA<br>GGGCCCGGGCGGGCGAGTGGGCGAGCGGGTAGGCCCAAGGCTATTGTCGTCGCTGATCTTTCTAGA<br>AGATCTCCTACAATATTCTCAGCTGCCATGGAAAATCGATGTTCTTCTTTTATTCTCTCAAGATTT<br>TCAGGCTGTATATTAAAACTTATATTAAGAACTATGCTAACCACCTCATCAGGAACCGTTGTAGGT<br>GGCGTGGGTTTTCTTGGCAATCGACTCTCATGAAAACTACGAGCTAAATATTCAATATGTTCCTCT<br>TGACCAACTTTATTCTGCATTTTTTTTGAACGAGGTTTAGAGCAAGCTTCAGGAAACTGAGACAGG<br>AATTTTATTAAAAATTTAAATTTTGAAGAAAGTTCAGGGTTAATAGCATCCATTTTTTGCTTTGCA<br>AGTTCCTCAGCATTCTTAACAAAAGACGTCTCTTTTGACATGTTTAAAGTTTAAACCTCCTGTGTG<br>AAATTATTATCCGCTCATAATTCCACACATTATACGAGCCGGAAGCATAAAGTGTAAAGTGTAAAG<br>CCTGGGGTGCCTAATGAGTGAGCTAACTCACATTAATTGCGTTGCGCTCACTGCCAATTGCTTTCC<br>AGTCGGGAAACCTGTCGTGCCAGCTGCATTAATGAATCGGCCAACGCGCGGGGAGAGGCGGTTTGC<br>GTATTGGGCGCTCTTCCGCTTCCTCGCTCACTGACTCGCTGCGCTCGGTCGTTCGGCTGCGGCGAG<br>CGGTATCAGCTCACTCAAAGGCGGTAATACGGTTATCCACAGAATCAGGGGATAACGCAGGAAAGA<br>ACATGTGAGCAAAAGGCCAGCAAAAGGCCAGGAACCGTAAAAAGGCCGCGTTGCTGGCGTTTTTCC<br>ATAGGCTCCGCCCCCCTGACGAGCATCACAAAAATCGACGCTCAAGTCAGAGGTGGCGAAACCCGA<br>CAGGACTATAAAGATACCAGGCGTTTCCCCCTGGAAGCTCCCTCGTGCGCTCTCCTGTTCCGACCC<br>TGCCGCTTACCGGATACCTGTCCGCCTTTCTCCCTTCGGGAAGCGTGGCGCTTTCTCATAGCTCAC<br>GCTGTAGGTATCTCAGTTCGGTGTAGGTCGTTCGCTCCAAGCTGGGCTGTGTGCACGAACCCCCCG<br>TTCAGCCCGACCGCTGCGCCTTATCCGGTAACTATCGTCTTGAGTCCAACCCGGTAAGACACGACT<br>TATCGCCACTGGCAGCAGCCACTGGTAACAGGATTAGCAGAGCGAGGTATGTAGGCGGTGCTACAG<br>AGTTCTTGAAGTGGTGGCCTAACTACGGCTACACTAGAAGAACAGTATTTGGTATCTGCGCTCTGC<br>TGAAGCCAGTTACCTTCGGAAAAAGAGTTGGTAGCTCTTGATCCGGCAAACAAACCACCGCTGGTA |

```
GCGGTGGTTTTTTTGTTTGCAAGCAGCAGATTACGCGCAGAAAAAAAGGATCTCAAGAAGATCCTT
TGATCTTTTCTACGGGGTCTGACGCTCAGTGGAACGAAAACTCACGTTAAGGGATTTTGGTCATGA
GATTATCAAAAAGGATCTTCACCTAGATCCTTTTAAATTAAAAATGAAGTTTTAAATCAATCTAAA
GTATATATGAGTAAACTTGGTCTGACAGTTACCAATGCTTAATCAGTGAGGCACCTATCTCAGCGA
TCTGTCTATTTCGTTCATCCATAGTTGCCTGACTCCCCGTCGTGTAGATAACTACGATACGGGAGG
GCTTACCATCTGGCCCCAGTGCTGCAATGATACCGCGAGACCCACGCTCACCGGCTCCAGATTTAT
CAGCAATAAACCAGCCAGCCGGAAGGGCCGAGCGCAGAAGTGGTCCTGCAACTTTATCCGCCTCCA
TCCAGTCTATTAATTGTTGCCGGGAAGCTAGAGTAAGTAGTTCGCCAGTTAATAGTTTGCGCAACG
TTGTTGCCATTGCTACAGGCATCGTGGTGTCACGCTCGTCGTTTGGTATGGCTTCATTCAGCTCCG
GTTCCCAACGATCAAGGCGAGTTACATGATCCCCCATGTTGTGCAAAAAAGCGGTTAGCTCCTTCG
GTCCTCCGATCGTTGTCAGAAGTAAGTTGGCCGCAGTGTTATCACTCATGGTTATGGCAGCACTGC
ATAATTCTCTTACTGTCATGCCATCCGTAAGATGCTTTTCTGTGACTGGTGAGTACTCAACCAAGT
CATTCTGAGAATAGTGTATGCGGCGACCGAGTTGCTCTTGCCCGGCGTCAATACGGGATAATACCG
CGCCACATAGCAGAACTTTAAAAGTGCTCATCATTGGAAAACGTTCTTCGGGGCGAAAACTCTCAA
GGATCTTACCGCTGTTGAGATCCAGTTCGATGTAACCCACTCGTGCACCCAACTGATCTTCAGCAT
CTTTTACTTTCACCAGCGTTTCTGGGTGAGCAAAAACAGGAAGGCAAAATGCCGCAAAAAAGGGAA
TAAGGGCGACACGGAAATGTTGAATACTCATACTCTTCCTTTTTCAATATTATTGAAGCATTTATC
AGGGTTATTGTCTCATGAGCGGATACATATTTGAATGTATTTAGAAAAATAAACAAATAGGGGTTC
CGCGCACATTTCCCCGAAAAGTGCCACCTGACGTCTAAGAAACCATTATTATCATGACATTAACCT
ATAAAAATAGGCGTATCACGAGGCC
```

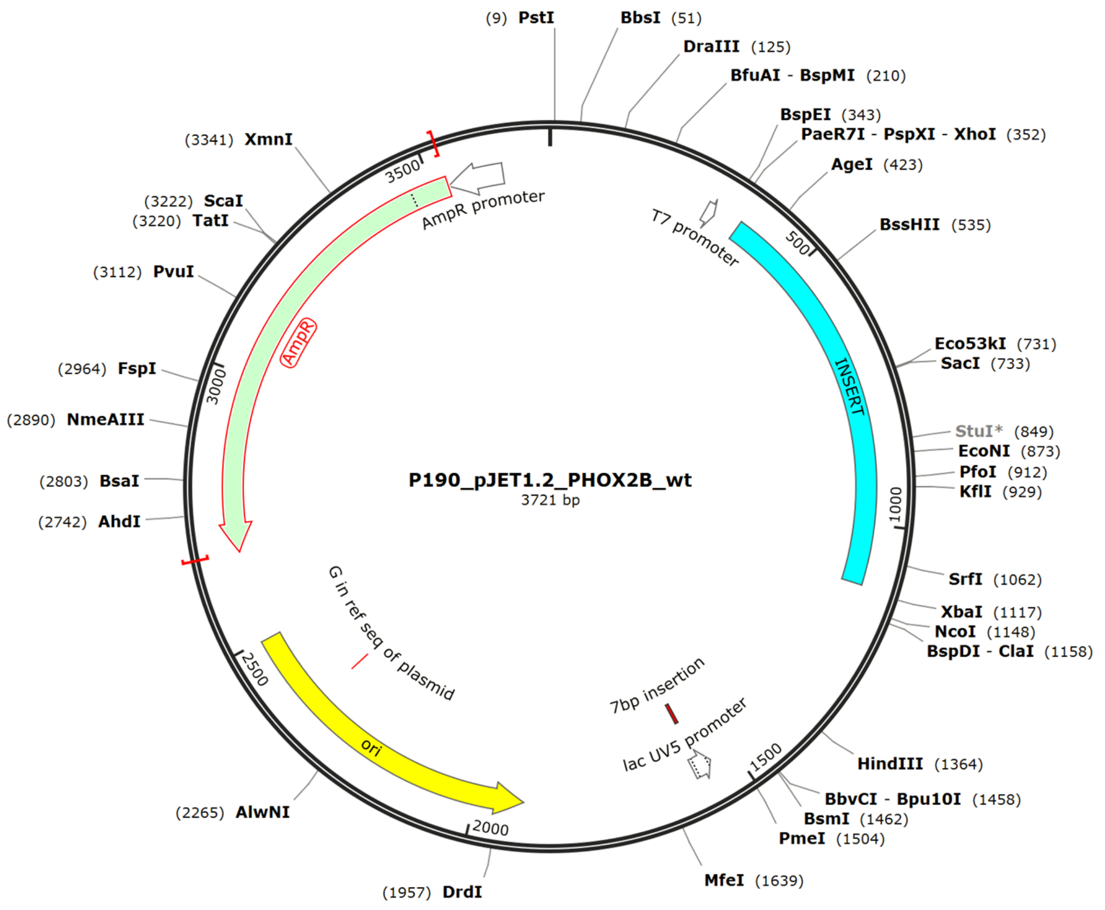

#### Table S10.

Sequence of modified pJET1.2/blunt cloning vector (CloneJET PCR Cloning Kit, Thermo Fisher, Catalog number: K1231): Circular DNA (3732 bp) with inserted $(GCN)_{26}$ repeats.

| Sequence (5′→ 3′) |
|---|
| **T7 promoter** **$(GCN)_{26}$ repeats** **DraIII Cutting** |
| GCCCCTGCAGCCGAATTATATTATTTTTGCCAAATAATTTTTAACAAAAGCTCTGAAGTCTTCTTC<br>ATTTAAATTCTTAGATGATACTTCATCTGGAAAATTGTCCCAATTAGTAGCAT**CACGCTGTG**AGTA<br>AGTTCTAAACCATTTTTTTATTGTTGTATTATCTCTAATCTTACTACTCGATGAGTTTTCGGTATT<br>ATCTCTATTTTTAACTTGGAGCAGGTTCCATTCATTGTTTTTTTCATCATAGTGAATAAAATCAAC<br>TGCTTTAACACTTGTGCCTGAACACCATATCCATCCGGCG**TAATACGACTCACTATAGG**GAGAGCG<br>GCCGCCAGATCTTCCGGATGGCTCGAGTTTTTCAGCAAGATCATGGGGAGGAGGGTGTTAAAACAA<br>GCCGAAGTAGAACTTGGGCCACCCTAACCGGTGCTTTTCTTTCCCATTTTCTTCTTTCTCCCCCTG<br>CTTCACCGTCTCTCCTTCCGTCTTGGGCCAGGTGTGGTTCCAGAACCGCCGCGCCAAGTTTCGCAA<br>GCAGGAGCGCGCAGCGGCAGCCGCAGCGGCCGCGGCCAAGAACGGCTCCTCGGGCAAAAAGTCTGA<br>CTCTTCCAGGGACGACGAGAGCAAAGAGGCCAAGAGCACTGACCCGGACAGCACTGGGGGCCCAGG<br>TCCCAATCCCAACCCCACCCCCAGCTGCGGGGCGAATGGAGGCGGCGGCGGCGGGCCCAGCCCGGC<br>TGGAGCTCCGGGGGCGGCGGGGCCCGGGGGCCCGGGAGGCGAACCCGGCAAGGGCGGC**GCAGCAGC**<br>**AGCGGCGGCGGCCGCGGCAGCGGCGGCGGCCGCGGCAGCGGCGGCGGCGGCAGCGGCAGCGGCGGC**<br>**AGCT**GGAGGCCTGGCTGCGGCTGGGGGCCCTGGACAAGGCTGGGCTCCCGGCCCCGGCCCCATCAC<br>CTCCATCCCGGATTCGCTTGGGGGTCCCTTCGCCAGCGTCCTATCTTCGCTCCAAAGACCCAACGG<br>TGCCAAAGCCGCCTTAGTGAAGAGCAGTATGTTCTGATCTGGAATCCTGCGGCGGCGGCGGCGGCG<br>GCGACAGCGGGCGAGCCAGGGCCCGGGCGGGCGAGTGGGCGAGCGGGTAGGCCCAAGGCTATTGTC<br>GTCGCTGATCTTTCTAGAAGATCTCCTACAATATTCTCAGCTGCCATGGAAAATCGATGTTCTTCT<br>TTTATTCTCTCAAGATTTTCAGGCTGTATATTAAAACTTATATTAAGAACTATGCTAACCACCTCA<br>TCAGGAACCGTTGTAGGTGGCGTGGGTTTTCTTGGCAATCGACTCTCATGAAAACTACGAGCTAAA<br>TATTCAATATGTTCCTCTTGACCAACTTTATTCTGCATTTTTTTTGAACGAGGTTTAGAGCAAGCT<br>TCAGGAAACTGAGACAGGAATTTTATTAAAAATTTAAATTTTGAAGAAAGTTCAGGGTTAATAGCA<br>TCCATTTTTTGCTTTGCAAGTTCCTCAGCATTCTTAACAAAAGACGTCTCTTTTGACATGTTTAAA<br>GTTTAAACCTCCTGTGTGAAATTATTATCCGCTCATAATTCCACACATTATACGAGCCGGAAGCAT<br>AAAGTGTAAAGCCTGGGGTGCCTAATGAGTGAGCTAACTCACATTAATTGCGTTGCGCTCACTGCC<br>AATTGCTTTCCAGTCGGGAAACCTGTCGTGCCAGCTGCATTAATGAATCGGCCAACGCGCGGGGAG<br>AGGCGGTTTGCGTATTGGGCGCTCTTCCGCTTCCTCGCTCACTGACTCGCTGCGCTCGGTCGTTCG<br>GCTGCGGCGAGCGGTATCAGCTCACTCAAAGGCGGTAATACGGTTATCCACAGAATCAGGGGATAA<br>CGCAGGAAAGAACATGTGAGCAAAAGGCCAGCAAAAGGCCAGGAACCGTAAAAAGGCCGCGTTGCT<br>GGCGTTTTTCCATAGGCTCCGCCCCCCTGACGAGCATCACAAAAATCGACGCTCAAGTCAGAGGTG<br>GCGAAACCCGACAGGACTATAAAGATACCAGGCGTTTCCCCCTGGAAGCTCCCTCGTGCGCTCTCC<br>TGTTCCGACCCTGCCGCTTACCGGATACCTGTCCGCCTTTCTCCCTTCGGGAAGCGTGGCGCTTTC<br>TCATAGCTCACGCTGTAGGTATCTCAGTTCGGTGTAGGTCGTTCGCTCCAAGCTGGGCTGTGTGCA<br>CGAACCCCCCGTTCAGCCCGACCGCTGCGCCTTATCCGGTAACTATCGTCTTGAGTCCAACCCGGT<br>AAGACACGACTTATCGCCACTGGCAGCAGCCACTGGTAACAGGATTAGCAGAGCGAGGTATGTAGG<br>CGGTGCTACAGAGTTCTTGAAGTGGTGGCCTAACTACGGCTACACTAGAAGAACAGTATTTGGTAT<br>CTGCGCTCTGCTGAAGCCAGTTACCTTCGGAAAAAGAGTTGGTAGCTCTTGATCCGGCAAACAAAC |

CACCGCTGGTAGCGGTGGTTTTTTTGTTTGCAAGCAGCAGATTACGCGCAGAAAAAAAGGATCTCA
AGAAGATCCTTTGATCTTTTCTACGGGGTCTGACGCTCAGTGGAACGAAAACTCACGTTAAGGGAT
TTTGGTCATGAGATTATCAAAAAGGATCTTCACCTAGATCCTTTTAAATTAAAAATGAAGTTTTAA
ATCAATCTAAAGTATATATGAGTAAACTTGGTCTGACAGTTACCAATGCTTAATCAGTGAGGCACC
TATCTCAGCGATCTGTCTATTTCGTTCATCCATAGTTGCCTGACTCCCCGTCGTGTAGATAACTAC
GATACGGGAGGGCTTACCATCTGGCCCCAGTGCTGCAATGATACCGCGAGACCCACGCTCACCGGC
TCCAGATTTATCAGCAATAAACCAGCCAGCCGGAAGGGCCGAGCGCAGAAGTGGTCCTGCAACTTT
ATCCGCCTCCATCCAGTCTATTAATTGTTGCCGGGAAGCTAGAGTAAGTAGTTCGCCAGTTAATAG
TTTGCGCAACGTTGTTGCCATTGCTACAGGCATCGTGGTGTCACGCTCGTCGTTTGGTATGGCTTC
ATTCAGCTCCGGTTCCCAACGATCAAGGCGAGTTACATGATCCCCCATGTTGTGCAAAAAAGCGGT
TAGCTCCTTCGGTCCTCCGATCGTTGTCAGAAGTAAGTTGGCCGCAGTGTTATCACTCATGGTTAT
GGCAGCACTGCATAATTCTCTTACTGTCATGCCATCCGTAAGATGCTTTTCTGTGACTGGTGAGTA
CTCAACCAAGTCATTCTGAGAATAGTGTATGCGGCGACCGAGTTGCTCTTGCCCGGCGTCAATACG
GGATAATACCGCGCCACATAGCAGAACTTTAAAAGTGCTCATCATTGGAAAACGTTCTTCGGGGCG
AAAACTCTCAAGGATCTTACCGCTGTTGAGATCCAGTTCGATGTAACCCACTCGTGCACCCAACTG
ATCTTCAGCATCTTTTACTTTCACCAGCGTTTCTGGGTGAGCAAAAACAGGAAGGCAAAATGCCGC
AAAAAAGGGAATAAGGGCGACACGGAAATGTTGAATACTCATACTCTTCCTTTTTCAATATTATTG
AAGCATTTATCAGGGTTATTGTCTCATGAGCGGATACATATTTGAATGTATTTAGAAAAATAAACA
AATAGGGGTTCCGCGCACATTTCCCCGAAAAGTGCCACCTGACGTCTAAGAAACCATTATTATCAT
GACATTAACCTATAAAAATAGGCGTATCACGAGGCC

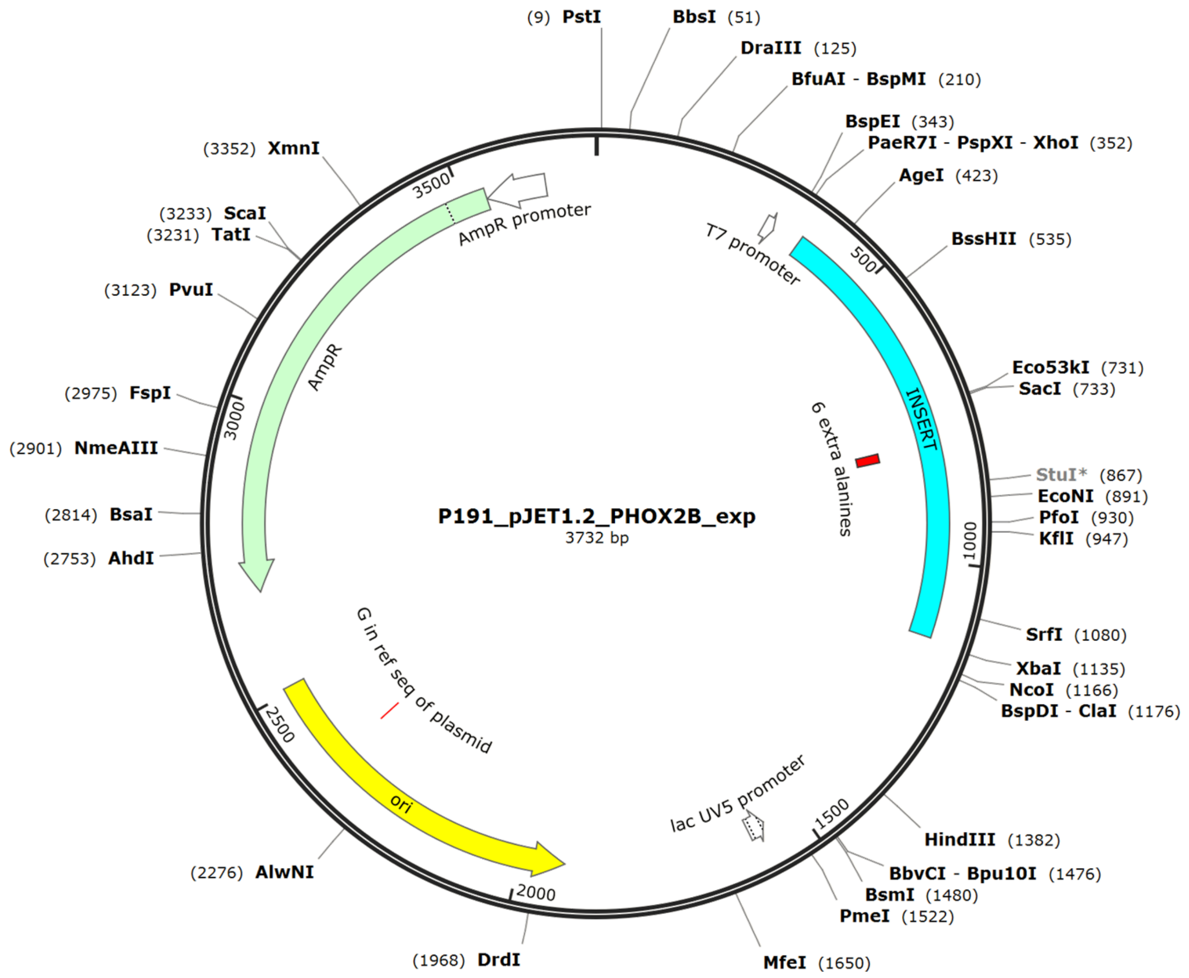

#### Table S11.

Sequence of DNA oligonucleotides used for assembly of '011' RNA:DNA nanostructure used for the quantitative characterization of RNA containing 20 GCN repeats (Figure 3).

▲ Complementary DNA oligonucleotide ▲ DNA Dumbbells

| Oligo number | Sequence (5'→ 3') |
|---|---|
| 1 | AGCGTGATGCTACTAATTGGGACAATTTTCCAGATGAAGT |
| 2 | ATCATCTAAGAATTTAAATGAAGAAGACTTCAGAGCTTTT |
| 3 | GTTAAAAATTATTTGGCAAAAATAATATAATTCGGCTGCA |
| 4 | GGGGCGGCCTCGTGATACGCCTATTTTTATAGGTTAATGT |
| 5 | CATGATAATAATGGTTTCTTAGACGTCAGGTGGCACTTTT |
| 6 | CGGGGAAATGTGCGCGGAACCCCTATTTGTTTATTTTTCT |
| 7 | AAATACATTCAAATATGTATCCGCTCATGAGACAATAACC |
| 8 | CTGATAAATGCTTCAATAATATTGAAAAAGGAAGAGTATG |
| 9 | AGTATTCAACATTTCCGTGTCGCCCTTATTCCCTTTTTTG |
| 10 | CGGCATTTTGCCTTCCTGTTTTTGCTCACCCAGAAACGCT |
| 11 | GGTGAAAGTAAAAGATGCTGAAGATCAGTTGGGTGCACGA |
| 12 | GTGGGTTACATCGAACTGGATCTCAACAGCGGTAAGATCC |
| 13 | TTGAGAGTTTTCGCCCCGAAGAACGTTTTCCAATGATGAG |
| 14 | CACTTTTAAAGTTCTGCTATGTGGCGCGGTATTATCCCGT |
| 15 | ATTGACGCCGGGCAAGAGCAACTCGGTCGCCGCATACACT |
| 16 | ATTCTCAGAATGACTTGGTTGAGTACTCACCAGTCACAGA |
| 17 | AAAGCATCTTACGGATGGCATGACAGTAAGAGAATTATGC |
| 18 | AGTGCTGCCATAACCATGAGTGATAACACTGCGGCCAACT |
| 19 | TACTTCTGACAACGATCGGAGGACCGAAGGAGCTAACCGC |
| 20 | TTTTTTGCACTCCTCTTTTGAGGAACAAGTTTTCTTGTAACATGGGGG |
| 21 | ATCATGTAACTCCTCTTTTGAGGAACAAGTTTTCTTGTTCGCCTTGAT |
| 22 | CGTTGGGAACTCCTCTTTTGAGGAACAAGTTTTCTTGTCGGAGCTGAA |
| 23 | TGAAGCCATATCCTCTTTTGAGGAACAAGTTTTCTTGTCCAAACGACG |
| 24 | AGCGTGACACTCCTCTTTTGAGGAACAAGTTTTCTTGTCACGATGCCT |
| 25 | GTAGCAATGGTCCTCTTTTGAGGAACAAGTTTTCTTGTCAACAACGTT |
| 26 | GCGCAAACTATCCTCTTTTGAGGAACAAGTTTTCTTGTTTAACTGGCG |
| 27 | AACTACTTACTCCTCTTTTGAGGAACAAGTTTTCTTGTTCTAGCTTCC |
| 28 | CGGCAACAATTCCTCTTTTGAGGAACAAGTTTTCTTGTTAATAGACTG |
| 29 | GATGGAGGCGTCCTCTTTTGAGGAACAAGTTTTCTTGTGATAAAGTTG |
| 30 | CAGGACCACTTCCTCTTTTGAGGAACAAGTTTTCTTGTTCTGCGCTCG |
| 31 | GCCCTTCCGGTCCTCTTTTGAGGAACAAGTTTTCTTGTCTGGCTGGTT |
| 32 | TATTGCTGATAAATCTGGAGCCGGTGAGCGTGGGTCTCGC |
| 33 | GGTATCATTGCAGCACTGGGGCCAGATGGTAAGCCCTCCC |
| 34 | GTATCGTAGTTATCTACACGACGGGGAGTCAGGCAACTAT |
| 35 | GGATGAACGAAATAGACAGATCGCTGAGATAGGTGCCTCA |
| 36 | CTGATTAAGCATTGGTAACTGTCAGACCAAGTTTACTCAT |
| 37 | ATATACTTTAGATTGATTTAAAACTTCATTTTTAATTTAA |

| | |
|---|---|
| 38 | AAGGATCTAGGTGAAGATCCTTTTTGATAATCTCATGACC |
| 39 | AAAATCCCTTAACGTGAGTTTTCGTTCCACTGAGCGTCAG |
| 40 | ACCCCGTAGAAAAGATCAAAGGATCTTCTTGAGATCCTTT |
| 41 | TTTTCTGCGCGTAATCTGCTGCTTGCAAACAAAAAAACCA |
| 42 | CCGCTACCAGCGGTGGTTTGTTTGCCGGATCAAGAGCTAC |
| 43 | CAACTCTTTTTCCGAAGGTAACTGGCTTCAGCAGAGCGCA |
| 44 | GATACCAAATACTGTCCTTCTAGTGTAGCCGTAGTTAGGC |
| 45 | CACCACTTCAAGAACTCTGTAGCACCGCCTACATACCTCG |
| **46** | CTCTGCTAATTCCTCTTTTGAGGAACAAGTTTTCTTGTCCTGTTACCA |
| **47** | GTGGCTGCTGTCCTCTTTTGAGGAACAAGTTTTCTTGTCCAGTGGCGA |
| **48** | TAAGTCGTGTTCCTCTTTTGAGGAACAAGTTTTCTTGTCTTACCGGGT |
| **49** | TGGACTCAAGTCCTCTTTTGAGGAACAAGTTTTCTTGTACGATAGTTA |
| **50** | CCGGATAAGGTCCTCTTTTGAGGAACAAGTTTTCTTGTCGCAGCGGTC |
| **51** | GGGCTGAACGTCCTCTTTTGAGGAACAAGTTTTCTTGTGGGGGTTCGT |
| **52** | GCACACAGCCTCCTCTTTTGAGGAACAAGTTTTCTTGTCAGCTTGGAG |
| **53** | CGAACGACCTTCCTCTTTTGAGGAACAAGTTTTCTTGTACACCGAACT |
| **54** | GAGATACCTATCCTCTTTTGAGGAACAAGTTTTCTTGTCAGCGTGAGC |
| **55** | TATGAGAAAGTCCTCTTTTGAGGAACAAGTTTTCTTGTCGCCACGCTT |
| **56** | CCCGAAGGGATCCTCTTTTGAGGAACAAGTTTTCTTGTGAAAGGCGGA |
| **57** | CAGGTATCCGTCCTCTTTTGAGGAACAAGTTTTCTTGTGTAAGCGGCA |
| 58 | GGGTCGGAACAGGAGAGCGCACGAGGGAGCTTCCAGGGGG |
| 59 | AAACGCCTGGTATCTTTATAGTCCTGTCGGGTTTCGCCAC |
| 60 | CTCTGACTTGAGCGTCGATTTTTGTGATGCTCGTCAGGGG |
| 61 | GGCGGAGCCTATGGAAAAACGCCAGCAACGCGGCCTTTTT |
| 62 | ACGGTTCCTGGCCTTTTGCTGGCCTTTTGCTCACATGTTC |
| 63 | TTTCCTGCGTTATCCCCTGATTCTGTGGATAACCGTATTA |
| 64 | CCGCCTTTGAGTGAGCTGATACCGCTCGCCGCAGCCGAAC |
| 65 | GACCGAGCGCAGCGAGTCAGTGAGCGAGGAAGCGGAAGAG |
| 66 | CGCCCAATACGCAAACCGCCTCTCCCCGCGCGTTGGCCGA |
| 67 | TTCATTAATGCAGCTGGCACGACAGGTTTCCCGACTGGAA |
| 68 | AGCAATTGGCAGTGAGCGCAACGCAATTAATGTGAGTTAG |
| 69 | CTCACTCATTAGGCACCCCAGGCTTTACACTTTATGCTTC |
| 70 | CGGCTCGTATAATGTGTGGAATTATGAGCGGATAATAATT |
| 71 | TCACACAGGAGGTTTAAACTTTAAACATGTCAAAAGAGAC |
| 72 | GTCTTTTGTTAAGAATGCTGAGGAACTTGCAAAGCAAAAA |
| 73 | ATGGATGCTATTAACCCTGAACTTTCTTCAAAATTTAAAT |
| 74 | TTTTAATAAAATTCCTGTCTCAGTTTCCTGAAGCTTGCTC |
| 75 | TAAACCTCGTTCAAAAAAAATGCAGAATAAAGTTGGTCAA |
| 76 | GAGGAACATATTGAATATTTAGCTCGTAGTTTTCATGAGA |
| 77 | GTCGATTGCCAAGAAAACCCACGCCACCTACAACGGTTCC |
| 78 | TGATGAGGTGGTTAGCATAGTTCTTAATATAAGTTTTAAT |
| 79 | ATACAGCCTGAAAATCTTGAGAGAATAAAAGAAGAACATC |
| 80 | GATTTTCCATGGCAGCTGAGAATATTGTAGGAGATCTTCT |
| 81 | AGAAAGATCAGCGACGACAATAGCCTTGGGCCTACCCGCT |
| 82 | CGCCCACTCGCCCGCCCGGGCCCTGGCTCGCCCGCTGTCG |
| 83 | CCGCCGCCGCCGCCGCCGCAGGATTCCAGATCAGAACATA |

| | |
|---|---|
| **84** | CTGCTCTTCACTAAGGCGGCTTTGGCACCGTTGGGTCTTT |
| **85** | GGAGCGAAGATAGGACGCTGGCGAAGGGACCCCCAAGCGA |
| **86** | ATCCGGGATGGAGGTGATGGGGCCGGGGCCGGGGGCCCAG |
| **87** | CCTTGTCCAGGGCCCCCAGCCGCAGCCAGGCCTCC |
| **88** | GCCGCCCTTGCCGGGTTCGCCTCCCGGGCCCCCGGGCCCC |
| **89** | GCCGCCCCCGGAGCTCCAGCCGGGCTGGGCCCGCCGCCGC |
| **90** | CGCCTCCATTCGCCCCGCAGCTGGGGGTGGGGTTGGGATT |
| **91** | GGGACCTGGGCCCCCAGTGCTGTCCGGGTCAGTGCTCTTG |
| **92** | GCCTCTTTGCTCTCGTCGTCCCTGGAAGAGTCAGACTTTT |
| **93** | TGCCCGAGGAGCCGTCCTTGGCCGCGGCCGCTGCGGCTGC |
| **94** | CGCTGCGCGCTCCTGCTTGCGAAACTTGGCGCGGCGGTTC |
| **95** | TGGAACCACACCTGGCCCAAGACGGAAGGAGAGACGGTGA |
| **96** | AGCAGGGGGAGAAAGAAGAAAATGGGAAAGAAAAGCACCG |
| **97** | GTTAGGGTGGCCCAAGTTCTACTTCGGCTTGTTTTAACAC |
| **98** | CCTCCTCCCCATGATCTTGCTGAAAAACTCGAGCCATCCG |
| **99** | GAAGATCTGGCGGCCGCTCTCCC |

▲ DNA oligonucleotides for GCN repeats

| | |
|---|---|
| **1** | CGCCGCTGCCGCGGCTTTGGATATCACTCATTAGTGGT |
| **2** | CGCTGCCGCGGCCGCTTTGGATATCACTCATTAGTGGT |
| **3** | CGCTGCCGCCGCCGCTTTGGATATCACTCATTAGTGGT |
| **4** | AGCTGCCGCCGCTGCTTTGGATATCACTCATTAGTGGT |

#### Table S12.

Sequence of DNA oligonucleotide used for assembly of '111' RNA:DNA nanostructure used for the quantitative characterization of RNA containing 26 GCN repeats (Figure 3).

▲ Complementary DNA oligonucleotide ▲ DNA Dumbbells

| Oligo number | Sequence (5'→ 3') |
|---|---|
| 1 | AGCGTGATGCTACTAATTGGGACAATTTTCCAGATGAAGT |
| 2 | ATCATCTAAGAATTTAAATGAAGAAGACTTCAGAGCTTTT |
| 3 | GTTAAAAATTATTTGGCAAAAATAATATAATTCGGCTGCA |
| 4 | GGGGCGGCCTCGTGATACGCCTATTTTTATAGGTTAATGT |
| 5 | CATGATAATAATGGTTTCTTAGACGTCAGGTGGCACTTTT |
| 6 | CGGGGAAATGTGCGCGGAACCCCTATTTGTTTATTTTTCT |
| 7 | AAATACATTCAAATATGTATCCGCTCATGAGACAATAACC |
| 8 | CTGATAAATGCTTCAATAATATTGAAAAAGGAAGAGTATG |
| 9 | AGTATTCAACATTTCCGTGTCGCCCTTATTCCCTTTTTTG |
| 10 | CGGCATTTTGCCTTCCTGTTTTTGCTCACCCAGAAACGCT |
| 11 | GGTGAAAGTAAAAGATGCTGAAGATCAGTTGGGTGCACGA |
| 12 | GTGGGTTACATCGAACTGGATCTCAACAGCGGTAAGATCC |
| 13 | TTGAGAGTTTTCGCCCCGAAGAACGTTTTCCAATGATGAG |
| 14 | CACTTTTAAAGTTCTGCTATGTGGCGCGGTATTATCCCGT |
| 15 | ATTGACGCCGGGCAAGAGCAACTCGGTCGCCGCATACACT |
| 16 | ATTCTCAGAATGACTTGGTTGAGTACTCACCAGTCACAGA |
| 17 | AAAGCATCTTACGGATGGCATGACAGTAAGAGAATTATGC |
| 18 | AGTGCTGCCATAACCATGAGTGATAACACTGCGGCCAACT |
| 19 | TACTTCTGACAACGATCGGAGGACCGAAGGAGCTAACCGC |
| 20 | TTTTTTGCACAACATGGGGGATCATGTAACTCGCCTTGAT |
| 21 | CGTTGGGAACCGGAGCTGAATGAAGCCATACCAAACGACG |
| 22 | AGCGTGACACCACGATGCCTGTAGCAATGGCAACAACGTT |
| 23 | GCGCAAACTATTAACTGGCGAACTACTTACTCTAGCTTCC |
| 24 | CGGCAACAATTAATAGACTGGATGGAGGCGGATAAAGTTG |
| 25 | CAGGACCACTTCCTCTTTTGAGGAACAAGTTTTCTTGTTCTGCGCTCG |
| 26 | GCCCTTCCGGTCCTCTTTTGAGGAACAAGTTTTCTTGTCTGGCTGGTT |
| 27 | TATTGCTGATTCCTCTTTTGAGGAACAAGTTTTCTTGTAAATCTGGAG |
| 28 | CCGGTGAGCGTCCTCTTTTGAGGAACAAGTTTTCTTGTTGGGTCTCGC |
| 29 | GGTATCATTGTCCTCTTTTGAGGAACAAGTTTTCTTGTCAGCACTGGG |
| 30 | GCCAGATGGTTCCTCTTTTGAGGAACAAGTTTTCTTGTAAGCCCTCCC |
| 31 | GTATCGTAGTTCCTCTTTTGAGGAACAAGTTTTCTTGTTATCTACACG |
| 32 | ACGGGGAGTCTCCTCTTTTGAGGAACAAGTTTTCTTGTAGGCAACTAT |
| 33 | GGATGAACGATCCTCTTTTGAGGAACAAGTTTTCTTGTAATAGACAGA |
| 34 | TCGCTGAGATTCCTCTTTTGAGGAACAAGTTTTCTTGTAGGTGCCTCA |
| 35 | CTGATTAAGCTCCTCTTTTGAGGAACAAGTTTTCTTGTATTGGTAACT |
| 36 | GTCAGACCAATCCTCTTTTGAGGAACAAGTTTTCTTGTGTTTACTCAT |
| 37 | ATATACTTTAGATTGATTTAAAACTTCATTTTTAATTTAA |

| | |
|---|---|
| 38 | AAGGATCTAGGTGAAGATCCTTTTTGATAATCTCATGACC |
| 39 | AAAATCCCTTAACGTGAGTTTTCGTTCCACTGAGCGTCAG |
| 40 | ACCCCGTAGAAAAGATCAAAGGATCTTCTTGAGATCCTTT |
| 41 | TTTTCTGCGCGTAATCTGCTGCTTGCAAACAAAAAAACCA |
| 42 | CCGCTACCAGCGGTGGTTTGTTTGCCGGATCAAGAGCTAC |
| 43 | CAACTCTTTTTCCGAAGGTAACTGGCTTCAGCAGAGCGCA |
| 44 | GATACCAAATACTGTCCTTCTAGTGTAGCCGTAGTTAGGC |
| 45 | CACCACTTCAAGAACTCTGTAGCACCGCCTACATACCTCG |
| **46** | CTCTGCTAATTCCTCTTTTGAGGAACAAGTTTTCTTGTCCTGTTACCA |
| **47** | GTGGCTGCTGTCCTCTTTTGAGGAACAAGTTTTCTTGTCCAGTGGCGA |
| **48** | TAAGTCGTGTTCCTCTTTTGAGGAACAAGTTTTCTTGTCTTACCGGGT |
| **49** | TGGACTCAAGTCCTCTTTTGAGGAACAAGTTTTCTTGTACGATAGTTA |
| **50** | CCGGATAAGGTCCTCTTTTGAGGAACAAGTTTTCTTGTCGCAGCGGTC |
| **51** | GGGCTGAACGTCCTCTTTTGAGGAACAAGTTTTCTTGTGGGGGTTCGT |
| **52** | GCACACAGCCTCCTCTTTTGAGGAACAAGTTTTCTTGTCAGCTTGGAG |
| **53** | CGAACGACCTTCCTCTTTTGAGGAACAAGTTTTCTTGTACACCGAACT |
| **54** | GAGATACCTATCCTCTTTTGAGGAACAAGTTTTCTTGTCAGCGTGAGC |
| **55** | TATGAGAAAGTCCTCTTTTGAGGAACAAGTTTTCTTGTCGCCACGCTT |
| **56** | CCCGAAGGGATCCTCTTTTGAGGAACAAGTTTTCTTGTGAAAGGCGGA |
| **57** | CAGGTATCCGTCCTCTTTTGAGGAACAAGTTTTCTTGTGTAAGCGGCA |
| 58 | GGGTCGGAACAGGAGAGCGCACGAGGGAGCTTCCAGGGGG |
| 59 | AAACGCCTGGTATCTTTATAGTCCTGTCGGGTTTCGCCAC |
| 60 | CTCTGACTTGAGCGTCGATTTTTGTGATGCTCGTCAGGGG |
| 61 | GGCGGAGCCTATGGAAAAACGCCAGCAACGCGGCCTTTTT |
| 62 | ACGGTTCCTGGCCTTTTGCTGGCCTTTTGCTCACATGTTC |
| 63 | TTTCCTGCGTTATCCCCTGATTCTGTGGATAACCGTATTA |
| 64 | CCGCCTTTGAGTGAGCTGATACCGCTCGCCGCAGCCGAAC |
| 65 | GACCGAGCGCAGCGAGTCAGTGAGCGAGGAAGCGGAAGAG |
| 66 | CGCCCAATACGCAAACCGCCTCTCCCCGCGCGTTGGCCGA |
| **67** | TTCATTAATGTCCTCTTTTGAGGAACAAGTTTTCTTGTCAGCTGGCAC |
| **68** | GACAGGTTTCTCCTCTTTTGAGGAACAAGTTTTCTTGTCCGACTGGAA |
| **69** | AGCAATTGGCTCCTCTTTTGAGGAACAAGTTTTCTTGTAGTGAGCGCA |
| **70** | ACGCAATTAATCCTCTTTTGAGGAACAAGTTTTCTTGTTGTGAGTTAG |
| **71** | CTCACTCATTTCCTCTTTTGAGGAACAAGTTTTCTTGTAGGCACCCCA |
| **72** | GGCTTTACACTCCTCTTTTGAGGAACAAGTTTTCTTGTTTTATGCTTC |
| **73** | CGGCTCGTATTCCTCTTTTGAGGAACAAGTTTTCTTGTAATGTGTGGA |
| **74** | ATTATGAGCGTCCTCTTTTGAGGAACAAGTTTTCTTGTGATAATAATT |
| **75** | TCACACAGGATCCTCTTTTGAGGAACAAGTTTTCTTGTGGTTTAAACT |
| **76** | TTAAACATGTTCCTCTTTTGAGGAACAAGTTTTCTTGTCAAAAGAGAC |
| **77** | GTCTTTTGTTTCCTCTTTTGAGGAACAAGTTTTCTTGTAAGAATGCTG |
| **78** | AGGAACTTGCTCCTCTTTTGAGGAACAAGTTTTCTTGTAAAGCAAAAA |
| 79 | ATGGATGCTATTAACCCTGAACTTTCTTCAAAATTTAAAT |
| 80 | TTTTAATAAAATTCCTGTCTCAGTTTCCTGAAGCTTGCTC |
| 81 | TAAACCTCGTTCAAAAAAAATGCAGAATAAAGTTGGTCAA |
| 82 | GAGGAACATATTGAATATTTAGCTCGTAGTTTTCATGAGA |
| 83 | GTCGATTGCCAAGAAAACCCACGCCACCTACAACGGTTCC |

| | |
|---|---|
| 84 | TGATGAGGTGGTTAGCATAGTTCTTAATATAAGTTTTAAT |
| 85 | ATACAGCCTGAAAATCTTGAGAGAATAAAAGAAGAACATC |
| 86 | GATTTTCCATGGCAGCTGAGAATATTGTAGGAGATCTTCT |
| 87 | AGAAAGATCAGCGACGACAATAGCCTTGGGCCTACCCGCT |
| 88 | CGCCCACTCGCCCGCCCGGGCCCTGGCTCGCCCGCTGTCG |
| 89 | CCGCCGCCGCCGCCGCCGCAGGATTCCAGATCAGAACATA |
| 90 | CTGCTCTTCACTAAGGCGGCTTTGGCACCGTTGGGTCTTT |
| 91 | GGAGCGAAGATAGGACGCTGGCGAAGGGACCCCCAAGCGA |
| 92 | ATCCGGGATGGAGGTGATGGGGCCGGGGCCGGGAGCCCAG |
| 93 | CCTTGTCCAGGGCCCCCAGCCGCAGCCAGGCCTCC |
| 94 | GCCGCCCTTGCCGGGTTCGCCTCCCGGGCCCCCGGGCCCC |
| 95 | GCCGCCCCCGGAGCTCCAGCCGGGCTGGGCCCGCCGCCGC |
| 96 | CGCCTCCATTCGCCCCGCAGCTGGGGGTGGGGTTGGGATT |
| 97 | GGGACCTGGGCCCCCAGTGCTGTCCGGGTCAGTGCTCTTG |
| 98 | GCCTCTTTGCTCTCGTCGTCCCTGGAAGAGTCAGACTTTT |
| 99 | TGCCCGAGGAGCCGTTCTTGGCCGCGGCCGCTGCGGCTGC |
| 100 | CGCTGCGCGCTCCTGCTTGCGAAACTTGGCGCGGCGGTTC |
| 101 | TGGAACCACACCTGGCCCAAGACGGAAGGAGAGACGGTGA |
| 102 | AGCAGGGGGAGAAAGAAGAAAATGGGAAAGAAAAGCACCG |
| 103 | GTTAGGGTGGCTCAAGTTCTACTTCGGCTTGTTTTAACAC |
| 104 | CCTCCTCCCCATGATCTTGCTGAAAAACTCGAGCCATCCG |
| 105 | GAAGATCTGGCGGCCGCTCTCCC |

▲ DNA oligonucleotides for GCN repeats

| | |
|---|---|
| 1 | CGCCGCCGCTGCTGCTGCTTTGGATATCACTCATTAGTGGT |
| 2 | CGCCGCTGCCGCGGCTTTGGATATCACTCATTAGTGGT |
| 3 | CGCTGCCGCGGCCGCTTTGGATATCACTCATTAGTGGT |
| 4 | CGCTGCCGCCGCCGCTTTGGATATCACTCATTAGTGGT |
| 5 | AGCTGCCGCCGCTGCTTTGGATATCACTCATTAGTGGT |

**Table S13.**

Sequence of modified pEGFP-N1 cloning vector: Circular DNA (4838 bp) with inserted GTG repeats.

| **Sequence (5'→ 3')** |
|---|
| **CMV promoter** **(CTG)$_{34}$** **EGFP** **polyA tail signal** |
| TAGTTATTAATAGTAATCAATTACGGGGTCATTAGTTCATAGCCCATATATGGAGTTCCGCGTTAC<br>ATAACTTACGGTAAATGGCCCGCCTGGCTGACCGCCCAACGACCCCCGCCCATTGACGTCAATAAT<br>GACGTATGTTCCCATAGTAACGCCAATAGGGACTTTCCATTGACGTCAATGGGTGGAGTATTTACG<br>GTAAACTGCCCACTTGGCAGTACATCAAGTGTATCATATGCCAAGTACGCCCCCTATTGACGTCAA<br>TGACGGTAAATGGCCCGCCTGGCATTATGCCCAGTACATGACCTTATGGGACTTTCCTACTTGGCA<br>GTACATCTACGTATTAGTCATCGCTATTACCATG**GTGATGCGGTTTTGGCAGTACATCAATGGGCG**<br>**TGGATAGCGGTTTGACTCACGGGGATTTCCAAGTCTCCACCCCATTGACGTCAATGGGAGTTTGTT**<br>**TTGGCACCAAAATCAACGGGACTTTCCAAAATGTCGTAACAACTCCGCCCCATTGACGCAAATGGG**<br>**CGGTAGGCGTGTACGGTGGGAGGTCTATATAAGCAGAGCT**GGTTTAGTGAACCGTCAGATCCGCTA<br>GCGCTACCGGACTCAGATCTCGAGCTCAAGCTTCGAATTCTGCAGTCGACGGTACGGTCTC**CTGCT**<br>**GCTGCTGCTGCTGCTGCTGCTGCTGCTGCTGCTGCTGCTGCTGCTGCTGCTGCTGCTGCTGCTGCT**<br>**GCTGCTGCTGCTGCTGCTGCTGCTGCTGCTG**CGTCTTCGTACCGCGGGCCCGGGATCCACCGGTCG<br>CCACC**ATGGTGAGCAAGGGCGAGGAGCTGTTCACCGGGGTGGTGCCCATCCTGGTCGAGCTGGACG**<br>**GCGACGTAAACGGCCACAAGTTCAGCGTGTCCGGCGAGGGCGAGGGCGATGCCACCTACGGCAAGC**<br>**TGACCCTGAAGTTCATCTGCACCACCGGCAAGCTGCCCGTGCCCTGGCCCACCCTCGTGACCACCC**<br>**TGACCTACGGCGTGCAGTGCTTCAGCCGCTACCCCGACCACATGAAGCAGCACGACTTCTTCAAGT**<br>**CCGCCATGCCCGAAGGCTACGTCCAGGAGCGCACCATCTTCTTCAAGGACGACGGCAACTACAAGA**<br>**CCCGCGCCGAGGTGAAGTTCGAGGGCGACACCCTGGTGAACCGCATCGAGCTGAAGGGCATCGACT**<br>**TCAAGGAGGACGGCAACATCCTGGGGCACAAGCTGGAGTACAACTACAACAGCCACAACGTCTATA**<br>**TCATGGCCGACAAGCAGAAGAACGGCATCAAGGTGAACTTCAAGATCCGCCACAACATCGAGGACG**<br>**GCAGCGTGCAGCTCGCCGACCACTACCAGCAGAACACCCCCATCGGCGACGGCCCCGTGCTGCTGC**<br>**CCGACAACCACTACCTGAGCACCCAGTCCGCCCTGAGCAAAGACCCCAACGAGAAGCGCGATCACA**<br>**TGGTCCTGCTGGAGTTCGTGACCGCCGCCGGGATCACTCTCGGCATGGACGAGCTGTACAAGTAA**A<br>GCGGCCGCGACTCTAGATCATAATCAGCCATACCACATTTGTAGAGGTTTTACTTGCTTTAAAAAA<br>CCTCCCACACCTCCCCCTGAACCTGAAACATAAAATGAATGCAATTGTTGTTGTT**AACTTGTTTAT**<br>**TGCAGCTTATAATGGTTACAAATAAAGCAATAGCATCACAAATTTCACAAATAAAGCATTTTTTTC**<br>**ACTGCATTCTAGTTGTGGTTTGTCCAAACTCATCAATGTATCTTA**AGGCGTAAATTGTAAGCGTTA<br>ATATTTTGTTAAAATTCGCGTTAAATTTTTGTTAAATCAGCTCATTTTTTAACCAATAGGCCGAAA<br>TCGGCAAAATCCCTTATAAATCAAAAGAATAGACCGAGATAGGGTTGAGTGTTGTTCCAGTTTGGA<br>ACAAGAGTCCACTATTAAAGAACGTGGACTCCAACGTCAAAGGGCGAAAAACCGTCTATCAGGGCG<br>ATGGCCCACTACGTGAACCATCACCCTAATCAAGTTTTTTGGGGTCGAGGTGCCGTAAAGCACTAA<br>ATCGGAACCCTAAAGGGAGCCCCCGATTTAGAGCTTGACGGGGAAAGCCGGCGAACGTGGCGAGAA<br>AGGAAGGGAAGAAAGCGAAAGGAGCGGGCGCTAGGGCGCTGGCAAGTGTAGCGGTCACGCTGCGCG<br>TAACCACCACACCCGCCGCGCTTAATGCGCCGCTACAGGGCGCGTCAGGTGGCACTTTTCGGGGAA<br>ATGTGCGCGGAACCCCTATTTGTTTATTTTTCTAAATACATTCAAATATGTATCCGCTCATGAGAC<br>AATAACCCTGATAAATGCTTCAATAATATTGAAAAAGGAAGAGTCCTGAGGCGGAAAGAACCAGCT<br>GTGGAATGTGTGTCAGTTAGGGTGTGGAAAGTCCCCAGGCTCCCCAGCAGGCAGAAGTATGCAAAG<br>CATGCATCTCAATTAGTCAGCAACCAGGTGTGGAAAGTCCCCAGGCTCCCCAGCAGGCAGAAGTAT<br>GCAAAGCATGCATCTCAATTAGTCAGCAACCATAGTCCCGCCCCTAACTCCGCCCATCCCGCCCCT<br>AACTCCGCCCAGTTCCGCCCATTCTCCGCCCCATGGCTGACTAATTTTTTTTATTTATGCAGAGGC<br>CGAGGCCGCCTCGGCCTCTGAGCTATTCCAGAAGTAGTGAGGAGGCTTTTTTGGAGGCCTAGGCTT<br>TTGCAAAGATCGATCAAGAGACAGGATGAGGATCGTTTCGCATGATTGAACAAGATGGATTGCACG<br>CAGGTTCTCCGGCCGCTTGGGTGGAGAGGCTATTCGGCTATGACTGGGCACAACAGACAATCGGCT |

```
GCTCTGATGCCGCCGTGTTCCGGCTGTCAGCGCAGGGGCGCCCGGTTCTTTTTGTCAAGACCGACC
TGTCCGGTGCCCTGAATGAACTGCAAGACGAGGCAGCGCGGCTATCGTGGCTGGCCACGACGGGCG
TTCCTTGCGCAGCTGTGCTCGACGTTGTCACTGAAGCGGGAAGGGACTGGCTGCTATTGGGCGAAG
TGCCGGGGCAGGATCTCCTGTCATCTCACCTTGCTCCTGCCGAGAAAGTATCCATCATGGCTGATG
CAATGCGGCGGCTGCATACGCTTGATCCGGCTACCTGCCCATTCGACCACCAAGCGAAACATCGCA
TCGAGCGAGCACGTACTCGGATGGAAGCCGGTCTTGTCGATCAGGATGATCTGGACGAAGAGCATC
AGGGGCTCGCGCCAGCCGAACTGTTCGCCAGGCTCAAGGCGAGCATGCCCGACGGCGAGGATCTCG
TCGTGACCCATGGCGATGCCTGCTTGCCGAATATCATGGTGGAAAATGGCCGCTTTTCTGGATTCA
TCGACTGTGGCCGGCTGGGTGTGGCGGACCGCTATCAGGACATAGCGTTGGCTACCCGTGATATTG
CTGAAGAGCTTGGCGGCGAATGGGCTGACCGCTTCCTCGTGCTTTACGGTATCGCCGCTCCCGATT
CGCAGCGCATCGCCTTCTATCGCCTTCTTGACGAGTTCTTCTGAGCGGGACTCTGGGGTTCGAAAT
GACCGACCAAGCGACGCCCAACCTGCCATCACGAGATTTCGATTCCACCGCCGCCTTCTATGAAAG
GTTGGGCTTCGGAATCGTTTTCCGGGACGCCGGCTGGATGATCCTCCAGCGCGGGGATCTCATGCT
GGAGTTCTTCGCCCACCCTAGGGGGAGGCTAACTGAAACACGGAAGGAGACAATACCGGAAGGAAC
CCGCGCTATGACGGCAATAAAAAGACAGAATAAAACGCACGGTGTTGGGTCGTTTGTTCATAAACG
CGGGGTTCGGTCCCAGGGCTGGCACTCTGTCGATCCATTGGGGCCAATACGCCCGCGTTTCTTCCT
TTTCCCCACCCCACCCCCCAAGTTCGGGTGAAGGCCCAGGGCTCGCAGCCAACGTCGGGGCGGCAG
GCCCTGCCATAGCCTCAGGTTACTCATATATACTTTAGATTGATTTAAAACTTCATTTTTAATTTA
AAAGGATCTAGGTGAAGATCCTTTTTGATAATCTCATGACCAAAATCCCTTAACGTGAGTTTTCGT
TCCACTGAGCGTCAGACCCCGTAGAAAAGATCAAAGGATCTTCTTGAGATCCTTTTTTTCTGCGCG
TAATCTGCTGCTTGCAAACAAAAAAACCACCGCTACCAGCGGTGGTTTGTTTGCCGGATCAAGAGC
TACCAACTCTTTTTCCGAAGGTAACTGGCTTCAGCAGAGCGCAGATACCAAATACTGTCCTTCTAG
TGTAGCCGTAGTTAGGCCACCACTTCAAGAACTCTGTAGCACCGCCTACATACCTCGCTCTGCTAA
TCCTGTTACCAGTGGCTGCTGCCAGTGGCGATAAGTCGTGTCTTACCGGGTTGGACTCAAGACGAT
AGTTACCGGATAAGGCGCAGCGGTCGGGCTGAACGGGGGGTTCGTGCACACAGCCCAGCTTGGAGC
GAACGACCTACACCGAACTGAGATACCTACAGCGTGAGCTATGAGAAAGCGCCACGCTTCCCGAAG
GGAGAAAGGCGGACAGGTATCCGGTAAGCGGCAGGGTCGGAACAGGAGAGCGCACGAGGGAGCTTC
CAGGGGGAAACGCCTGGTATCTTTATAGTCCTGTCGGGTTTCGCCACCTCTGACTTGAGCGTCGAT
TTTTGTGATGCTCGTCAGGGGGGCGGAGCCTATGGAAAAACGCCAGCAACGCGGCCTTTTTACGGT
TCCTGGCCTTTTGCTGGCCTTTTGCTCACATGTTCTTTCCTGCGTTATCCCCTGATTCTGTGGATA
ACCGTATTACCGCCATGCAT
```

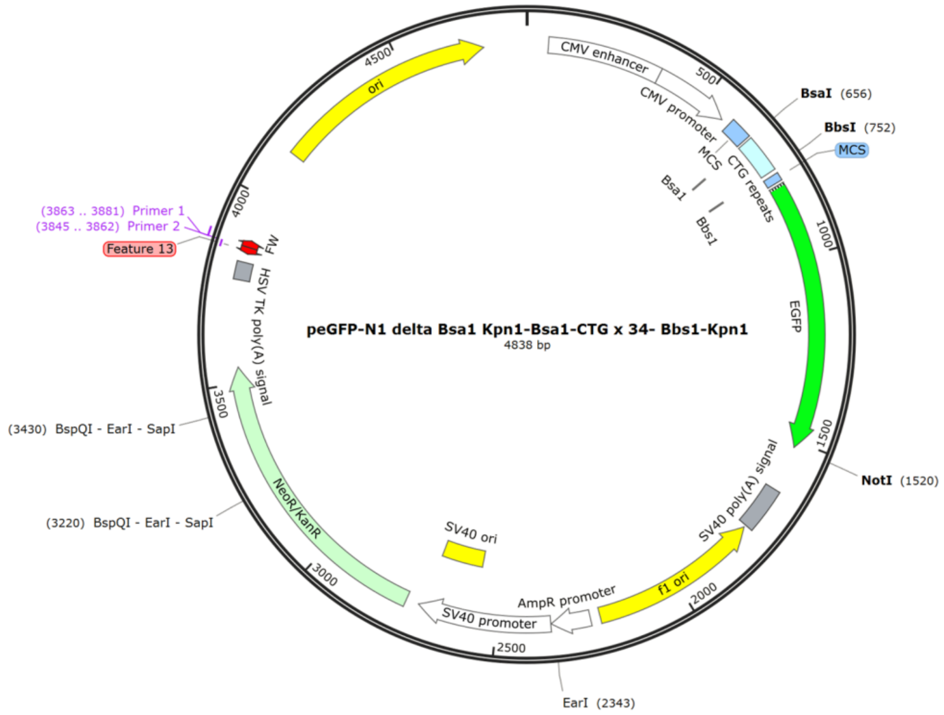

**Table S14.**

Sequence of DNA oligonucleotides used for assembly of ‘11’ RNA:DNA nanostructure used for the study of EGFP mRNA containing 34 CUG repeats (Figure 4c).

▲ Complementary DNA oligonucleotides ▲ Overhang for ‘1’ generation

| Oligo number | Sequence (5′→ 3′) |
|---|---|
| 1 | TTTTTTTTTTTTTTTTTTTTTTTTT |
| 2 | TTTTTTTTTTTTTTTTTTTTTTTTT |
| 3 | TTTTTTTTTTTTTTTTTTTTTTTTT |
| 4 | TGCAGTGAAAAAAATGCTTTATTTGTGAAATTT |
| **5** | GTGATGCTATTGCTTTATTTGTAACTTTGGATATCACTCATTAGTGGT |
| **6** | CATTATAAGCTGCAATAAACAAGTTTTTGGATATCACTCATTAGTGGT |
| 7 | AACAACAACAATTGCATTCATTTTATGTTTCAGGTTCAGG |
| 8 | GGGAGGTGTGGGAGGTTTTTTAAAGCAAGTAAAACCTCTA |
| 9 | CAAATGTGGTATGGCTGATTATGATCTAGAGTCGCGGCCG |
| 10 | CTTTACTTGTACAGCTCGTCCATGCCGAGAGTGATCCCGG |
| 11 | CGGCGGTCACGAACTCCAGCAGGACCATGTGATCGCGCTT |
| 12 | CTCGTTGGGGTCTTTGCTCAGGGCGGACTGGGTGCTCAGG |
| 13 | TAGTGGTTGTCGGGCAGCAGCACGGGGCCGTCGCCGATGG |
| 14 | GGGTGTTCTGCTGGTAGTGGTCGGCGAGCTGCACGCTGCC |
| 15 | GTCCTCGATGTTGTGGCGGATCTTGAA |
| 16 | GTTCACCTTGATGCCGTTCTTCTGCTTG |
| **17** | TCGGCCATGATATAGACGTTGTGGCTTTGGATATCACTCATTAGTGGT |
| **18** | TGTTGTAGTTGTACTCCAGCTTGTGTTTGGATATCACTCATTAGTGGT |
| 19 | CCCCAGGATGTTGCCGTCCTCCTTGAAGTCGATGCCCTTC |
| 20 | AGCTCGATGCGGTTCACCAGGGTGTCGCCCTCGAACTTCA |
| 21 | CCTCGGCGCGGGTCTTGTAGTTGCCGTCGTCCTTGAAGAA |
| 22 | GATGGTGCGCTCCTGGACGTAGCCTTCGGGCATGGCGGAC |
| 23 | TTGAAGAAGTCGTGCTGCTTCATGTGGTCGGGGTAGCGGC |
| 24 | TGAAGCACTGCACGCCGTAGGTCAGGGTGGTCACGAGGGT |
| 25 | GGGCCAGGGCACGGGCAGCTTGCCGGTGGTGCAGATGAAC |
| 26 | TTCAGGGTCAGCTTGCCGTAGGTGGCATCGCCCTCGCCCT |
| 27 | CGCCGGACACGCTGAACTTGTGGCCGTTTACGTCGCCGTC |
| 28 | CAGCTCGACCAGGATGGGCACCACCCCGGTGAACAGCTCC |
| 29 | TCGCCCTTGCTCACCATGGTGGCGACCG |
| 30 | GTGGATCCCGGGCCCGCGGTACGAAGACG |
| 31 | GAGACCGTACCGTCGACTGCAGAATTCGAAGCTTGAGCTC |
| 32 | GAGATCTGAGTCCGGTAGCGCTAGCGGATCTGAC |

#### Table S15.

Sequence of DNA oligonucleotides used for assembly of linear MS2 RNA nanostructure used for the optimization of RNA:DNA nanostructure assembly at low concentrations.

▲ Complementary DNA oligonucleotide

| Oligo number | Sequence (5′→ 3′) |
|---|---|
| 1 | TGGGTGGTAACTAGCCAAGCAGCTAGTTACCAAATCGGGAGAATC |
| 2 | CCGGGTCCTCTCTTTAGGGGGAGGTCCCTGGG |
| 3 | CCGAAGCCCGCCCACCTTTCGGTGGAGCCGG |
| 4 | ACCGCTTTCGCACCCGTGCTCTTTCGAGCACACCCACCCCGTTTACGGGG |
| 5 | GTCCCTCGGTCAGCTACCGAGGAGAGCTCGCT |
| 6 | GGCCCACACTCCTGAGGGAATGTGGG |
| 7 | AACCGGCGTTAGCCACTCCGAAGTGCGTATAACGCGCAC |
| 8 | GCCGGCGGACTTCATGCTGTCGGTGATTTCACCTCCAGTATGGAAC |
| 9 | CACGCTATGTAGCGACCACTGTCGTGCTTTTCGCTGAAGAACTTGCGTTCTCGAGCGATA |
| 10 | CGAGCAAGACGGAAACCCGAGGTACGGGTATCC |
| 11 | GCGAGCAGCCGCCCGTACGGAGTCTTGGTGTATACC |
| 12 | GAGACTGCCGTAGGCGGGCTGACTACG |
| 13 | TAGTAGTCGGCAGCGAGGTCCGTCCCACCGAAGAACATCGAA |
| 14 | GGCACCTGGGAGGAGAGCCGT |
| 15 | ACCCACACCTTATAGAGGCGTGGATCTGACATAC |
| 16 | CTCCGACAACTCCCCAACCCCGTAGCCGATTTAATATCAGCA |
| 17 | TCAGGGCGAAGAGATTGTCAACAGGTTTCTTGATGTAAAACGGTTTGACATCGAC |
| 18 | ACCACGGTAAAAGTGCGCGCCGCAG |
| 19 | CTCTCGCGAAAGAGCCCGGACACGAACGTTTTACGA |
| 20 | AGATTCGGTTTAAAACCGTAGTAGGCAAGTGCCTCTAGCACA |
| 21 | CGGGGTGCAATCTCACTGGGACA |
| 22 | TATAATATCGTCCCCGTAGATGCCTATGGTTCCGGCGTTACCAAAATGGATTT |
| 23 | GGGTCGCTTTGACTATTGCCCAGAATATCATGGACTCTAG |
| 24 | CTCAAATGTGAACCCATTTCCCATTGTGGAAAATAGTTCCCATCGTATCGTCTCGC |
| 25 | CATCTACGATTCCGTAGTGTGAGCGGATACGATCGAGATATGAA |
| 26 | TATAGCTCTGGTGGGAGAAAACTCCACA |
| 27 | CCAGGCGATCGGAGATGGAATCGGATG |
| 28 | CAGACGATAAGTCTATCGTCGCAAGCGAACCATCTACGCT |
| 29 | GCCCTGCTGAGCCAGACGCT |
| 30 | GGTTGATCGATTGATCATTCAGGTCTATACCAACGGATTTGAGCCGG |
| 31 | CGTCTGATGAAAGCACCGACCCCTTTCTGGA |
| 32 | GGTACATATTCATATCAGGCTCCTTACAGGCAGCCCGATCTATTTTATTATTCTTCGGAA |
| 33 | CTGTAAACACTCCGTTCCCTACAACGAGCCTAAATTCATATGACTCGTTATAGCGG |
| 34 | ACCGCGTGTCTGATCCACGGCGCAC |
| 35 | ATTGGTCTCGGACCAATAGAGCCGCTCTCAGAG |
| 36 | CGCGGGGGGTAACGGTTGCTTGTTCAGCGAA |
| 37 | CTTCTTGTAAGGCGCTGCATCCTGCAACTTGTGCCCCAT |

| | |
|---|---|
| 38 | AGGAGCACCGTTGGAGAACGTGCATTGC |
| 39 | CCAAACAACGACGATCGGTAGCCAGAGAG |
| 40 | GAGGTTGCCAATAAGGCTACGGATGC |
| 41 | TGGTTTGTAAAACATCCGGATCCCATGACAAGGATTTGTCATGTAAGAAA |
| 42 | CCTTCTCTATTTATCTGACCGCGATCACCATTCGCCTCCCGTAGCTTA |
| 43 | GCGATAGCTAAGGTACGACGGGTC |
| 44 | GCCTCGTCATTACCAGAACCTAAGGTCGGATGCTTT |
| 45 | GTGAGCAATTCGTCCCTTAAGTAAGCAATTGCTGTAAAGTCGTCACTGTG |
| 46 | CGGATCACCGCTTCCAGTAGCGACAGAA |
| 47 | GCAATTGATTGGTAAATTTCGAGAGAAAGATCGCGAGGAAGATCAAT |
| 48 | ACATAAAGAGTTGAACTTCTTTGTTGTCTTCGACATGGGTAATCCTCATGTTTGAATGG |
| 49 | CCGGCGTCTATTAGTAGATGCCGGAGTTTGCTG |
| 50 | CGATTGCTGAGGGAATCGGGTTTCCATCTTTTAGGAGA |
| 51 | CCTTGCATTGCCTTAACAATAAGCTCGCAGTCGGAATTCGTA |
| 52 | GCGAAAATTGGAATGGTTAGTTCCATATTTAAGTACGAACGCCATGCG |
| 53 | GCTACAGGAAGCTCTACACCACC |
| 54 | AACAGTCTGGGTTGCCACTTTAGGCACCTCGACTTT |
| 55 | GATGGTGTATTTGCGATTCTGCGCAGAGCTCTGACGA |
| 56 | ACGCTACAGGTTACTTTGTAAGCCTGTGAACGCGAGTTAGA |
| 57 | GCTGATCCATTCAGCGACCCCGTTAGCGAAGTTGCTTG |
| 58 | GGGCGACAGTCACGTCGCCAGTTCCGCCATT |
| 59 | GTCGACGAGAACGAACTGAGTAAAGTTAGAAGCCATGC |
| 60 | TTCAAACTCCGGTTGAGGGCTCTATCTAGAGAGCCGT |
| 61 | TGCCTGATTAATGCTAACGCATCTAAGGTATGGACCATCGAGAAA |
| 62 | GGAGACTTTACGTACGCGCCAGTTGTTGGCCATACGGATTGTAC |
| 63 | CCCTCGATGCATGGCTGAGATTTGGGCCTTAGCAG |
| 64 | TGCCCTGTCTCTCCACAGTCCACCCGTAGG |
| 65 | GAGCGTCAACGCTTATGATGGACTCACCCGTTATTACGTCAGTAACTGTTCCTGACATG |
| 66 | TAGGAGCATCCCACGGGGGC |
| 67 | CGTAAGGCCCTCGAGCATGTTACCTACAGGTAG |
| 68 | GAGCCAGTCGACAACGAATGAGAAAGGCACCTTTT |
| 69 | CCCACACTATACCTAGTGGGTTCAAGATACCTAGAG |
| 70 | ACGACAACCATGCCAAACGTGCATCGTTTATGTAAAACCATATCACGATA |
| 71 | CGTCGCGATATGTTGCACGTTGTCTGGAAGTTTGCAGCTGGATA |
| 72 | CGACAGACGGCCATCTAACTTGATGTTAGTACCGAC |
| 73 | CTGACGTACGGCTCTCATAGGAAGAAACTCTTGAAG |
| 74 | GTGAACCTTCGTAAGCATCTCATATGCACCCTGGATATCACTC |
| 75 | ATTAGTGGTAACCAACCGAACTGCAACTCCAACCA |
| 76 | CCTGCCGGCCACGTGTTTTGATCGAAACTTTCGATCTTCGTTTAGGGCAAGG |
| 77 | TAGCGGAGCGCCTGGCGCCAA |
| 78 | TTACCGCGACGAGCGGCAGTGTACGCC |
| 79 | TTCACGAGCGCAATGGTTTGCGTCGCGAGTTGTGAGGCTGTCGA |
| 80 | CCTGGCCTCTGCTAAAGCAACACCAAGGT |
| 81 | TAAAATTACCCTGGGTGACCTTTTGCAGGACTTCGGTCGACGCC |
| 82 | CGGTTCGCAACGTTCTGCGGCACTTCGATGTAAGTCAAGTTTTGGCTTACAGGGAAG |
| 83 | AGGCTGTAGCAGGAGCGTGCGTCGAG |

| | |
|---|---|
| **84** | GGAGAAGCCGAAACCGGCTTTCTCCTCGTACGG |
| **85** | GCGACCCCACGATGACCCACTTCGC |
| **86** | TTGTAGGCACCTTGATCTATCGATGTGACACTTAACGCCCCC |
| **87** | CGTGAATACGGAGAGGGGTAGTGCCACTGTTTCG |
| **88** | TTTTGGCCCCAGTCGAGTTAAAACGACCGGGAGTCCAGTTCGAACGATATTTTAAAGAG |
| **89** | AATGAGTTATCTTCAGTCTCACCGTCCGCGTAAACGCGAACGGAG |
| **90** | GGGACGAAGGTCTCGTTCTCCCTATCAAGGGTACTAAAAG |
| **91** | CTCGCACAGGTCAAACCTCCTAGGAATGGAATTCCG |
| **92** | GCTACCTACAGCGATAGCCATGGTAGCGTCTCG |
| **93** | CTAAAGACATTAAAAATGGCATTAGCTCGACAGGAAGTTGAGCAGGA |
| **94** | CCCCGAAAGGGGTCCCACCC |

**Table S16.**

Sequence of DNA oligonucleotides used for assembly of 'PPP' MS2 RNA:DNA nanostructure used for detection of nanostructures in total RNA background.

▲ Complementary DNA oligonucleotide ▲ Overhang for '1' generation

| Oligo number | Sequence (5'→ 3') |
|---|---|
| 1 | TGGGTGGTAACTAGCCAAGCAGCTAGTTACCAAATCGGGA |
| 2 | GAATCCCGGGTCCTCTCTTTAGGGGGAGGTCCCTGGGCCG |
| 3 | AAGCCCGCCCACCTTTCGGTGGAGCCGGACCGCTTTCGCA |
| 4 | CCCGTGCTCTTTCGAGCACACCCACCCCGTTTACGGGGGT |
| 5 | CCCTCGGTCAGCTACCGAGGAGAGCTCGCTGGCCCACACT |
| 6 | CCTGAGGGAATGTGGGAACCGGCGTTAGCCACTCCGAAGT |
| 7 | GCGTATAACGCGCACGCCGGCGGACTTCATGCTGTCGGTG |
| 8 | ATTTCACCTCCAGTATGGAACCACGCTATGTAGCGACCAC |
| 9 | TGTCGTGCTTTTCGCTGAAGAACTTGCGTTCTCGAGCGAT |
| 10 | ACGAGCAAGACGGAAACCCGAGGTACGGGTATCCGCGAGC |
| 11 | AGCCGCCCGTACGGAGTCTTGGTGTATACCGAGACTGCCG |
| 12 | TAGGCGGGCTGACTACGTAGTAGTCGGCAGCGAGGTCCGT |
| 13 | CCCACCGAAGAACATCGAAGGCACCTGGGAGGAGAGCCGT |
| 14 | ACCCACACCTTATAGAGGCGTGGATCTGACATACCTCCGA |
| 15 | CAACTCCCCAACCCCGTAGCCGATTTAATATCAGCATCAG |
| 16 | GGCGAAGAGATTGTCAACAGGTTTCTTGATGTAAAACGGT |
| 17 | TTGACATCGACACCACGGTAAAAGTGCGCGCCGCAGCTCT |
| 18 | CGCGAAAGAGCCCGGACACGAACGTTTTACGAAGATTCGG |
| 19 | TTTAAAACCGTAGTAGGCAAGTGCCTCTAGCACACGGGGT |
| 20 | GCAATCTCACTGGGACATATAATATCGTCCCCGTAGATGC |
| 21 | CTATGGTTCCGGCGTTACCAAAATGGATTTGGGTCGCTTT |
| 22 | GACTATTGCCCAGAATATCATGGACTCTAGCTCAAATGTG |
| 23 | AACCCATTTCCCATTGTGGAAAATAGTTCCCATCGTATCG |
| 24 | TCTCGCCATCTACGATTCCGTAGTGTGAGCGGATACGATC |
| 25 | GAGATATGAATATAGCTCTGGTGGGAGAAAACTCCACACC |
| 26 | AGGCGATCGGAGATGGAATCGGATGCAGACGATAAGTCTA |
| 27 | TCGTCGCAAGCGAACCATCTAC |
| 28 | GCTGCCCTGCTGAGCCAGACGCT |
| 29 | GGTTGATCGATTGATCATTCAGGTCtttggatatcactcattagtggt |
| 30 | TATACCAACGGATTTGAGCCGGCGTtttggatatcactcattagtggt |
| 31 | CTGATGAAAGCACCGACCCCTTTCTtttggatatcactcattagtggt |
| 32 | GGAGGTACATATTCATATCAGGCTCCTTACAGGCAGCCCG |
| 33 | ATCTATTTTATTATTCTTCGGAACTGTAAACACTCCGTTC |
| 34 | CCTACAACGAGCCTAAATTCATATGACTCGTTATAGCGGA |
| 35 | CCGCGTGTCTGATCCACGGCGCACATTGGTCTCGGACCAA |
| 36 | TAGAGCCGCTCTCAGAGCGCGGGGGGTAACGGTTGCTTGT |
| 37 | TCAGCGAACTTCTTGTAAGGCGCTGCATCCTGCAACTTGT |

| | |
|---|---|
| 38 | GCCCCATAGGAGCACCGTTGGAGAACGTGCATTGCCCAAA |
| 39 | CAACGACGATCGGTAGCCAGAGAGGAGGTTGCCAATAAGG |
| 40 | CTACGGATGCTGGTTTGTAAAACATCCGGATCCCATGACA |
| 41 | AGGATTTGTCATGTAAGAAACCTTCTCTATTTATCTGACC |
| 42 | GCGATCACCATTCGCCTCCCGTAGCTTAGCGATAGCTAAG |
| 43 | GTACGACGGGTCGCCTCGTCATTACCAGAACCTAAGGTCG |
| 44 | GATGCTTTGTGAGCAATTCGTCCCTTAAGTAAGCAATTGC |
| 45 | TGTAAAGTCGTCACTGTGCGGATCACC |
| 46 | GCTTCCAGTAGCGACAGAAGCAATTGAT |
| 47 | TGGTAAATTTCGAGAGAAAGATCGCtttggatatcactcattagtggt |
| 48 | GAGGAAGATCAATACATAAAGAGTTtttggatatcactcattagtggt |
| 49 | GAACTTCTTTGTTGTCTTCGACATGtttggatatcactcattagtggt |
| 50 | GGTAATCCTCATGTTTGAATGGCCGGCGTCTATTAGTAGA |
| 51 | TGCCGGAGTTTGCTGCGATTGCTGAGGGAATCGGGTTTCC |
| 52 | ATCTTTTAGGAGACCTTGCATTGCCTTAACAATAAGCTCG |
| 53 | CAGTCGGAATTCGTAGCGAAAATTGGAATGGTTAGTTCCA |
| 54 | TATTTAAGTACGAACGCCATGCGGCTACAGGAAGCTCTAC |
| 55 | ACCACCAACAGTCTGGGTTGCCACTTTAGGCACCTCGACT |
| 56 | TTGATGGTGTATTTGCGATTCTGCGCAGAGCTCTGACGAA |
| 57 | CGCTACAGGTTACTTTGTAAGCCTGTGAACGCGAGTTAGA |
| 58 | GCTGATCCATTCAGCGACCCCGTTAGCGAAGTTGCTTGGG |
| 59 | GCGACAGTCACGTCGCCAGTTCCGCCATTGTCGACGAGAA |
| 60 | CGAACTGAGTAAAGTTAGAAGCCATGCTTCAAACTCCGGT |
| 61 | TGAGGGCTCTATCTAGAGAGCCGTTGCCTGATTAATGCTA |
| 62 | ACGCATCTAAGGTATGGACCATCGAGAAAGGAGACTTTAC |
| 63 | GTACGCGCCAGTTGTTGGCCATACGGA |
| 64 | TTGTACCCCTCGATGCATGGCTGAGATT |
| 65 | TGGGCCTTAGCAGTGCCCTGTCTCTtttggatatcactcattagtggt |
| 66 | CCACAGTCCACCCGTAGGGAGCGTCtttggatatcactcattagtggt |
| 67 | AACGCTTATGATGGACTCACCCGTTtttggatatcactcattagtggt |
| 68 | ATTACGTCAGTAACTGTTCCTGACATGTAGGAGCATCCCA |
| 69 | CGGGGGCCGTAAGGCCCTCGAGCATGTTACCTACAGGTAG |
| 70 | GAGCCAGTCGACAACGAATGAGAAAGGCACCTTTTCCCAC |
| 71 | ACTATACCTAGTGGGTTCAAGATACCTAGAGACGACAACC |
| 72 | ATGCCAAACGTGCATCGTTTATGTAAAACCATATCACGAT |
| 73 | ACGTCGCGATATGTTGCACGTTGTCTGGAAGTTTGCAGCT |
| 74 | GGATACGACAGACGGCCATCTAACTTGATGTTAGTACCGA |
| 75 | CCTGACGTACGGCTCTCATAGGAAGAAACTCTTGAAGGTG |
| 76 | AACCTTCGTAAGCATCTCATATGCACCCTGGATATCACTC |
| 77 | ATTAGTGGTAACCAACCGAACTGCAACTCCAACCACCTGC |
| 78 | CGGCCACGTGTTTTGATCGAAACTTTCGATCTTCGTTTAG |
| 79 | GGCAAGGTAGCGGAGCGCCTGGCGCCAATTACCGCGACGA |
| 80 | GCGGCAGTGTACGCCTTCACGAGCGCAATGGTTTGCGTCG |
| 81 | CGAGTTGTGAGGCTGTCGACCTGGCCTCTGCTAAAGCAAC |
| 82 | ACCAAGGTTAAAATTACCCTGGGTGACCTTTTGCAGGACT |
| 83 | TCGGTCGACGCCCGGTTCGCAACGTTCTGCGGCACTTCGA |

| | |
|---|---|
| 84 | TGTAAGTCAAGTTTTGGCTTACAGGGAAGAGGCTGTAGCA |
| 85 | GGAGCGTGCGTCGAGGGAGAAGCCGAAACCGGCTTTCTCC |
| 86 | TCGTACGGGCGACCCCACGATGACCCACTTCGCTTGTAGG |
| 87 | CACCTTGATCTATCGATGTGACACTTAACGCCCCCCGTGA |
| 88 | ATACGGAGAGGGGTAGTGCCACTGTTTCGTTTTGGCCCCA |
| 89 | GTCGAGTTAAAACGACCGGGAGTCCAGTTCGAACGATATT |
| 90 | TTAAAGAGAATGAGTTATCTTCAGTCTCACCGTCCGCGTA |
| 91 | AACGCGAACGGAGGGGACGAAGGTCTCGTTCTCCCTATCA |
| 92 | AGGGTACTAAAAGCTCGCACAGGTCAAACCTCCTAGGAAT |
| 93 | GGAATTCCGGCTACCTACAGCGATAGCCATGGTAGCGTCT |
| 94 | CGCTAAAGACATTAAAAATGGCATTAGCTCGACAGGAAGT |
| 95 | TGAGCAGGACCCCGAAAGGGGTCCCACCC |

#### Table S17.

Sanger sequencing of subcloned region holding CTG and CCTG tandem repeats in modified pJET1.2/blunt cloning vector (CloneJET PCR Cloning Kit, Thermo Fischer, Catalog number: K1231). Primers pJET1.2 F and R were used.

12 CTG

| **Sequence (5′→ 3′)** |
|---|
| CCGCGRRAWMTTCGGGAWGGCTCGAGTTTTTCAGCAGATCATGGTGCAGTGTAGCCGGGAATG**CTG**<br>**CTGCTGCTGCTGCTGCTGCTGCTGCTGCTGCTG**GGGGGATCACAGACCATTCTCGGCTTAAATCTT<br>TCTAGAAGATCTCCTACAATATTCTCAGCTGCCATGGAAAATCGATGTTCTTCTTTTATTCTCTCA<br>AGATTTTCAGGCTGTATATTAAAACTTATATTAAGAACTATGCTAACCACCTCATCAGGAACCGTT<br>GTAGGTGGCGTGGGTTTTCTTGGCAATCGACTCTCATGAAAACTACGAGCTAAATATTCAATATGT<br>TCCTCTTGACCAACTTTATTCTGCATTTTTTTTGAACGAGGTTTAGAGCAAGCTTCAGGAAACTGA<br>GACAGGAATTTTATTAAAAATTTAAATTTTGAAGAAAGTTCAGGGTTAATAGCATCCATTTTTTGC<br>TTTGCAAGTTCCTCAGCATTCTTAACAAAAGACGTCTCTTTTGACATGTTTAAAGTTTAAACCTCC<br>TGTGTGAAATTATTATCCGCTCATAATTCCACACATTATACGAGCCGGAAGCATAAAGTGTAAAGC<br>CTGGGGTGCCTAATGAGTGAGCTAACTCACATTAATTGCGTTGCGCTCACTGCCAATTGCTTTCCA<br>GTCGGGAAACCTGTCGTGCCAGCTGCATTAATGGAATCGGGCCAACSCSCCGGAAAGCCGTTTGSC<br>GTATTGGGCGCTCTTCCGCTTCCCTCSCTCAMTGACTCGCTGGCGCTCGGTCGTCKGGCTGGCGGC<br>GAGCGGTATCAGCTCACTGAAGCGTATACGTTATCCMAGATTCRGGGGATAAACGC |

24 CTG

| **Sequence (5′→ 3′)** |
|---|
| CCGSGRAAWCYTTCGGGATGGCTCGAGTTTTTCAGCAGATCATGGTGCAGTGTAGCCGGGAATG**CT**<br>**GCTGCTGCTGCTGCTGCTGCTGCTGCTGCTGCTGCTGCTGCTGCTGCTGCTGCTGCTGCTGCTGCT**<br>**GCTG**GGGGGATCACAGACCATTCTCGGCTTAAATCTTTCTAGAAGATCTCCTACAATATTCTCAGC<br>TGCCATGGAAAATCGATGTTCTTCTTTTATTCTCTCAAGATTTTCAGGCTGTATATTAAAACTTAT<br>ATTAAGAACTATGCTAACCACCTCATCAGGAACCGTTGTAGGTGGCGTGGGTTTTCTTGGCAATCG<br>ACTCTCATGAAAACTACGAGCTAAATATTCAATATGTTCCTCTTGACCAACTTTATTCTGCATTTT<br>TTTTGAACGAGGTTTAGAGCAAGCTTCAGGAAACTGAGACAGGAATTTTATTAAAAATTTAAATTT<br>TGAAGAAAGTTCAGGGTTAATAGCATCCATTTTTTGCTTTGCAAGTTCCTCAGCATTCTTAACAAA<br>AGACGTCTCTTTTGACATGTTTAAAGTTTAAACCTCCTGTGTGAAATTATTATCCGCTCATAATTC<br>CACACATTATACGAGCCGGAAGCATAAAGTGTAAAGCCTGGGGTGCCTAATGAGTGAGCTAACTCA<br>CATTAATTGCGTTGCGCTCACTGCCAATTGCTTTCAGTCGGGAACCTGTCGTGCCAGCTGCATATG<br>AATCGTCAACGCGCGGARAGCGTTGCGTATGGGCGCTCTTCGCTCTYSTYCACTGACTCGCTGCGC<br>TCGGTCGATCGTCTGCGCAGCCGGTATCAGCTTCCACCTCC |

30 CTG

| Sequence (5′→ 3′) |
| --- |
| GCGCRRRTWWCTTCCGGGAGGCTCGAGTTTTTCAGCAAGATCATGGTGCAGTGTAGCCGGGAATG**C** |
| **TGCTGCTGCTGCTGCTGCTGCTGCTGCTGCTGCTGCTGCTGCTGCTGCTGCTGCTGCTGCTGCTGC** |
| **TGCTGCTGCTGCTGCTGCTGCTG**GGGGGATCACAGACCATTCTCGGCTTAAATCTTTCTAGAAGAT |
| CTCCTACAATATTCTCAGCTGCCATGGAAAATCGATGTTCTTCTTTTATTCTCTCAAGATTTTCAG |
| GCTGTATATTAAAACTTATATTAAGAACTATGCTAACCACCTCATCAGGAACCGTTGTAGGTGGCG |
| TGGGTTTTCTTGGCAATCGACTCTCATGAAAACTACGAGCTAAATATTCAATATGTTCCTCTTGAC |
| CAACTTTATTCTGCATTTTTTTTGAACGAGGTTTAGAGCAAGCTTCAGGAAACTGAGACAGGAATT |
| TTATTAAAAATTTAAATTTTGAAGAAAGTTCAGGGTTAATAGCATCCATTTTTTGCTTTGCAAGTT |
| CCTCAGCATTCTTAACAAAAGACGTCTCTTTTGACATGTTTAAAGTTTAAACCTCCTGTGTGAAAT |
| TATTATCCGCTCATAATTCCACACATTATACGAGCCGGAAGCATAAAGTGTAAGCCTGGGGTGCCT |
| AATGAGTGAGCTAACTCACATTAATTGCGTTGCGCTCACTGCCAATTGCTTTCCAGTCGGCAAKCT |
| GTCGTGTCTAGCTGYATTATGAATCGTAACGCSCGGGGGAAGAAGCGTTGCGTATTGGASCTCTTC |
| GCTCTCGCTCMTGACTCGCTGSGCTCGGCTCGATCGGACTG |

51 CTG

| Sequence (5′→ 3′) |
| --- |
| CCGGCARRWWMWTTCGGGAAGGCTCGARTTTTTCAGCAAGATCATGGTGCAGTGTAGCCGGGWATG |
| **CTGCTGCTGCTGCTGCTGCTGCTGCTGCTGCTGCTGCTGCTGCTGCTGCTGCTGCTGCTGCTGCTG** |
| **CTGCTGCTGCTGCTGCTGCTGCTGCTGCTGCTGCTGCTGCTGCTGCTGCTGCTGCTGCTGCTGCTG** |
| **CTGCTGCTGCTGCTGCTGCTG**GGGGGATCACAGACCATTCTCGGCTTAAWTCTTTCTAGAASATCT |
| CCTACAATATTCTCAGCTGCCATGGAAAATCGATGTTCTTCTTTTATTCTCTCAWGATTTTCAGGC |
| TGTATATTAAWACTWATATTAAGAACTATGCTAACCACCTCATCAGGAACCGTTGTAGGTGGCGTG |
| GGTTTTCTTGGCAATCGACTCTCATGAAAACTACGAGCTAAATATTCAATATGTTCCTCTTGACCA |
| WCTTTATTCTGCATTTTTTTTGAACGAGGWTTAGAGCAWGCTTCAGGAAACTGAGACAGGAATTTT |
| ATTAAAAATTTWAATTTTGAAGAAAGTTCAGGGTTAATAGCATCCWTTTTTTGCTTTGCAASTTCC |
| TCAGCATTCWTAACAAAAGACGTCTCTTTTTGACATGTTAAAGTATAAACCTSCTGTGTGAAATTA |
| TTATCGCTCATAATTCCACACATTATACGAGCCAGCAGCATAAAGTGYTARAGGCCTGGGGGTGAC |
| TAATGAGTGAGCTAACTCACATGATTGCGTTGCGCTCGACTGACAATTGGCTTTCCARTCCRGCAT |
| SCCTGGTCGTGCCAGCTTGCATTGAATGGAATCGGACTACCGGCTCGAAGAGG |

57 CCTG

| Sequence (5′→ 3′) |
| --- |
| CCGSRRRTWMTTCGGAWGGCTCGAGTTTTTCAGCAGATGGCCTTATAACCATGCAATGTGTCCATT |
| AAGTTGGACTTGGAATGAGTGAATGAGTATTACTGCCAGTGTGTGTGTGTGTGTGTGTGTGTGTGT |

GTGTGTGTGTGTGTCTGTCTGTCTGTCTGTCTGTCTGTCTGTCTG**CCTGCCTGCCTGCCTGCCTGC**
**CTGCCTGCCTGCCTGCCTGCCTGCCTGMCTGCCTGCCTGCCTGCCTGCCTGCCTGCCTGCCTGCCT**
**GCCTGCCTGCCTGCCTGCCTGCCTGCCTGCCTGCCTGCCTGCCTGCCTGCCTGCCTGCCTGCCTGC**
**CTGCCTGCCTGCCTGCCTGCCTGCCTGCCTGCCTGCCTGCCTGCCTGCCTGCCTGCCTGCCTGCCT**
**GCCTGCCTG**WCTGTCTGKCTCACTTTGTCCCCYAAGCTGAAKTGCATGGAATTATCACGGGTCTAT
CWTTCTAGAAG

#### Table S18.

Sanger sequencing of subcloned region holding GCN tandem repeats in modified pJET1.2/blunt cloning vector (CloneJET PCR Cloning Kit, Thermo Fischer, Catalog number: K1231). Primers pJET1.2 F and R were used.

20 GCN

| **Sequence (5′→ 3′)** |
|---|
| AKSSSSSRRGGGCCACTCSRAWCTCGGWGGCTCGAGTTTTTCAGCAGATCATGGGGAGGAGGGTGT<br>TAAAACAAGCCGAAGTAGAACTTGGGCCACCCTAACCGGTGCTTTTCTTTCCCATTTTCTTCTTTC<br>TCCCCCTGCTTCACCGTCTCTCCTTCCGTCTTGGGCCAGGTGTGGTTCCAGAACCGCCGCGCCAAG<br>TTTCGCAAGCAGGAGCGCGCAGCGGCAGCCGCAGCGGCCGCGGCCAAGAACGGCTCCTCGGGCAAA<br>AAGTCTGACTCTTCCAGGGACGACGAGAGCAAAGAGGCCAAGAGCACTGACCCGGACAGCACTGGG<br>GGCCCAGGTCCCAATCCCAACCCCACCCCCAGCTGCGGGGCGAATGGAGGCGGCGGCGGCGGGCCC<br>AGCCCGGCTGGAGCTCCGGGGGCGGCGGGGCCCGGGGGCCCGGGAGGCGAACCCGGCAAGGGCGGC<br>**GCAGCAGCAGCGGCGGCGGCCGCGGCAGCGGCGGCGGCAGCGGCAGCGGCGGCAGCT**GGAGGC<br>CTGGCTGCGGCTGGGGGCCCTGGACAAGGCTGGGCTCCCGGCCCCGGCCCCATCACCTCCATCCCG<br>GATTCGCTTGGGGGTCCCTTCGCCAGCGTCCTATCTTCGCTCCAAGACCCAACGGTGCCAAAGCCG<br>CCTTAGTGAAGAGCAGTATGTTCTGATCTGGAATCCTGCGCGCGCGCGCGACAGCGGGCGGAGC<br>CAGGACYCGGCGGGCGAGTGGGCGAGCGGGTAGCCCAGCTATGTCGTCGCTGATCTTTTCTAGAGA<br>TCTCTACAATATTTCTCAGCTGCCATGAAATCGAGGTCCTTCGTTCATTCYCYCARAATTTTCCAG |

26 GCN

| **Sequence (5′→ 3′)** |
|---|
| TCGSSRRWWMMTTCGGGAWGGCTCGAGTTTTTCAGCAAGATCATGGGGAGGAGGGTGTTAAAACAA<br>GCCGAAGTAGAACTTGGGCCACCCTAACCGGTGCTTTTCTTTCCCATTTTCTTCTTTCTCCCCCTG<br>CTTCACCGTCTCTCCTTCCGTCTTGGGCCAGGTGTGGTTCCAGAACCGCCGCGCCAAGTTTCGCAA<br>GCAGGAGCGCGCAGCGGCAGCCGCAGCGGCCGCGGCCAAGAACGGCTCCTCGGGCAAAAAGTCTGA<br>CTCTTCCAGGGACGACGAGAGCAAAGAGGCCAAGAGCACTGACCCGGACAGCACTGGGGGCCCAGG<br>TCCCAATCCCAACCCCACCCCCAGCTGCGGGGCGAATGGAGGCGGCGGCGGCGGGCCCAGCCCGGC<br>TGGAGCTCCGGGGGCGGCGGGGCCCGGGGGCCCGGGAGGCGAACCCGGCAAGGGCGGC**GCAGCAGC**<br>**AGCGGCGGCGGCCGCGGCAGCGGCGGCGGCCGCGGCAGCGGCGGCGGCGGCAGCGGCAGCGGCGGC**<br>**AGCT**GGAGGCCTGGCTGCGGCTGGGGGYCCTGGACAAGGCTGGGCTCYCGGYCCCGGCYCCATCAC<br>CTCCATCCCGGATTCGCTTGGGGGTCCCTTTCGCCAGCGTCCTATCTTCGCTCCAAAGACYCAACG<br>GTGCCAAAGYCGCCTTAGTGAAGAGCAGTWTGTTYCTGATYTGGAATCCTGCGGCGGCGGCGCGGC<br>GGCGACAGCGGKCGAGYCAGGGTCYCGGTYGGGCGARTTGGGYCGAGYCGGTAGCYCAGGCTTATT<br>KGTCGTTCGCTGAATCTTTTCYTAGAAGATCTCCTACAAATATTTCTCARGCTTGCCCWKGGGAAA<br>A |

#### Table S19.

Sanger sequencing of subcloned region with and without CTG tandem repeats in modified pEGFP cloning vector:

0 CTG

| Sequence (5′→ 3′) |
|---|
| GCKKKKRWGGGGGRRGGSYTAAAWARSMRAGYTGGTTTAGTGAACCGTCAGAKCCGCTAGCGCTAC<br>CGGACTCAGATCTCGAGCTCAGCTTCGAATTCTGCAGTCGACGGTACCGCGGGCCCGGGATCCACC<br>GGTCGCCACCATGGTGAGCAAGGGCGAGGAGCTGTTCACCGGGGTGGTGCCCATCCTGGTCGAGCT<br>GGACGGCGACGTAAACGGCCACAAGTTCAGCGTGTCCGGCGAGGGCGAGGGCGATGCCACCTACGG<br>CAAGCTGACCCTGAAGTTCATCTGCACCACCGGCAAGCTGCCCGTGCCCTGGCCCACCCTCGTGAC<br>CACCCTGACCTACGGCGTGCWGTGCTTCAGCCGCTACCCCGACCACATGAAGCAGCACGACTTCTT<br>CAAGTCCGCCATGCCCGAAGGCTACGTCCAGGAGCGCACCATCTTCTTCAAGGACGACGGCAACTA<br>CAWGACCCGCGCCGAGGTGAAGKTCGAGGGCGACACCCTGGTGAACCGCATCGAGCTGAAGGGCAT<br>CGACTTCARGGAGGACGGCAACATCCTGGGGGSACAAGCTGGAGTACAACTACAACAGCCACACGT<br>YTATATCATGGCCGACAAGCAGAAGAACGGCATCAAGGKGAACTTCAARATCCGGCCACAACATCG<br>AGGGACGGMAGCGTGGARGCTCGCCGACCMACTACCAGYAGAACMCCCCCATCGGGCGACGGCCCC<br>GTGCTGCTGCCCGAMAACCACTACCTGGAGCACCCMGTCCGCCCTGAKCAAAGAMCCCAACGARAA<br>GSGSGAWYCMCMTGGTCCCTGCTKGAWTCYGTGGACCGCCCGCCCGGKAWMYMCYTCTCGSGATTG<br>GRA |

34 CTG

| Sequence (5′→ 3′) |
|---|
| TKWWKGGGAGRWKYWWAWAARCARAGCTGGTTTAGTGACCGGTCAAGATCCGGCTRGCGCTACCGG<br>ACTCAGATCTCGAGCTCAAGCTTCGAATTCTGCAGTCGACGGTACGGTCTC**CTGCTGCTGCTGCTG**<br>**CTGCTGCTGCTGCTGCTGCTGCTGCTGCTGCTGCTGCTGCTGCTGCTGCTGCTGCTGCTGCTGCTG**<br>**CTGCTGCTGCTGCTGCTGCTG**CGTCTTCGTACCGCGGGCCCGGGATCCACCGGTCGCCACCATGGT<br>GAGCAAGGGCGAGGAGCTGTTCACCGGGGTGGTGCCCATCCTGGTCGAGCTGGACGGCGACGTAAA<br>CGGCCACAAGTTCAGCGTGTCCGGCGAGGGCGAGGGCGATGCCACCTACGGCAAGCTGACCCTGAA<br>GTTCATCTGCACCACCGGCAAGCTGCCCGTGCCCTGGCCCACCCTCGTGACCACCCTGACCTACGG<br>CGTGCAGTGCTTCAGCCGCTACCCCGACCACATGAAGCAGCACGACTTCTTCAAGTCCGCCATGCC<br>CGAAGGCTACGTCCAGGAGCGCACCATCTTCTTCAAGGACGACGGCAACTACAAGACCCGCGCCGA<br>GGTGAAGTTCGAGGGCGACACCCTGGTGAACCGCATCGAGCTGAAGGGCATCGACTTCAAGGAGGA<br>CGGCAACATCCTGGGGCACAAGCTGGAGTACAACTACAACAGCCACAACGTCTATATCATGGCCG<br>ACAAGCAGGAGAACGGCATCAGGTGAACTTTCAAGATCCGCCWCACATCGAAGACGGCAGCGTGGC<br>AGCTCGCCGACAMTACCRGCRAACAYCCCGATCGCGAACGGTCCGTGACTGCTGACCGACAACYAC<br>TTACC |

#### Table S20.

Representative measurement durations and counts of linear RNA:DNA nanostructures detected.

| Figure | Sample | Experiment duration (h) | Unfolded events | Number of nanopores | Measurement termination |
|---|---|---|---|---|---|
| **1** | 51 CUG and 57 CCUG RNA:DNA nanostructure | ~2 | ~100 | 1 | Measurement was stopped |
| **2** | 12, 24, 30, 51 CUG RNA:DNA nanostructure mix | ~6 | ~400 | 1 | Measurement was stopped |
| **3** | 20 and 26 GCN RNA:DNA nanostructure mix | ~4 | ~120 | 1 | Measurement was stopped |
| **4** | mRNA and 34 CUG mRNA RNA:DNA nanostructure | ~1 to 8 | ~10 | 3 | Nanopore clogging |
| **S32** | MS2 nanostructure | ~1 | ~20 | 1 | Nanopore clogging |

**Table S21.**

The table includes the IV curves of the nanopores used to characterize the RNA:DNA nanostructure prior to the measurement. The ionic current and RMS noise at 600 mV is presented for each pore presented in the Main text.

| Pore number | Sample | Current, RMS noise at 600 mV | Current / voltage curve (IV curve) |
|---|---|---|---|
| 1 | '1111' CUG RNA:DNA nanostructure (Figure 1d) | 9.5 nA<br>6.1 pA | 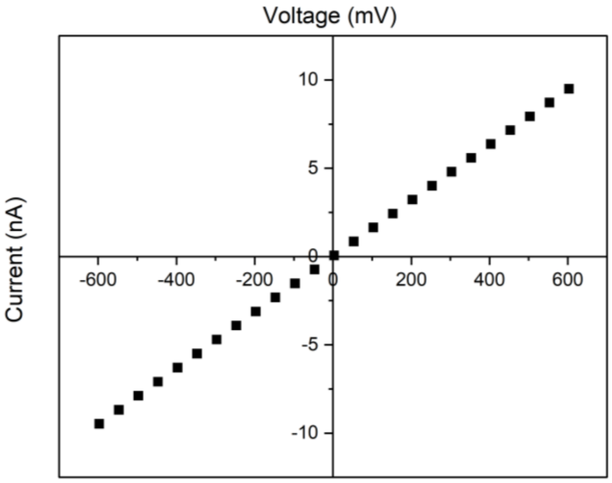 |
| 2 | '1001' CCUG RNA:DNA nanostructure (Figure 1e) | 8.4 nA<br>5.8 pA | 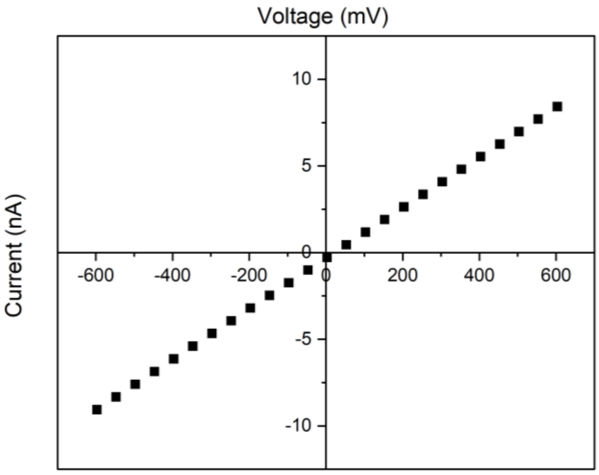 |

| | | | |
|---|---|---|---|
| 3 | RNA:DNA nanostructures for counting CUG repeats (Figure 2) | 10.1 nA<br>6.5 pA | 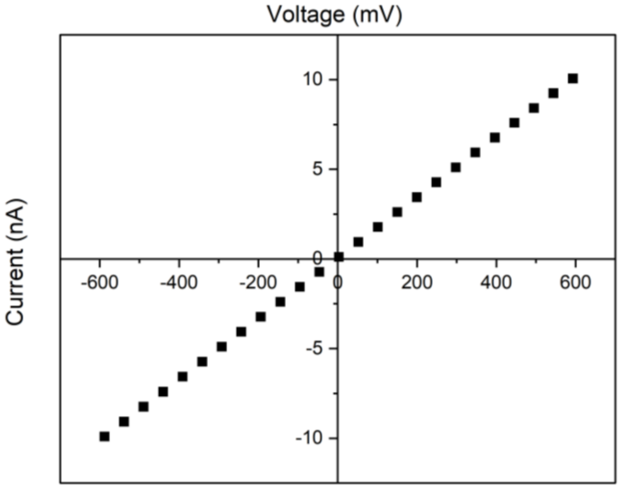 |
| 4 | GCN RNA:DNA nanostructures (Figure 3) | 9.0 nA<br>5.7 pA | 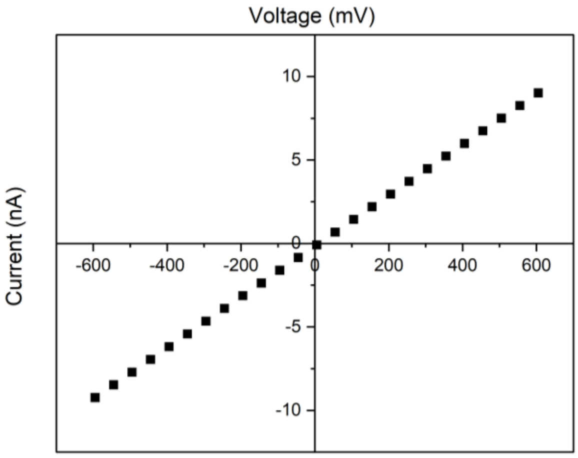 |
| 5 | wild type mRNA nanostructure (Figure 4) | 12.6 nA<br>6.6 pA | 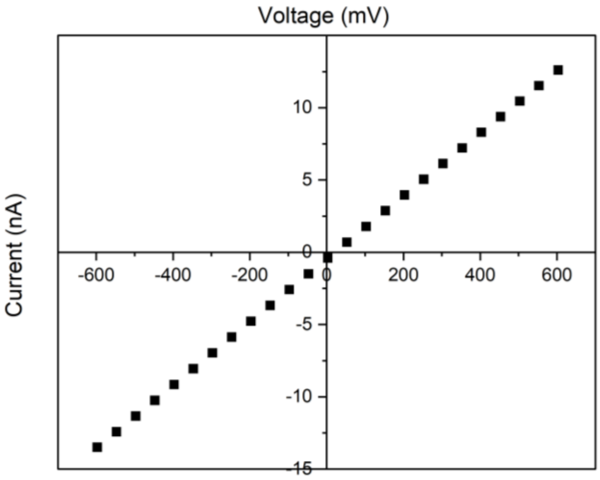 |

| | | | |
|---|---|---|---|
| 6 | CUG mRNA nanostructure (Figure 4) | 7.3 nA<br>5.4 pA | 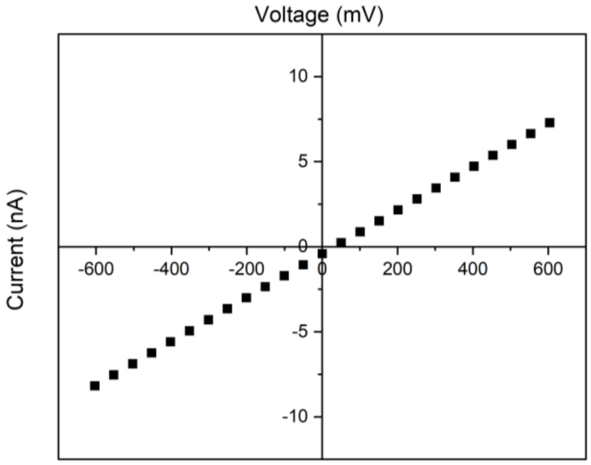 |

#### Table S22.

Tukey's Honest Significant Difference test (HSD). Distributions are labelled as $T_n$, where *n* correspond to the repeat array length. *Q* represents the critical value obtained from the Studentized Range Distribution table and *p* the p-value.

HSD procedure for data in Figure 2d (spike depth) and data in Figure S17 (spike area)

| Figure 2d (spike depth) | | |
|---|---|---|
| **Pairwise Comparisons** | **$HSD_{0.05}$ = 0.0435**<br>**$HSD_{0.01}$ = 0.0529** | **$Q_{0.05}$ = 3.6482**<br>**$Q_{0.01}$ = 4.4303** |
| **$T_{12}$:$T_{24}$** | 0.27 | Q = 22.34 ($p < 0.00001$) |
| **$T_{12}$:$T_{30}$** | 0.34 | Q = 28.83 ($p < 0.00001$) |
| **$T_{12}$:$T_{51}$** | 0.52 | Q = 43.59 ($p < 0.00001$) |
| **$T_{24}$:$T_{30}$** | 0.08 | Q = 6.49 ($p = 0.00003$) |
| **$T_{24}$:$T_{51}$** | 0.25 | Q = 21.25 ($p < 0.00001$) |
| **$T_{30}$:$T_{51}$** | 0.18 | Q = 14.76 ($p < 0.00001$) |
| Figure S17 (spike area) | | |
| **Pairwise Comparisons** | **$HSD_{0.05}$ = 1.0964**<br>**$HSD_{0.01}$ = 1.3314** | **$Q_{0.05}$ = 3.6482**<br>**$Q_{0.01}$ = 4.4303** |
| **$T_{12}$:$T_{24}$** | 3.90 | Q = 12.98 ($p < 0.00001$) |
| **$T_{12}$:$T_{30}$** | 5.94 | Q = 19.78 ($p < 0.00001$) |
| **$T_{12}$:$T_{51}$** | 9.88 | Q = 32.89 ($p < 0.00001$) |

| | | |
|---|---|---|
| **$T_{24}$:$T_{30}$** | 2.04 | Q = 6.80 ($p$ = 0.00001) |
| **$T_{24}$:$T_{51}$** | 5.98 | Q = 19.91 ($p$ < 0.00001) |
| **$T_{30}$:$T_{51}$** | 3.94 | Q = 13.11 ($p$ < 0.00001) |
